# Chromatin remodeling with combined FACT and BET inhibition disrupts oncogenic transcription in Diffuse Midline Glioma

**DOI:** 10.1101/2024.06.06.597703

**Authors:** Holly Holliday, Aaminah Khan, Anahid Ehteda, Hieu Nguyen, Samuel E. Ross, Nisitha Jayatilleke, Anjana Gopalakrishnan, Eyden Wang, Yolanda Colino Sanguino, Daisy Kavanagh, Xinyi Guo, Jie Liu, David Lawrence, Claire X. Sun, Rebecca Lehmann, Chi Kin Ip, Alvin Lee, Laura Rangel-Sanchez, Wenyan Li, Robert Salomon, Ron Firestein, Robert J Weatheritt, Fatima Valdes-Mora, Marcel E. Dinger, Timothy N. Phoenix, Chelsea Mayoh, Benjamin S. Rayner, Maria Tsoli, David S. Ziegler

## Abstract

Aberrant epigenetic regulation is a hallmark of Diffuse Midline Glioma (DMG), an incurable pediatric brain tumor. The H3K27M driver histone mutation leads to transcriptional dysregulation, indicating that targeting the epigenome and transcription may be key therapeutic strategies against this highly aggressive cancer. One such target is the Facilitates Chromatin Transcription (FACT) histone chaperone. We found FACT to be enriched at developmental gene promoters, coinciding with regions of open chromatin and binding motifs of core DMG regulatory transcription factors. Furthermore, FACT interacted and co-localized with the Bromodomain and Extra-Terminal Domain (BET) protein BRD4 at promoters and enhancers, suggesting functional cooperation between FACT and BRD4 in DMG. *In vitro*, a combinatorial therapeutic approach using the FACT inhibitor CBL0137, coupled with BET inhibition revealed potent and synergistic cytotoxicity across a range of DMG cultures. These results were recapitulated *in vivo*, significantly extending survival in three independent orthotopic PDX models of DMG. Mechanistically, we show that CBL0137 treatment decreased chromatin accessibility, synergizing with BET inhibition to cause broad transcriptional collapse, silencing several key oncogenes including *MYC, PDGFRA*, *MDM4* and *SOX2*, as well as causing alterations to the splicing landscape. Notably, this combination also elicited immune-related effects, including activation of the interferon response and antigen presentation mechanisms in DMG cells and induction of an activated state in macrophages and T cells, as demonstrated in an immunocompetent setting with spatial transcriptomics. Altogether, our data highlights the therapeutic promise of simultaneously targeting FACT and BET proteins in DMG, offering a dual tumor-intrinsic and immune-mediated strategy for combating this devastating pediatric brain tumor.

## Introduction

Diffuse midline glioma (DMG) is a highly aggressive brain tumor predominantly diagnosed in children under 10 years old. Recently classified as ‘DMG K27-altered’ by the 2021 World Health Organization, this designation recognizes the characteristic epigenetic dysregulation that arises from either the recurrent lysine-to-methionine driver mutations in Histone H3.1 and H3.3 genes (H3K27M), or overexpression of the oncohistone mimic EZHIP [1–7]. Both H3K27M and EZHIP inhibit Polycomb Repressive Complex 2 (PRC2) activity, leading to a global loss of Histone H3K27 trimethylation (H3K27me3), increased H3K27 acetylation (H3K27ac) and DNA hypomethylation. These changes establish an aberrant epigenetic landscape permissive to heightened transcriptional activity [8–14].

While direct therapeutic targeting of H3K27M has not been possible, targeting of epigenetic and transcription regulators has shown promise in several pre-clinical models of DMG (reviewed in [15, 16]). Among these regulators, the histone chaperone Facilitates Chromatin Transcription (FACT) comprising of two protein subunits; SSRP1 and SPT16, has emerged as a compelling target [17–19]. FACT plays a pivotal role in maintaining chromatin integrity by facilitating nucleosome turnover during transcription, DNA replication and repair. This chaperone is abundantly expressed in cancer cells, making it an attractive therapeutic target [20, 21].

The curaxin compound CBL0137 indirectly inhibits FACT by intercalating with DNA and trapping FACT on chromatin (a process known as ‘c-trapping’), effectively depleting the functional pool of this important chaperone [22–24]. We and others have demonstrated CBL0137 to be active against multiple tumor types, including neuroblastoma, glioblastoma, medulloblastoma, and diffuse intrinsic pontine glioma (DIPG)/DMG [19, 25–29]. This growing body of preclinical work has led to the advancement of CBL0137 to a Phase I/II clinical trial for pediatric patients with solid tumors, including DMG (NCT04870944). However, the aggressive nature of DMG necessitates combination therapies to address therapeutic resistance and reduce effective therapeutic doses. While initial studies combining CBL0137 with the pan-HDAC inhibitor panobinostat has shown promise against DMG pre-clinically [19], concerns over panobinostat’s high toxicity, and limited efficacy, led to its withdrawal from clinical studies and highlights the need for alternative combination strategies [30].

Disruption of transcription has proven effective against many ‘transcription addicted’ malignancies, often achieved by targeting Bromodomain and Extra-Terminal Domain (BET) proteins such as BRD4 [31]. BRD4 recruits the transcriptional machinery to hyper-acetylated chromatin at super-enhancers [32, 33], making it an attractive target against DMG, as demonstrated in the preclinical setting ([34–39] and reviewed in [40]). The tool compound JQ1, among other BET inhibitors, has demonstrated pronounced efficacy against DMG, through transcriptional disruption leading to cytostatis [35]. While next-generation BET inhibitors, including trotabresib (CC-90010), have entered clinical trials for brain tumors (NCT04047303, NCT04324840, NCT03936465)[40], their modest efficacy as single agents underscores the need for improved BET-targeting compounds and rational combination strategies.

Here we describe the promising efficacy of FACT and BET inhibition as a dual epigenetic therapy in cell line and animal models of DMG. Mechanistically, this epigenetic intervention induces chromatin compaction at gene promoters, and leads to widespread transcriptional collapse. Notably, oncogenes such as *MYC, PDGFRA, MDM4* and *SOX2* are directly regulated by FACT and BRD4, being silenced through dual inhibition. Additionally, we observe aberrant splicing, upregulation of interferon genes and the antigen presentation machinery, and changes in immune cell transcriptional signatures and phenotypes. Together, our findings provide proof-of-principle evidence for combining FACT and BET inhibition to co-target key oncogenic drivers and otherwise ‘untargetable’ pathways in DMG, and modulating the tumor-immune landscape in DMG for effective tumor clearance. While further optimization of BET inhibitors will be critical for clinical translation, these data establish a promising foundation for future therapeutic development.

## Results

### FACT and BRD4 are overexpressed, interact, and co-localize at active regulatory elements in DMG

To elucidate the genes bound and potentially regulated by FACT, we mapped the genome-wide binding sites of the FACT subunit SPT16 in H3.3K27M-mutant HSJD-DIPG007 and SU-DIPGVI, as well as H3-WT VUMC-DIPG10 cells using Cleavage Under Targets and Release Using Nuclease (CUT&RUN). We identified 4,259 peaks in HSJD-DIPG007 cells, 1,792 peaks in SU-DIPGVI cells, and 707 peaks in VUMC-DIPG10 cells (**Table S1A-B).** SPT16 peaks were over-enriched at proximal gene promoters in all cell lines (**Fig 1A**). SPT16-bound genes were enriched for mesenchymal differentiation across all lines, with transcription factor binding and GTPase signaling uniquely enriched in the histone mutant cells (**Fig 1B**). We focused subsequent analyses on the H3K27M lines cell lines, due to fewer peaks with weaker enrichment in VUMC-DIPG10 cells (**Fig S1A**). E-box motifs and motifs for EMT transcription factors were enriched at SPT16 binding sites, including key DMG regulators TCF12 and OLIG2 [41–43], and the invasive glioma drivers SNAI1/2 [44, 45] (**Fig 1C**). These findings suggest that FACT promotes transcriptional programs driven by lineage-defining and EMT transcription factors in DMG.

**Figure 1.**
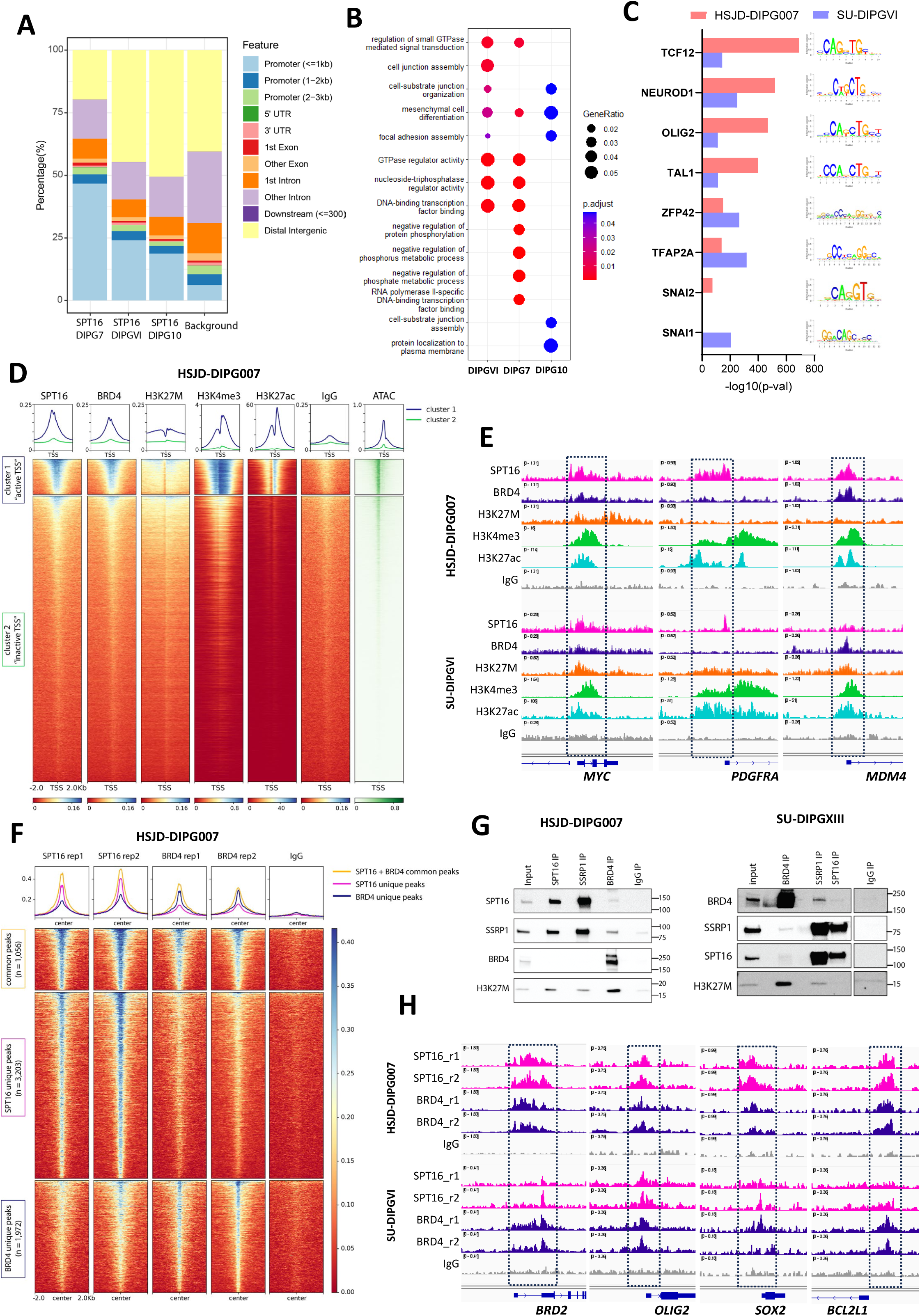
FACT and BRD4 expression and co-localization in DMG. **A)** Genomic distribution of SPT16 peaks in HSJD-DIPG007 (DIPG7), SU-DIPGVI (DIPGVI), and VUMC-DIPG10 (DIPG10) compared to a background of ∼1000 randomly shuffled genomic regions. **B)** Over-represented gene ontologies for genes associated with SPT16 peaks. **C)** Transcription factor motif enrichment analysis and motif logos for SPT16 peaks in HSJD-DIPG007 and SU-DIPGVI cells. **D)** Heatmaps depicting SPT16, BRD4, H3K27M, H3K4me3, IgG CUT&RUN signal and ATAC-seq signal at transcription start sites (TSS) in HSJD-DIPG007 neurospheres. Public H3K27ac ChIP-seq data was downloaded from [41]. Heatmaps are ranked based on SPT16 signal and separated into two k-means clusters. Average signal is shown in the profile plots. Representative data from n=2 experiments. **E)** Gene tracks display SPT16, BRD4, H3K27M, H3K4me3, IgG CUT&RUN and H3K27ac[41] ChIP-seq signal at TSSs of cancer-related genes *MYC, PDGFRA* and *MDM4* in HSJD-DIPG007 and SU-DIPGVI cells. **F)** Profile plots and heatmaps depicting the average SPT16, BRD4 and IgG CUT&RUN at SPT16 + BRD4 common peaks (orange), SPT16 unique peaks (pink), BRD4 unique peaks (blue) in HSJD-DIPG007 cells. **G)** Co-Immunoprecipitation and western blotting in HSJD-DIPG007 and SU-DIPGXIII nuclear extracts showing interactions between BRD4 with the FACT subunits and H3K27M. **H)** Gene tracks displaying SPT16 and BRD4 CUT&RUN signal at the TSSs of *BRD2, OLIG2, SOX2,* and *BCL2L1.* See related Figure S1.

We examined SPT16 occupancy at promoters genome wide, and assessed chromatin accessibility at these regions using Assay for Transposase-Accessible Chromatin using sequencing (ATAC-seq). The FACT complex is known to interact with H3K27M [19, 46]. Given that H3K27M and H3K27ac are known to incorporate into heterotypic H3K27M-H3K27ac nucleosomes, bound by acetyl-binding BET proteins at enhancer elements [36, 47], we investigated FACT colocalization with these nucleosomes. Transcription start sites (TSSs) were sorted by SPT16 enrichment and divided into two clusters which corresponded to highly accessible TSSs and less accessible TSSs. SPT16, BRD4 and H3K27M were enriched at accessible TSSs, which were also marked by the active histone modifications H3K4me3 and H3K27ac (**Fig 1D and Fig S1B**). Examples of SPT16 binding to the TSS of three DMG oncogenes (*MYC, PDGFRA,* and *MDM4*) are shown (**Fig 1E**).

To explore SPT16 and BRD4 co-occupancy, we visualized their enrichment at peaks classified as either unique to each protein, or common sites bound by both proteins. As expected, both SPT16 and BRD4 showed robust enrichment at the common peaks. Notably, each protein was also enriched at peaks unique to the other when compared to the IgG negative control, indicating co-localization of FACT and BRD4 at shared chromatin regions (**Fig 1F and S1C**). To confirm the physical interaction between these proteins, we performed endogenous co-immunoprecipitation of the FACT subunits, BRD4, and H3K27M from nuclear extracts of three H3K27M cell lines (HSJD-DIPG007, SU-DIPGVI and SU-DIPGXIII). BRD4 co-immunoprecipitated the FACT subunits consistent with previous reports in other cell types [48–50] **(Fig 1G and Fig S1D)**. Furthermore, FACT and BRD4 were found to interact with H3K27M, consistent with previous studies [19, 36, 46] **(Fig 1G and Fig S1D).** Notably, FACT was only able to co-precipitate BRD4 and H3K27M in two of three cell lines tested. This discrepancy may be due to transient and possibly indirect interactions between FACT, BRD4, and H3K27M, making it challenging to reproducibly capture, especially when examining endogenous protein levels under native conditions. Genes bound by SPT16 and BRD4 were involved in transcription, chromatin remodeling and p53 signaling (**Fig S1E**). Several example genes co-bound by FACT and BRD4 are shown at genes encoding the BET protein *BRD2,* OPC lineage transcription factors *OLIG2 and SOX2*, and survival gene *BCL2L1* (**Fig 1H**).

Since DMG is heavily driven by epigenetic and transcription dysregulation, we hypothesized that FACT and BET proteins, shown to be upregulated in other cancers [19, 20, 51–54], would also be elevated DMG. Consistently, *BRD4, SSRP1* and *SUPT16H*, were significantly overexpressed in DMG patients from the ZERO Childhood Cancer Program compared to normal brain tissue [55, 56] (**Fig S1F**). However, their expression was not higher in DMG than in other pediatric brain tumors, suggesting a broad role for these proteins in diverse malignancies. These genes were also upregulated in H3K27M-DMG cell lines compared to non-malignant cells, but not higher than in other pediatric cancer lines [57] (**Fig S1G**). Furthermore, SSRP1 and BRD4 proteins were upregulated in a panel of DMG cell lines compared to normal astrocytes (**Fig S1H**). Finally, whole genome CRISPR screening revealed that *BRD4*, *SSRP1*, and *SUPT16H* are essential for cell viability (gene effect < -0.5) of HGG cell lines from the Childhood Cancer Model Atlas, regardless of their histone mutant status (**Fig S1I**). In summary, FACT and BRD4 are overexpressed, interact, and co-localize at transcriptionally active chromatin, in the vicinity of key genes required for growth and development. This prompted us to test the combination of FACT and BET inhibition as a therapeutic strategy.

### Combined FACT and BET inhibition synergistically decrease DMG cell proliferation

Treatment with either of the BET inhibitors JQ1 or trotabresib resulted in decreased proliferation of primary DMG cells, with 72h IC_50_ values ranging from 1-50μM and 0.6-7μM, respectively, across a range of DMG cultures (**Fig 2A-B and Table S2-3**). Further testing on isogenic cell lines, with or without the H3K27M mutation, revealed significantly increased sensitivity to single-agent CBL0137, JQ1, and trotabresib in H3K27M cells compared to H3K27M KO cells. Notably, significant differences were observed in both IC_50_ and area under the curve (AUC) values with different H3K27M mutational status, with a 5-fold increase in sensitivity observed in H3K27M mutant compared to H3K27-KO DMG cells for CBL0137 and JQ1, and a 4-fold increase for trotabresib treatment (**Fig 2C, Fig S2A and Table S4**). These findings suggest that H3K27M-driven chromatin dysregulation increases dependence on FACT and BRD4.

**Figure 2.**
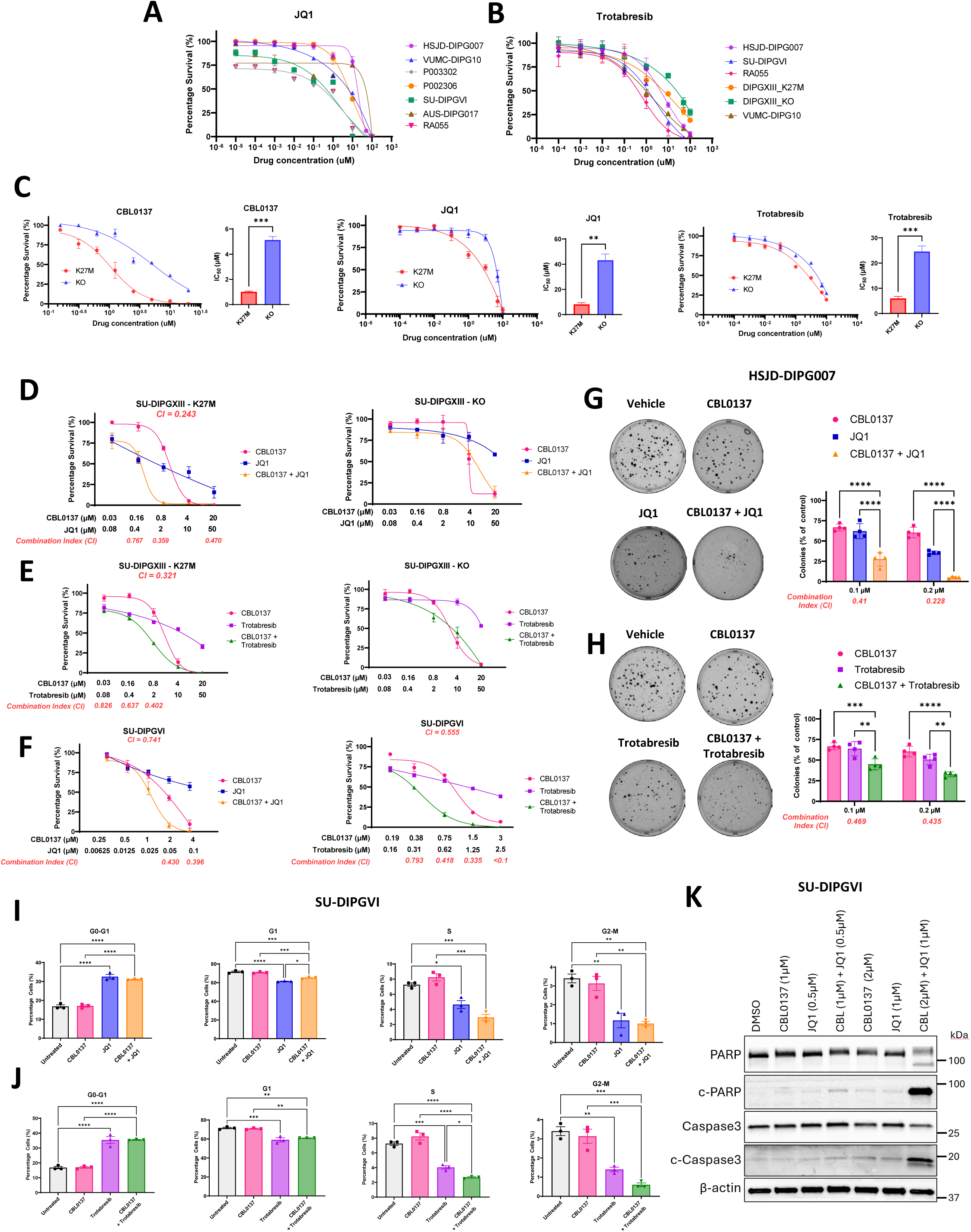
In vitro therapeutic efficacy of combinatorial FACT and BRD4 inhibition. **A-B)** Dose-response curves representing cell viability in a panel of patient-derived DMG cell lines treated with JQ1 **(A)** and trotabresib **(B)** following 72 hours. **C)** Dose response curves in isogenic SU-DIPGXIII cells (+/- H3K27M) and bar plots depicting IC50 values for CBL0137, JQ1 and trotabresib. **D-E**) Cell viability curves in SU-DIPGXIII isogenic cells treated with a combination of CBL0137 and JQ1 (**D**) or trotabresib (**E**). (**F**) Dose-response curves for SU-DIPGVI cells treated with CBL0137 combined with JQ1 and trotabresib. All dose response curves are represented as percentage viability compared to untreated controls, mean ± SEM (n = 3). **G-H**) Colony formation assays in HSJD-DIPG007 cells treated with the indicated drugs for 2 weeks. Data is presented as mean ± SEM (n=4). Representative images shown from the 0.2µM concentration. (**I-J**) Flow cytometry cell cycle distributions in SU-DIPGVI cells treated with CBL0137 combined with JQ1 (**I**) or trotabresib (**J**) for 48 hours. (n=3). (**K**) Western blot analysis of apoptosis markers in SU-DIPGVI cells treated with CBL0137 and JQ1 for 24 hours. Representative blot shown (n=3). P-values were calculated using a two-tail t-test (**C**) or an ANOVA test with Tukey’s multiple comparisons test for single and combination treatments (**G-J**). * p<0.05, **p<0.01, ***p<0.001, ****p<0.0001. Combination index (CI) values were calculated using Calcusyn, with synergistic CI values indicated. See related Figure S2.

Next, we assessed the effect of combined inhibition of both FACT and BRD4 in the isogenic histone-mutant and WT DMG cells. We found potent efficacy of the combination of CBL0137 and either BET inhibitor (JQ1 or trotabresib) in SU-DIPGXIII H3K27M mutant cell lines, and this synergy was lost upon deletion of the H3K27M mutation (**Fig 2D-E**). A similar synergistic effect between CBL0137 and the BET inhibitors was observed in four additional histone-mutant, patient-derived DMG cultures (H3.3 K27M: SU-DIPGVI, HSJD-DIPG007, RA055; H3.1 K27M: P005401) (**Fig 2F and Fig S2B-D**). However, we also found the combination to be synergistic against histone wild-type (WT) HGG/DMG cultures (P016802, VUMC-DIPG10, and AUS-DIPG17) (**Fig S2E-G).** These findings suggest that while H3K27M directly contributes to dependency on FACT and BRD4, alternative oncogenic drivers in histone WT tumors, such as MYCN amplification or CDKN2A/B loss, can create similar vulnerabilities to chromatin-targeting therapies (**Table S5).**

The combinations also led to synergistic inhibition of colony formation, at concentrations of each compound ranging between 100-200nM for HSJD-DIPG007 and RA055, and as low as 25-50nM for SU-DIPGVI (**Fig 2G-H and Fig S2H**). Lower drug concentrations were used in clonogenic assays to model sustained, sub-lethal exposure over time, in contrast to higher doses in short-term viability assays. Analysis of cell cycle distribution using flow cytometry revealed that combined FACT and BET inhibition induced G0-G1 arrest, reducing the proportion of cells in S, G2, and M phases, an effect driven by BET inhibition (**Fig. 2I-J, Fig. S2J-K**). Additionally, Western blotting showed increased levels of cleaved PARP and cleaved Caspase-3 following combination treatment, indicative of apoptotic cell death (**Fig. 2K**). Together, these data highlight the potential of combined FACT and BRD4 targeted inhibition as an effective strategy in DMG.

### Combined FACT and BET inhibition causes chromatin condensation at gene promoters in DMG

Given that CBL0137 and BET inhibitors target epigenetic and transcriptional regulators [17–19, 58], we next sought to determine the effect of these drugs on chromatin accessibility and transcription. To measure chromatin accessibility, ATAC-seq was performed in two H3.3 K27M DMG cell lines (HSJD-DIPG007 and SU-DIPGVI) treated with either CBL0137 or JQ1 as monotherapies or in combination after 4 hours to examine the early and direct changes to chromatin at doses previously used in other mechanistic studies [20, 29, 35, 59, 60]. Combinatorial treatment of DMG with CBL0137 and JQ1 led to a decrease in ATAC-seq signal at TSSs, consistently observed across independent replicates from both cell lines tested (**Fig S3A-B**). This effect was more driven by CBL0137, with Principal Component Analysis (PCA) revealing that the CBL0137 treated samples clustered closely with combination treated samples (**Fig S3C-D**).

We next examined chromatin accessibility at SPT16 and BRD4 CUT&RUN peaks, finding that both drugs alone, and in combination, decreased chromatin accessibility at these genomic regions (**Fig 3A and Fig S3E-F**), with the combination significantly decreasing chromatin accessibility at FACT and BRD4 binding sites more than each single agent in SU-DIPGVI cells, an effect more strongly driven by CBL0137 (**Fig 3A, Fig S3F**). This suggests that FACT and BRD4 are important for maintaining open chromatin, and that inhibition of these factors with CBL0137 and JQ1 leads to chromatin closure/condensation at these sites. Furthermore, differential accessibility analysis revealed that combination treatment resulted in ∼15-fold more closed chromatin regions compared to open regions in both cell lines tested (HSJD-DIPG007: 11,704 regions closed vs. 828 regions opened; SU-DIPGVI: 41,457 regions closed vs. 2,620 regions opened; FDR < 0.05, |LFC| > 1) (**Fig 3B, Fig S3G-H**). There was a significant degree of overlap in the regions closed by the combination treatment in both the models **(Fig 3C)** (hypergeometric p-value = 0.008). These regions were enriched at proximal gene promoters (**Fig 3D**) with their associated genes involved in pathways critical for DMG tumorigenesis such as development, proliferation, and histone modification (**Table S1C** and **Fig 3E**). For example, combination treatment led to chromatin condensation at the promotor of the FACT-bound oncogenes *MYC*, *PDGFRA*, and *MDM4* (**Fig 3F).**

**Figure 3.**
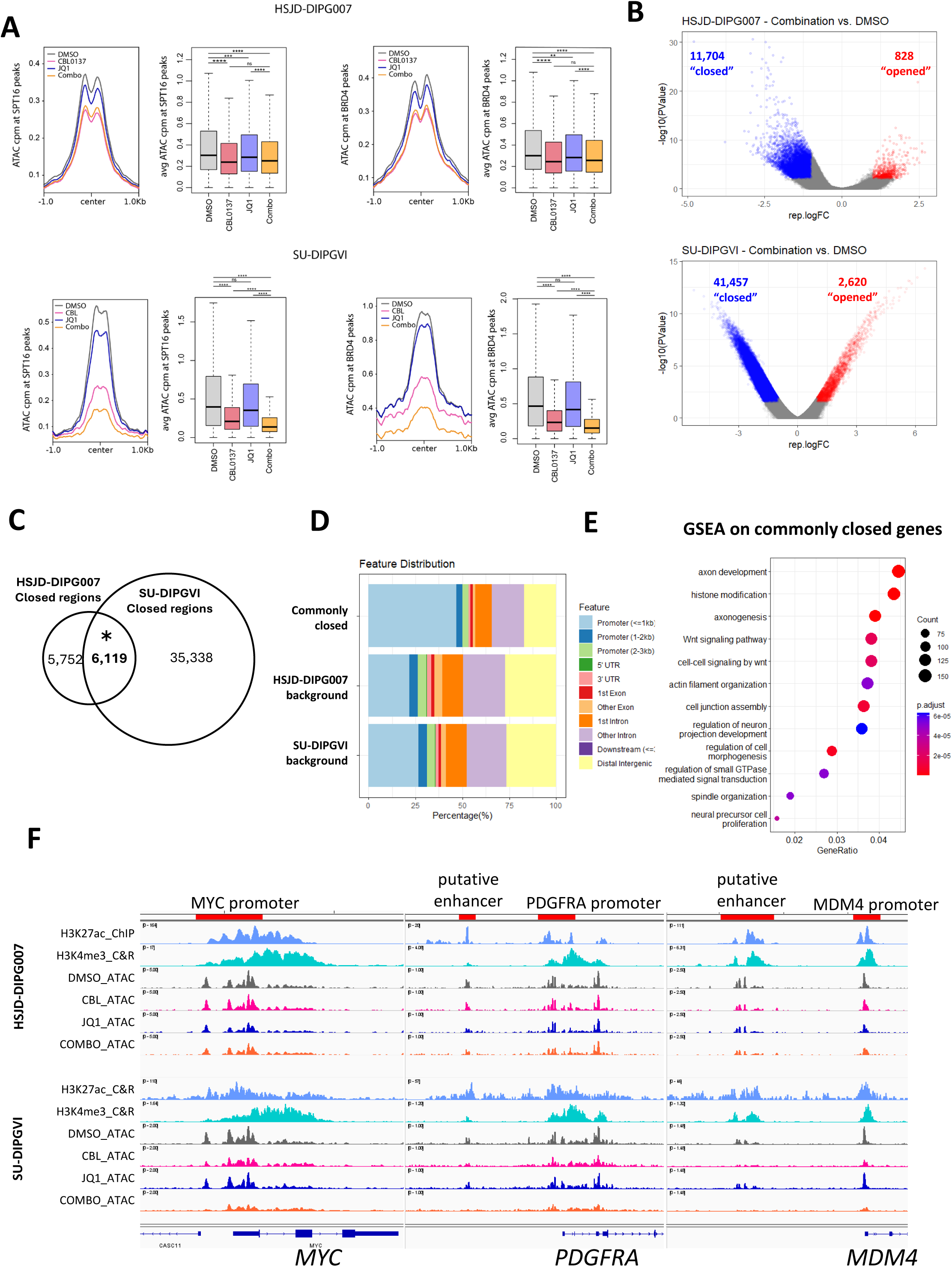
CBL0137 and JQ1 condense chromatin at gene promoters. **A)** Profile and box plots displaying ATAC-seq signal at SPT16 peaks (left) and BRD4 peaks (right) in DMG cells treated with DMSO, CBL0137 (1µM for HSJD-DIPG007 and 2µM for SU-DIPGVI), JQ1 (1µM) or the combination for 4 hours. A Wilcox test was used to test significance between treatment groups. **p<0.01, ***p<0.001, ****p<0.0001. Representative data from n=2 experiments for HSJD-DIPG007 and n=3 experiments for SU-DIPGVI with remaining replicates shown in Fig S3E-F. **B)** Volcano plot showing differentially accessible regions. Blue data points represent significantly (FDR < 0.05, LFC < -1) downregulated “closed” regions, red data points represent significantly upregulated “opened” (FDR < 0.05, LFC > 1) regions, with the number of regions indicated. **C)** Venn diagram displaying the overlap between the closed regions common to both cell lines. A hypergeometric test was used to determine over enrichment. **D)** Genomic distribution of the overlapping closed regions from panel C compared to background ATAC-seq genomic windows. **E**) Dot plot of over-represented gene ontologies in the genes associated with the overlap regions from panel C. **F)** Gene tracks display ATAC-seq signal at promoters and putative enhancers of MYC, PDGFRA, and MDM4. H3K27ac (from [41] [47]) and H3K4me3 (this study) enrichment is also shown. Representative replicate from n=2-3 experiments. See also Fig S3.

The H3K27M oncohistone hinders the spreading of H3K27me3 from PRC2-landing sites, resulting in a decrease in H3K27me3 genome wide [2–4, 11]. We have previously shown that CBL0137 increases global H3K27me3 levels and that this is potentiated by the HDACi panobinostat [19]. Consistently, in this study, we observed an increase in global H3K27me3 levels induced by CBL0137 through western blotting (**Fig S3I-J**), and at specific loci using CUT&RUN (p-val < 0.05) (**Fig S3K**). However, this increase was not further enhanced by JQ1 (**Fig S3I-J**), suggesting that the mechanism by which the CBL0137 interacts with JQ1 is distinct from its mechanism of synergy with panobinostat [19].

### Combined CBL0137 and JQ1 treatment leads to broad transcriptional silencing

To assess the impact of chromatin condensation caused by CBL0137 and JQ1 treatment on transcription, we performed nascent RNA-seq using TimeLapse-seq [61]. Unlike conventional RNA-seq, where the effects of transcriptional inhibition can be masked by stable pre-existing transcripts, TimeLapse-seq enables the specific identification of newly synthesized mRNAs directly regulated by treatment [61]. Notably, the impact of each drug on nascent transcripts was far more extensive than on total RNA (**Fig 4A, Fig S4A and Table S1D**). Combination treatment had the most widespread effect compared to each monotherapy, with over 60% of all detected genes being significantly (FDR<0.05, LFC< -1) downregulated (9,159/13,153 for HSJD-DIPG007 and 8,930/13,919 for SU-DIPGVI), indicating that CBL0137 and JQ1 induce a broad suppression of transcription (**Fig 4A and Fig S4A**).

**Figure 4.**
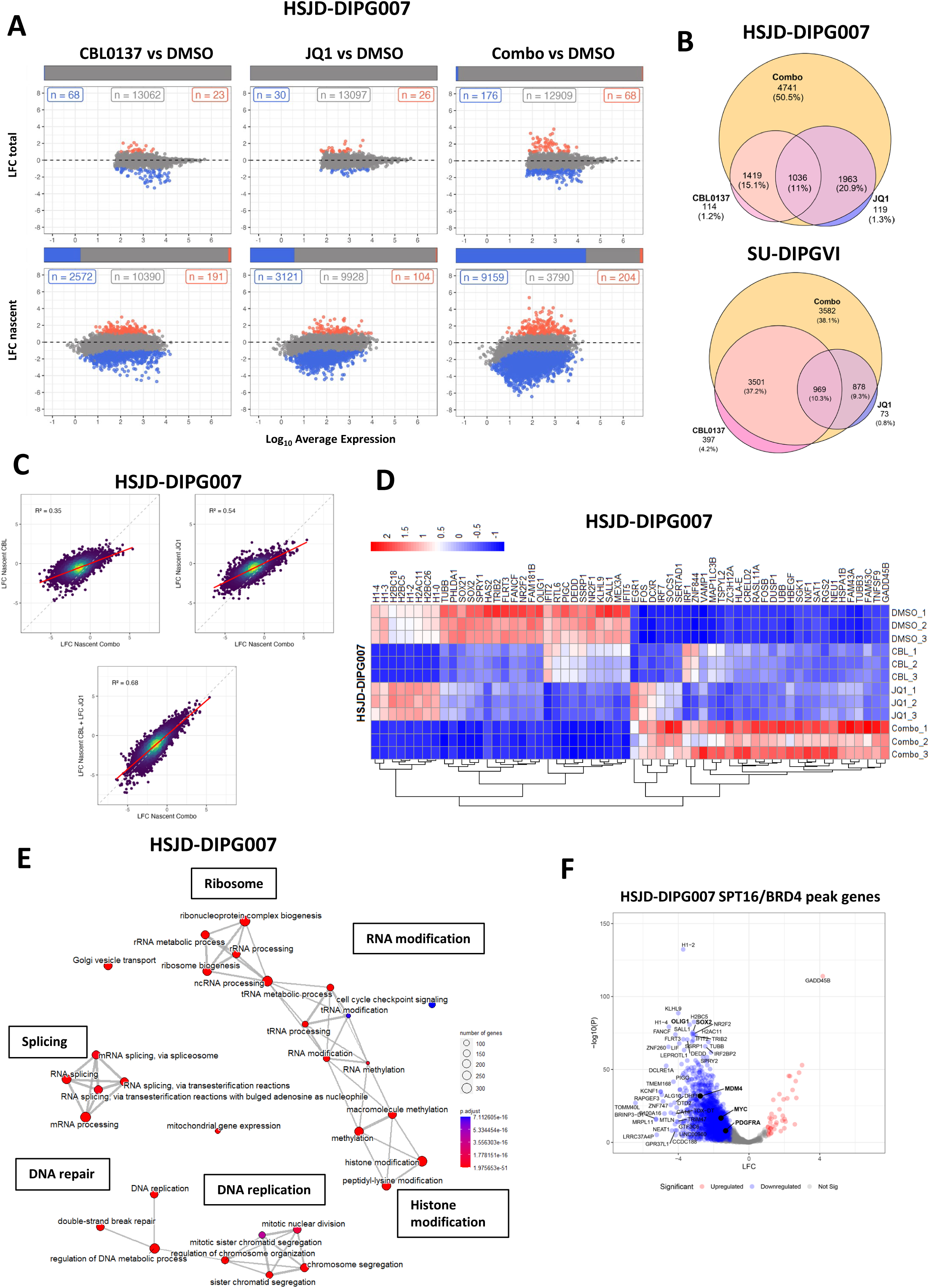
CBL0137 and JQ1 induce a broad suppression of transcription. **A)** Total RNA (upper) and nascent RNA (lower) MA plots showing significantly differentially expressed genes in HSJD-DIPG007 cells treated with CBL0137 (1µM), JQ1 (1µM), or the combination. Blue points represent significantly (FDR < 0.05, LFC < -1) downregulated genes, red points represent significantly upregulated genes (FDR < 0.05, LFC >1) with the number of genes indicated. Bars indicate the proportion of DE genes. **B)** Proportional Venn diagrams illustrating the overlap of downregulated nascently transcribed genes in cells treated with single agents and combination therapy. **C**) Correlations of log fold-change in nascently transcribed genes between combination therapy, each monotherapy, and the sum of both monotherapies in HSJD-DIPG007 cells. **D**) Nascent RNA-seq heatmap displaying top 30 up and downregulated genes DMSO vs Combination treated HSJD-DIPG007 cells. **E)** Enrichment map of over-represented gene ontologies among nascently transcribed genes downregulated by combination treatment in HSJD-DIPG007 cells, with high-level processes highlighted in labelled boxes. **F**) Volcano plot showing differential expression of nascently transcribed genes associated with SPT16 and BRD4 peaks in HSJD-DIPG007 cells. Key genes are labelled. See also Fig S4.

To investigate the nature of the combination response we overlapped the genes downregulated by each treatment compared to DMSO and found that a large subset (38-50%) of repressed genes were unique to the combination treatment, indicating that the combined action of CBL0137 and JQ1 are required to silence these targets (**Fig 4B**). To determine if the combination effect was additive or synergistic, we compared the fold-change in gene expression between the combination and single agents. If the combination were synergistic in silencing gene expression, the magnitude of fold-change in the combination treatment would exceed the sum of the fold-changes observed with each single agent. However, although the combination strongly correlated with the additive effect (R² = 0.68– 0.77; **Fig 4C, Fig S4B**), it did not surpass it. This suggests that, while the combination suppresses more genes overall, the effect on individual genes is additive rather than synergistic.

Examining individual genes revealed notable top upregulated genes induced by the combination, related to stress (*GADD45B, EGR1, UBB*), neuronal differentiation (*TUBB3*), and antigen presentation and interferon response (*HLA-E, IRF1/7, SOCS1*) (**Fig 4D, Fig S4C, Table S1D**) with GADD45B and TUBB3 protein induction confirmed by western blot (**Fig S4D**). Conversely, downregulated genes included several histone genes and oligodendrocyte progenitor cell transcription factors (*OLIG1, SOX1/2*) (**Fig 4D, Fig S4C, Table S1D**). Gene set enrichment analysis of the genes downregulated by combination treatment revealed several critical processes involved in cancer cell growth and proliferation such as DNA replication, DNA repair, histone modification, RNA modification, ribosome biogenesis, and splicing **(Fig 4E** and **Fig S4E**). Mass spectrometry of CBL0137-treated DMG cells showed concordant downregulation of proteins involved in translation and RNA processing (**Fig S4F** and **Table S1G**). Integration of CUT&RUN with nascent RNA-seq identified genes whose transcription is directly sustained by FACT and BRD4. Genes associated with FACT and/or BRD4 binding exhibited strong transcriptional repression following combination treatment (**Fig 4F, Fig S4G),** including critical DMG stem/progenitor regulators OLIG2 and SOX1/2.

### CBL0137 and JQ1 combination treatment represses transcription of key oncogenes

Among the directly downregulated targets were key oncogenes, including *MYC, PDGFRA, MDM4,* and *SOX2* (**Fig 4F**, **Fig 5A**), which also displayed reduced chromatin accessibility at FACT and BRD4-bound promoters/enhancers (**Fig S5A**). Transcriptional and protein repression of these targets was validated in independent RNA-seq and western blot experiments (**Fig 5B-C, Fig S5B-C, Table S1E**). MYC is an established transcriptional target of both FACT and BRD4 [26, 58], PDGFRA is a known super-enhancer associated glioma oncogene [35, 36], SOX2 is a potent stem cell transcription factor [62], and MDM4 is a p53 negative regulator with additional oncogenic roles [63, 64]. Given MDM4’s role as a p53 inhibitor, we assessed p53 levels and observed CBL0137-mediated p53 protein upregulation selectively in p53-wildtype HSJD-DIPG007 cells, but not in p53-mutant SU-DIPGVI cells (**Fig S5E**).

**Figure 5.**
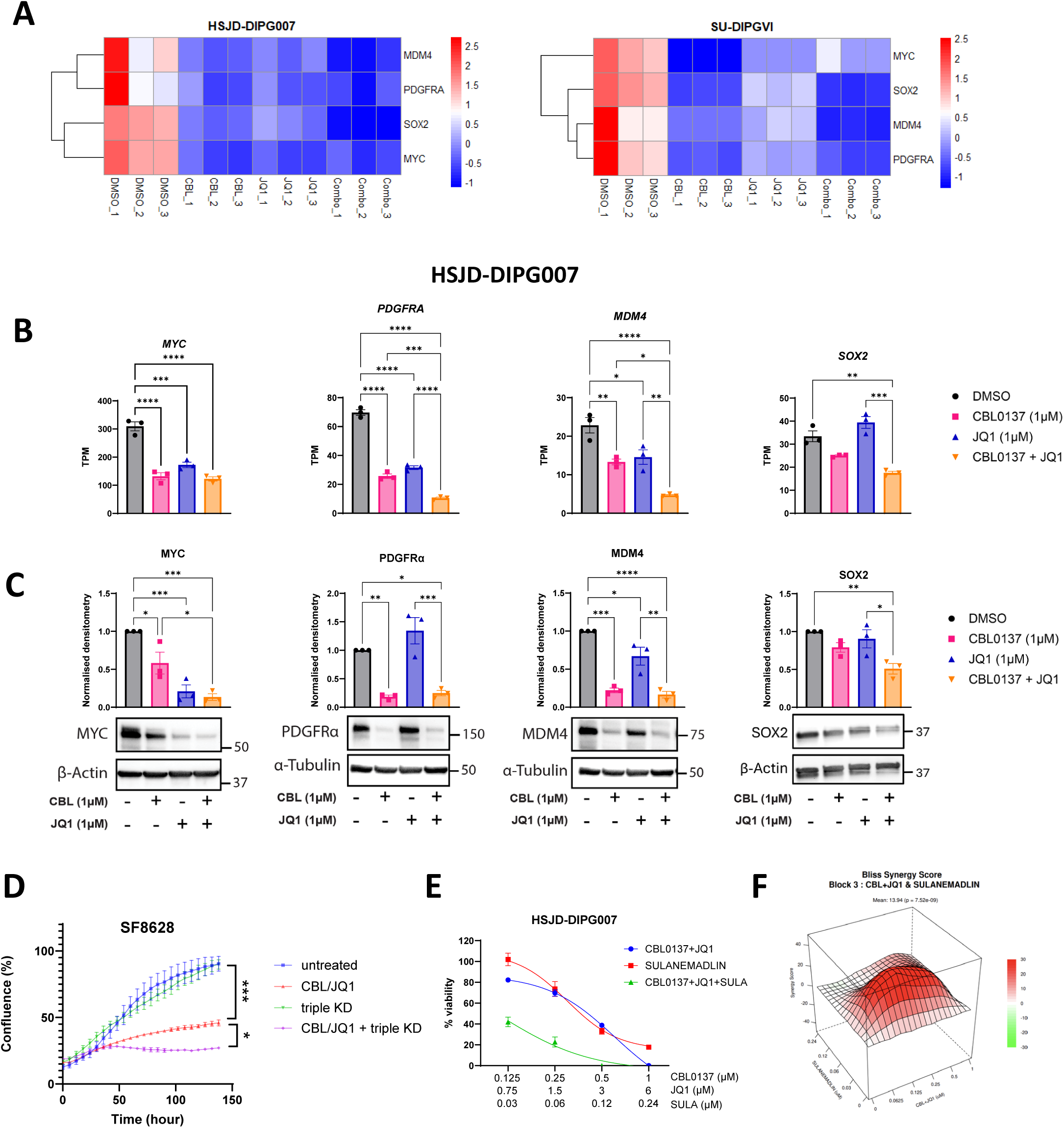
Chromatin compaction by CBL0137 and JQ1 decreases expression of oncogenes. **A)** Nascent RNA-seq heatmap displaying *MYC, MDM4, PDGFRA,* and *SOX2* genes in the indicated treatment groups for HSJD-DIPG007 and SU-DIPGVI cells. **B**) RNA-seq TPM values (mean +/- SEM; n=3) for *MYC, PDGFRA*, *MDM4* and *SOX2* in HSJD-DIPG007 cells treated for 4 h with CBL0137 and JQ1. **C**) Representative MYC, PDGFRα, MDM4 and SOX2 western blots and bar charts displaying densitometry (mean +/- SEM; n=3) normalized to β-Actin or α-Tubulin and expressed as a fold-change relative to the DMSO control. 24 h treatment. PDGFRα and MDM4 were run on the same gel, and use the same loading control. **D**) Proliferation of SF8628 cells measured by time-lapse microscopy every 6 h for 6 days. Cells were transfected with non-targeting siRNA or siRNA targeting MYC, MDM4 and PDGFRA (tripleKD) and treated with DMSO or the combination of CBL0137 (0.2µM) and JQ1 (0.5µM). **E**) Cytotoxicity assay comparing CBL0137+JQ1, sulanemadlin, and the combination of all 3 drugs in HSJD-DIPG007 cells after 96 h and Bliss synergy score matrix comparing CBL0137+JQ1 and sulanemadlin. Significance was calculated using a one-way ANOVA with Sidak’s multiple comparisons test for single and combination treatments. * p<0.05, **p<0.01, ***p<0.001, ****p<0.0001.

We next investigated whether depleting MYC, MDM4, and PDGFRA could further enhance the anti-proliferative action of the drug combination. To this end, we performed siRNA knockdowns (KD) for each gene individually and in combination (triple KD) in SF8628 H3K27M DMG cells, achieving 40–70% knockdown (**Fig. S5F**), which was comparable to the transcriptional repression observed with combination drug treatment (**Fig. S5G**). The synergy between CBL0137 and JQ1 was confirmed in these cells (**Fig. S5H**). Treatment with CBL0137 and JQ1 resulted in a significant reduction in cell proliferation, and this effect was potentiated with depletion of *MYC*, *MDM4*, and *PDGFRA* (**Fig. 5D**). We next evaluated a triple combination of CBL0137 and JQ1 with the clinically relevant MDM2/4 inhibitor sulanemadlin. As expected, sulanemadlin exhibited high potency in p53 wild-type cells, while isogenic HSJD-DIPG007 cells lacking p53 were more than 200-fold resistant [65] (**Fig. S5I**). Similarly, p53-mutant DMG cell lines (SU-DIPGVI and SU-DIPGXVII) also displayed resistance (**Fig. S5I**), confirming p53 restoration as the primary mechanism of action for sulanemadlin in DMG. Cytotoxicity assays demonstrated that sulanemadlin synergized with CBL0137 and JQ1 in both p53 wild-type HSJD-DIPG007 cells and in p53-mutant SU-DIPGVI cultures (**Fig. 5E-F, Fig. S5J**). These findings uncover a novel therapeutic combination involving MDM inhibition, identified through our investigation into the mechanisms of CBL0137 and JQ1.

### CBL0137 and JQ1 alter the splicing landscape in DMG

Since CBL0137 and JQ1 led to downregulation of splicing genes in our nascent RNA-seq experiments (**Fig 4E**, **Fig S4E, Fig S6A**), and since chromatin structure is known to influence splicing through a co-transcriptional process [66], we next examined alternative splicing in an independent RNA-seq experiment. Nascent RNA-seq was performed after 2 hours of treatment, and splicing was analyzed at 4 hours to allow time for splicing to occur. Single-agent treatment resulted in limited differential splicing. However, combination treatment led to hundreds of differential splicing events compared to control cells, with an average of 800 events in HSJD-DIPG007 cells and 2000 events in SU-DIPGVI cells. Some of these splicing events occurred within the same gene, affecting approximately 400 unique genes in HSJD-DIPG007 and 1000 unique genes in SU-DIPGVI (**Fig. S6B** and **Table S1F**). The most common form of alternative splicing was intron retention, accounting for 40-50% of events (**Fig S6C**). Examples of intron retention are shown at three loci, which also exhibited condensed chromatin in the vicinity of the retained introns (**Fig S6D**). These findings suggest that, in addition to disrupting chromatin dynamics and transcription, CBL0137 and JQ1 may also alter the splicing landscape in DMG.

### Dual inhibition of FACT and BRD4 enhances survival in orthotopic models of DMG

To evaluate the therapeutic efficacy of combined FACT and BET inhibition, we used three orthotopic xenograft models using distinct patient-derived DMG cells: SU-DIPGVI-LUC, HSJD-DIPG007, and RA055. The dosing schedule (CBL0137 every 4 days and JQ1 five days a week) was based on the drug’s pharmacokinetics, tolerability, and prior studies. CBL0137’s relatively longer half-life (∼6h) supports infrequent dosing, while JQ1’s shorter half-life (∼1h) necessitates more frequent administration [19, 36, 67–69].

In the SU-DIPGVI-LUC model, single-agent treatments with CBL0137 and JQ1 exhibited either negligible or limited anti-tumor effects, respectively. However, the combination regimen significantly extended survival compared to single-agent treatments and vehicle cohorts by delaying tumor growth, as seen by xenogen imaging (median survival: vehicle: 73 days, CBL0137: 76 days, JQ1: 78 days, CBL0137/JQ1: 95.5 days; **Fig 6A-B** and **Table S6**). The combination regimen was well tolerated, with no observed weight loss or overt hematological or biochemical toxicities over the four-week treatment period (**Fig S7A-C, Table S9**). We have previously demonstrated that CBL0137 can penetrate the blood–brain barrier (BBB) [19], and here show that JQ1 is likewise detectable in the brain, as measured by liquid chromatography–tandem mass spectrometry (LC–MS/MS) (**Fig S7D**). Immunohistochemical analysis of brain tissue collected following 4 weeks of combination treatment revealed a significant reduction in cell proliferation, as evidenced by decreased Ki67 staining, and a concurrent reduction in H3K27ac staining (**Fig 6C-D**). Extension in survival with combined FACT and BET inhibition was also evident in the HSJD-DIPG007 model, with the median survival of the combination-treated cohort extended to 93.5 days compared to 70.5 days in the vehicle group (**Fig 6E, Fig S7B** and **Table S7**).

**Figure 6.**
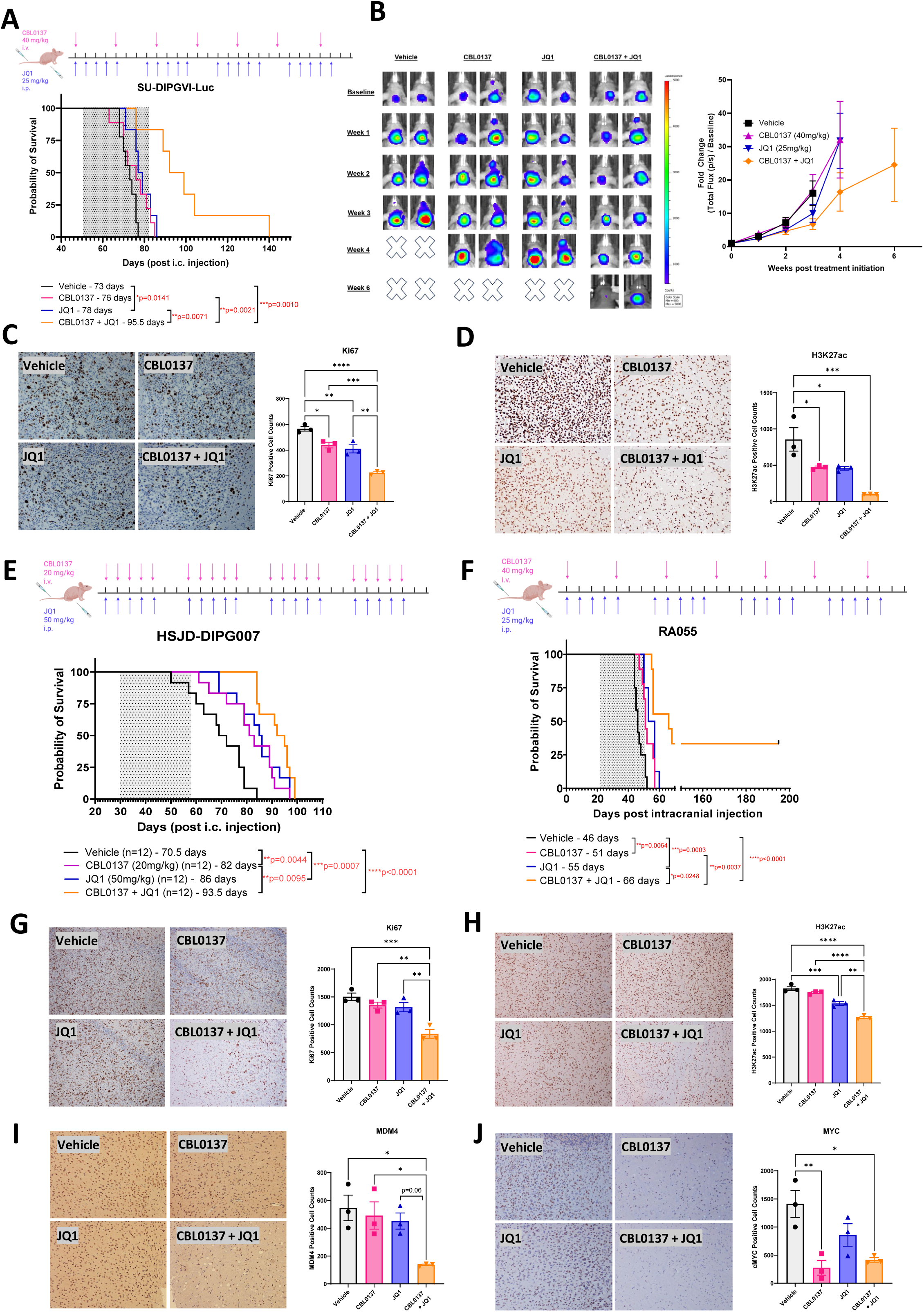
Therapeutic efficacy of CBL0137 and JQ1 combination treatment in orthotopic models of DMG. **A)** Treatment scheme and survival curve of SU-DIPGVI-Luciferase with CBL0137/JQ1 treatment. Median survival of cohorts (days): Vehicle (n=9) = 73, CBL0137 (n=9) = 76, JQ1 (n=6) = 78, CBL0137 + JQ1 (n=6) = 95.5. Exact p-values listed in Supplementary Table 6. **B)** Xenogen luminescence imaging of SU-DIPGVI-Luciferase tumor burden and averaged luminescence signal measured over time in each cohort. **C-D**) IHC staining and quantification for Ki67 (**C**) and H3K27ac (**D**) of SU-DIPGVI-LUC tumors after 4 weeks of treatment (n=3). **E**) treatment schema and survival curve for the HSJD-DIPG007 model with CBL0137/JQ1 treatment. Median survival of cohorts (days): Vehicle (n=12) = 70.5, CBL0137 (n=12) = 82, JQ1 (n=12) = 86, CBL0137 + JQ1 (n=12) = 93.5. Exact p-values listed in Supplementary Table 7. **F**) Treatment scheme and survival curve of RA055 model with CBL0137/JQ1 treatment. Median survival of cohorts (days): Vehicle (n=12) = 46, CBL0137 (n=9) = 51, JQ1 (n=8) = 55, CBL0137 + JQ1 (n=9) = 66. Exact p-values listed in Supplementary Table 8. **F-J**) Immunohistochemistry staining of RA055 tumors for Ki67 (**G**), H3K27ac (**H**), MDM4 (**I**), and MYC (**J**). Three images were taken from brain samples collected from three mice in each cohort at the end of the treatment period. Scale: 50µm. For survival curves, statistical analysis was performed using the Log-Rank (Mantel-Cox) with multiple test corrections applied. For IHC analysis data is presented as mean values ± SEM. *p<0.05, **p<0.01, ***p<0.001, ****p<0.0001. p-values were calculated using one-way ANOVA with Tukey’s multiple comparisons test for treated and untreated cohorts. See also Fig S7.

These promising results were recapitulated in the highly aggressive RA055 model, characterized by a median survival of 46 days in untreated mice post intracranial injection. Treatment with the combination of CBL0137 and JQ1 significantly enhanced survival, with approximately one-third of treated mice surviving until the experimental endpoint of 201 days (**Fig 6F**, **Fig S7D** and **Table S8**). Consistent with the SU-DIPGVI-LUC model, Ki67 cell proliferation and H3K27ac immunohistochemistry staining were significantly reduced with combination treatment compared to single-agent and untreated tissue samples (**Fig 6G-H**). We also observed a significant reduction in MDM4 and MYC positive cells in PDX tumors treated with the combination, validating our *in vitro* mechanistic findings in an independent *in vivo* model (**Fig 6I-J**). Taken together, this combination strategy is effective against multiple aggressive patient-derived DMG models. Furthermore, the combination of CBL0137 with the clinically-relevant BET inhibitor ZEN-3694 [70, 71] was also well tolerated *in vivo*, with no evidence of weight loss or toxicity (**Fig S7F-G, Table S10**), and pharmacokinetic analysis showed that ZEN-3694 reaches the brain at levels comparable to plasma (**Fig. S7H**). These results support the feasibility of translating combined FACT and BET inhibition into a clinically actionable strategy for the treatment of DMG.

### CBL0137 and JQ1 influence immune-associated pathways in DMG and tumor-infiltrating immune cells

Interferon signaling plays a key role in anti-tumor immunity and involves upregulation of the antigen presentation machinery (reviewed in [72, 73]), and epigenetic therapies can enhance expression of the antigen presentation machinery in other cancers [64][74–77]. Similarly, in DMG cells we observed that combination treatment upregulated transcription of interferon response genes (*IRF1, IRF3, IRF7, SOCS1*) and MHC-I antigen presentation genes (*HLA-A, HLA-B, HLA-C, HLA-E*) (**Fig 7A and Fig S8A**). as well as positive enrichment of the ‘antigen processing and presentation of exogenous peptide antigen’ pathway (**Fig 7B**). To complement our acute RNA-seq study, we examined expression changes following an extended 48-hour treatment using qPCR. These results confirmed that CBL0137 elevates expression of interferon and antigen presentation genes (*IRF1, HLA-A, IFNA, IFNB,* and *OAS1*) (**Fig S8B**). Finally, flow cytometry confirmed increased in HLA-ABC and HLA-E surface expression in response to combination treatment (**Fig 7C and Fig S8C**).

**Figure 7.**
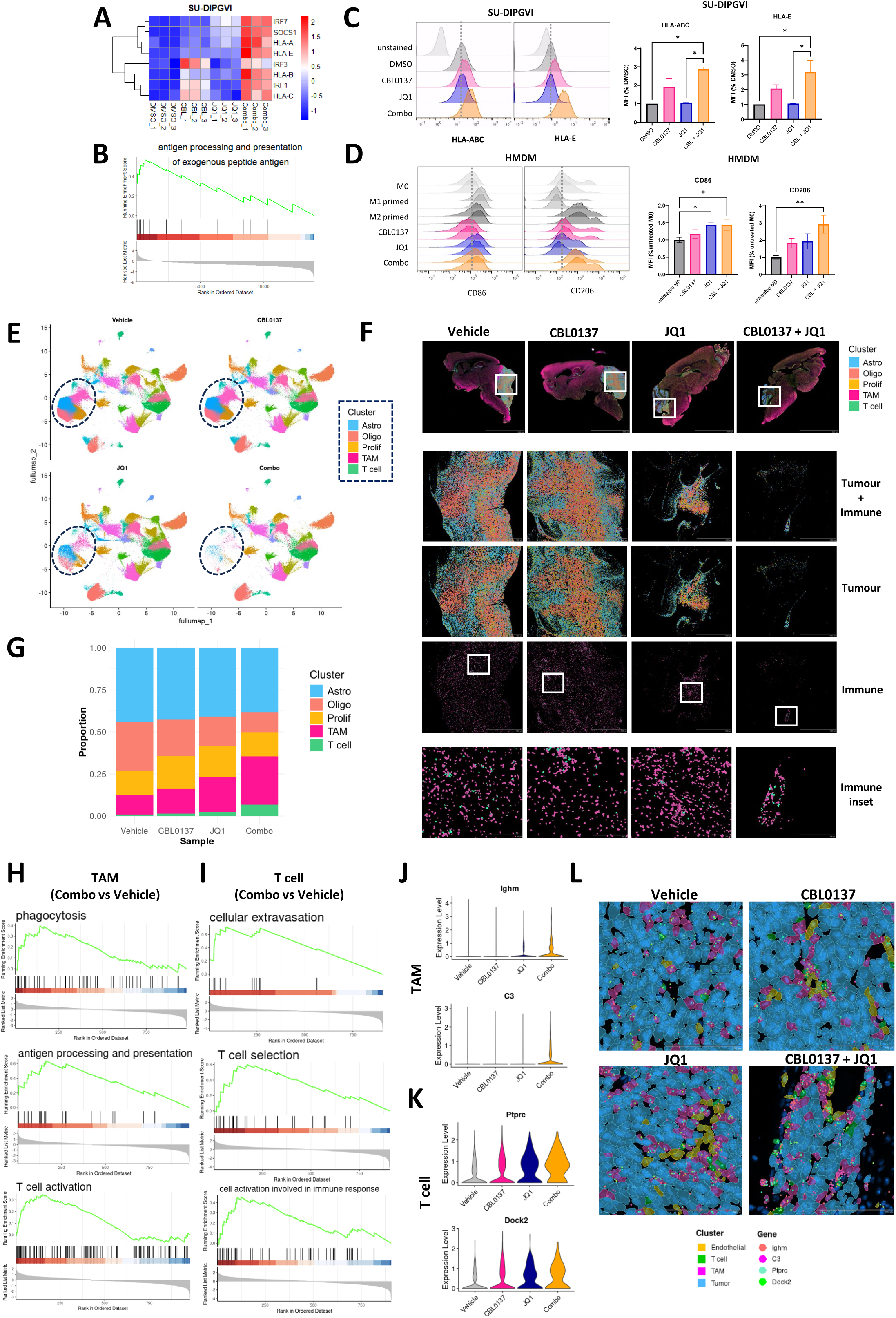
CBL0137 and JQ1 influence immune-associated pathways in DMG and tumor-infiltrating immune cells. **A)** Nascent RNA-seq heatmap displaying interferon and antigen presentation genes in the indicated treatment groups for SU-DIPGVI cells. **B**) GSEA enrichment plot showing positive enrichment (upregulation) of the antigen presentation pathway in SU-DIPGVI cells (DMSO vs Combination). **C)** Flow cytometry histograms for HLA-ABC and HLA-E in treated SU-DIPGVI cells (24h). Representative data from at least 3 independent experiments. Median Fluorescence Intensity (MFI) values are shown. **D**) Flow cytometry histograms for CD86 and CD206 in human monocyte-derived macrophages (HMDM) cells in M1 or M2 polarization conditions or treated with CBL0137 and JQ1 for 48h. Representative data from 3 independent experiments (3 donors) shown. MFI values were quantified as above. P-values for MFI comparisons **(C,D)** were calculated using a one-way ANOVA with Sidak’s multiple comparison test. data is presented as mean values ± SEM. *p<0.05, **p<0.01. **E**) Normalized Xenium single cell UMAP plot of immunocompetent murine glioma model treated with CBL0137 and JQ1 for 3 weeks. UMAP is split into the 4 treatments cohorts. Tumor-associated clusters are indicated in the dashed circles. **F**) Xenium Explorer images of immunocompetent model. Upper panels: whole brain sagittal slices with tumor cells (Astrocyte, oligodendrocyte lineage and proliferative subtypes) and immune cells (tumor associated macrophage; TAM, and T cells) overlaid onto the immunofluorescence image. Scale = 5000µM. Middle panels: Higher magnification insets of tumor region showing the distribution of tumor and immune clusters that are highlighted in panel E. Scale = 1000 µM. Lower panel: higher magnification insets displaying immune clusters only. Scale = 200µM. **G**) Composition of each tumor-associated cluster relative to the total of all tumor-associated clusters. **H-I**) GSEA enrichment plots comparing combination to vehicle (red = upregulated, blue = downregulated) in TAMs (**H**) and T cells (**I**). **J-K**) Violin plots for select genes significantly upregulated in the vehicle vs combination treated TAMs (*Ighm* and *C3*) (**J**) and T cells (*Ptprc* and *Dock2*) (**K**). **L**) Xenium Explorer images of tumor, immune and endothelial cells with individual transcripts represented by dots. Scale = 100µM. See also Fig S8.

Given the prominence of immunosuppressive macrophages in DMG [15], and the epigenetic regulation of their polarization [78], we next assessed whether CBL0137 and JQ1 modulate the inflammatory response in cultured human monocyte-derived macrophages (HMDMs).Treatment increased mRNA expression of pro-inflammatory M1 marker genes (*IRF3, IRF8, IFNB, HIF1A, p53 and IL1B)* [79] (**Fig S8D**). Using flow cytometry, we show that combination treatment increased surface expression of both the M1 marker CD86, a T cell co-stimulatory molecule, and the M2 marker CD206, implicated in phagocytosis and antigen cross-presentation [80] (**Fig 7D**). Moreover, combination-treated HMDMs displayed increased phagocytic activity in a fluorescent bead uptake assay **(Fig S8E**).

We investigated the effects of CBL0137 and JQ1 in an *in vivo* immunocompetent model. C57Bl6 mice were orthotopically engrafted with syngeneic H3.3 K27M glioma cells as previously described [73, 74], and tumors were allowed to establish. Following three weeks of treatment, we performed *in situ* spatial transcriptomic profiling using the Xenium Prime 5000-gene panel. Single cells were clustered and annotated based on marker gene expression and spatial localization (**Fig. S8F–H, Table S1H**), with a focus on tumor cells and infiltrating immune cells. We identified three distinct tumor populations characterized by elevated expression of astrocyte lineage markers (cluster 3: *Gfap, Sox9*), oligodendrocyte lineage markers (cluster 9: *Sox10, Pdgfra, Mag*), and proliferation-associated genes (cluster 11: *Mki67, Top2a, Plk1*) (**Fig. S8H-I, Table S1I).** Among CD45+ (*Ptprc*+) immune cells, we identified tumor-associated macrophages (TAMs; cluster 6: *Cd68, Itgam, Cx3cr1*) and infiltrating T cells (cluster 24: *Il2rb, Cd3d/g, Cd4, Cd8a/b1, Gzma/b*) (**Fig. S8H, Table S1H).**

Combination treatment depleted both tumor and tumor-associated immune cell populations (**Fig. 7E–F**) and altered the cellular composition within the tumor microenvironment (**Fig. 7G**). Notably, there was a decrease in the tumor-to-immune cell ratio, as well as a relative increase in T cell abundance within the tumor following combination therapy **(Fig. 7G)**. To investigate transcriptional changes, we subclustered each cell type and performed differential gene expression analysis comparing vehicle- and combination-treated cells. In the tumor compartment (comprising all three tumor clusters), we observed downregulation of chromatin associated genes, transcriptional factors, and stem cell maintenance genes (**Fig. S8J).** Leading edge genes driving enrichment included *Myc* and *Sox2* (**Fig. S8K, Table S1J),** consistent with our *in vitro* findings which showed that FACT and BRD4 inhibition epigenetically silence these critical oncogenic drivers. Furthermore, the top upregulated gene was the interferon stimulated gene *Ifi27* (**Fig. S8K, Table S1J),** further supporting the induction of an interferon response in tumor cells by combination therapy.

In TAMs, there was positive enrichment of genes involved in phagocytosis (including *Ighm* and *C3*), antigen presentation, and T cell activation – hinting at tumor cell engulfment and antigen cross-presentation to T cells [81–83] (**Fig 7H, J, Table S1J**). Furthermore, T cells displayed transcriptional signatures consistent with infiltration and activation, including pathways related to cellular extravasation, T cell selection and immune activation. This included *Ptprc* and *Dock2* (**Fig 7I, K, Table S1J**). CD45, encoded by *Ptprc*, plays a critical role in T cell activation by initiating T cell receptor signaling, and DOCK2 is involved in formation of T cell immune synapses [84, 85]. Finally, immune cells were localized within close proximity to endothelial cells, together with their transcriptional profiles, this supports the notion of extravasation from blood vessels into the tumor. Visualization of individual transcripts for phagocytic and T cell activation genes (*Ighm, C3, Ptprc* and *Dock2*) confirmed increased expression within TAMs and T cells upon combination treatment (**Fig. 7L**).

Taken together, combination treatment exerts coordinated effects on both glioma and immune compartments – suppressing transcriptional stem cell programs and enhancing antigen presentation and interferon signaling in tumor cells, while simultaneous reprogramming TAMs towards a more phagocytic and immunostimulatory state and promoting T cell activation.

## Discussion

Despite hundreds of clinical trials, survival outcomes for DMG patients have not changed in over three decades [15]. The current standard-of-care, radiotherapy, only delays tumor progression and the majority of children still survive less than one year from diagnosis. Since the discovery of the H3K27M mutation in 2012, efforts to target multiple layers of epigenetic regulation, including DNA methylation [12], histone modification [86–89], chromatin readers [35, 36], chromatin remodelers [90–92], and transcription machinery [35, 93], have been ongoing. Given DMG’s aggressive, heterogeneous nature and multiple undruggable oncogenic pathways, monotherapies are unlikely to be effective. Here we have uncovered a novel mechanism-anchored epigenetic combination therapy - FACT and BET inhibition - which simultaneously disrupts multiple oncogenic pathways and demonstrates potent pre-clinical activity against DMG.

FACT disassembles and reassembles nucleosomes during cellular processes that require DNA access, such as during transcription or replication. This overcomes the physical hindrance of nucleosome-compacted DNA, allowing RNA Pol II or the replication machinery to travel through chromatin unimpeded [22, 23]. Our findings indicate that FACT maintains open chromatin at promoters and enhancers, supporting transcription driven by developmental TFs (TCF12, OLIG2, SNAI1) in DMG. This is consistent with a recent study showing that FACT preserves open chromatin at regulatory elements to promote transcription by core pluripotency factors in embryonic stem cells [94]. BRD4 binds to acetylated chromatin via its tandem bromodomains and recruits positive transcription elongation factor (P-TEFb) to active enhancers, releasing paused RNA pol II, leading to transcriptional initiation and elongation [95]. Combined targeting of FACT and BRD4 led to chromatin compaction and widespread transcriptional silencing, including several notable oncogenes. For example, *MYC and PDGFR*α, are both super enhancer-activated oncogenes involved in gliomagenesis [12, 96]. Another downstream target was *MDM4*, a negative regulator of p53, suggesting this treatment could potentially restore p53 function in p53-WT DMGs [63]. Thus, this drug intervention simultaneously disarms multiple epigenetic-driven oncogenes in DMG. However, it is unlikely that the cytotoxic effects arise solely from silencing several genes. Rather, we propose that the global disruption of transcriptional output – a fundamental stressor in rapidly dividing cells – is the key driver of cell death.

While off-target effects cannot be fully excluded, the consistent phenotypic and transcriptomic responses across multiple models suggest that the therapeutic effects are primarily driven by on-target inhibition of transcriptional regulation. Model-specific differences in transcriptional response and drug sensitivity may reflect intrinsic features such as p53 status, with HSJD-DIPG007 (p53 WT) showing more JQ1-driven changes and SU-DIPGVI (p53 mutant) more CBL0137-driven effects.

The combination of CBL0137 with either of the BET inhibitors tested had higher efficacy against H3K27M cells compared to isogenic cells lacking H3K27M. This could be explained by two non-mutually exclusive hypotheses: i) Direct inhibition of H3K27M function by targeting co-factors like FACT and BRD4, or ii) increased dependency on these factors due to downstream epigenetic reprogramming. The first hypothesis is supported by prior studies and our data showing chromatin localization and co-immunoprecipitation of FACT and BRD4 with H3K27M. An IP-mass spectrometry study previously identified SPT16 and SSRP1 as H3K27M interactors in mitotic DMG extracts [46], and additional studies have reported interactions between H3K27M and both FACT and BRD4 [19, 36, 46]. Alternatively, the hypomethylated chromatin creates a dependency on FACT and BRD4 to maintain open chromatin and high levels of transcription. Treatment with CBL0137 and JQ1 then leads to chromatin compaction at transcription start sites, leading to transcriptional collapse. This mechanism mirrors recent studies targeting the SWI/SNF chromatin remodeling complex, where the epigenetic reprogramming characteristic of DMG creates a dependency on this complex to drive pathogenic gene expression [90–92]. Notably, CBL0137 and JQ1 were still effective against histone WT DMG, and both FACT and BRD4 emerged as critical dependencies in pHGG cell lines irrespective of H3 mutation status. The synergy of this combination has also been observed in other H3 WT pediatric brain tumors, namely medulloblastoma and Atypical Teratoid/Rhabdoid Tumor (AT/RT) cell lines [97], indicating potential broader therapeutic applications in pediatric brain cancers beyond H3K27M DMG.

We also uncovered a role for this combination treatment in alternative splicing, possibly through downregulation of the splicing machinery. As splicing is influenced by chromatin structure, we hypothesize that alternative splicing may also result from transcriptional stalling. Specifically, RNA pol II may encounter increased resistance when transcribing through compacted chromatin, leading to splicing errors [66], particularly in the context of reduced expression of splicing genes. This hypothesis warrants further exploration, for example by mapping RNA pol II elongation rates. 3D genome architecture is an emerging contributor to tumorigenesis in epigenetically driven cancers, including DMG and ependymoma [41, 98]. Both CBL0137 and BET inhibitors can disrupt topologically associated domains (TADs) and enhancer–promoter loops [41, 60]. Although not explored in this study, examining the impact of combination therapy on genome topology represents an exciting avenue for future investigation.

The *in vivo* efficacy of the combined FACT and BRD4 inhibition in multiple orthotopic models of DMG indicates its potential as a promising combinatorial treatment strategy. Employing an alternative dosing regimen involving an increased dose of CBL0137 (40 mg/kg) paired with a reduced dose of JQ1 (25 mg/kg) not only proved well-tolerated but also resulted in a significantly enhanced survival response in highly aggressive DMG models. This underscores the importance of exploring diverse dosing regimens for the development of combination therapies.

Epigenetic therapies have been shown to activate an immune response through various mechanisms [99]. For example, FACT, HDAC and DNMT inhibitors, have been shown to trigger an interferon response and enhance expression of the antigen presentation machinery in other cancers [64][74–77]. While BET inhibitors, such as JQ1, have been associated with suppression of interferon signaling due to downregulation of key interferon-stimulated genes in glioblastoma [100], we observed transcriptional activation of the interferon pathway in DMG cells treated with the combination of CBL0137 and JQ1. This suggests CBL0137 may counteract the interferon-suppressive effects of BET inhibition, possibly through induction of double-stranded RNA (dsRNA), which is a known trigger of the type I interferon response. Indeed, CBL0137 has been shown to induce dsRNA and activate antiviral pathways in various non-DMG cancer model [27, 101, 102]. This is particularly relevant given our observation of increased intron retention, which may generate neoantigens [103, 104]. When presented via MHC-I, these can enhance tumor immune visibility. Consistent with this, we detected increased MHC-I surface expression on DMG cells following treatment. Together, these findings support a model whereby chromatin disruption by CBL0137 induces splicing errors and/or dsRNA, leading to interferon activation and increased antigen presentation.

In macrophages, we observed upregulation of both M1- and M2-like markers, suggesting that the drug combination influences the plasticity of TAMs, which are known to adopt mixed or transitional states [105, 106]. In an immunocompetent mouse model, we observed decreased tumor burden and immune rewiring consistent with our *in vitro* findings. This included TAM reprogramming toward a phagocytic phenotype, as well as T cells exhibiting heightened infiltration and activation signatures. Altogether, our data indicates that this combination elicits both tumor-intrinsic and immune-mediated effects. By simultaneously disrupting oncogenic transcriptional programs and enhancing immune detection, CBL0137 and BET inhibition may offer a dual therapeutic strategy for DMG and other immunologically ‘cold’ pediatric brain tumors, particularly in combination with immunotherapies such as CAR T cells.

Our data nonetheless highlights the therapeutic potential of this combinatorial approach of BETi and FACTi. CBL0137 is currently under clinical investigation in pediatric DMG (PEPN2111), and our findings provide a strong mechanistic rationale for future combinations with transcriptional or chromatin-disrupting agents to enhance therapeutic efficacy. Together, this work lays a foundation for novel strategies to exploit the epigenetic dependencies of DMG and improve outcomes for children with this devastating disease.

## Materials and Methods

### Cell culture

DMG cells were maintained as neurospheres in Tumor Stem Cell (TSM) media consisting of a 1:1 mixture of DMEM/F12 and Neurobasal medium (Invitrogen) supplemented with Glutamax, Pyruvate, non-essential amino acids, HEPES buffer, antibiotic/antimycotic, B-27 (Invitrogen), Heparin (0.0002%), EGF (20 ng/mL), bFGF (20 ng/mL), PDGF-AA (10 ng/mL) and PDGF-BB (10 ng/mL) (StemCell Technologies). SU-DIPGXIII K27M and K27M-KO isogenic cells were maintained according to [12], adherently on flasks coated with poly-L-ornithine (0.01%) (Sigma) and laminin (10 µg/mL) (Sigma), in Neurocult NS-A proliferation media (StemCell Technologies) supplemented with heparin (0.0002%), EGF (20 ng/mL) and bFGF (10 ng/mL) (StemCell Technologies). SF8628 cells were maintained adherently in DMEM supplemented with 10% FBS and 2mM L-Glutamine. All lines were cultured at 37°C and 5% carbon dioxide. Cells tested negative for mycoplasma contamination and were confirmed to match their original tumors by STR profiling.

Human monocyte-derived macrophages (HMDMs) were a kind gift from Prof. Kerry-Anne Rye, Cardiometabolic Research Group, UNSW, following isolation and culture as previously described [107], maintained in RPMI + 10% human serum with the media being replaced every 3 days. In polarisation studies, HMDM were either primed for 48 h with 1 µg/mL LPS (Sigma) and 20 ng/mL recombinant human IFNγ (Sigma) for the M1 phenotype, 20 ng/mL recombinant human IL-4 (Sigma) and 20 ng/mL recombinant human IL-13 (Sigma) for the M2 phenotype.

### CUT&RUN

Neurospheres were collected by centrifugation at 800 rpm for 3 min and dissociated into single cells with 1 mL Accutase (Sigma) for 5-10 minutes with occasional pipetting, diluted with 5 mL TSM-B, and passed through a 40 µm cell strainer. Cells were centrifuged at 1,200 rpm for 5 min and resuspended in complete media. Single cell suspensions were counted using a haemocytometer and cell viability was confirmed to be over 95%. CUT&RUN was performed using the EpiCypher CUTANA CUT&RUN kit according to the manufacturer’s instructions. Briefly, 500,000 viable cells per reaction were washed and permeabilized with 0.01% Digitonin. Cells were incubated with 0.5 µg antibody, or a 1:50 dilution for SPT16 (**Table 1**), on a nutator overnight at 4°C. Following washing, pAG-MNase was added to the reactions and CaCl_2_ was used to activate antibody-bound MNase for targeted chromatin digestion and release of DNA fragments. The entire volume of purified DNA (ranging from 1-10 ng) was used as input for NGS library preparation, according to the EpiCypher CUTANA Library Prep Kit instructions. Library concentration was determined using a Qubit Fluorometer (HS dsDNA) (Thermo) and library size determined using a 4200 Tapestation system on a high sensitivity D1000 ScreenTape (Agilent). Resulting libraries were sequenced with 50-150bp paired-end reads, with a minimum of 7M reads per sample at Ramaciotti Centre for Genomics (UNSW Sydney, Australia). Two biological CUT&RUN replicates were performed.

**Table 1.**
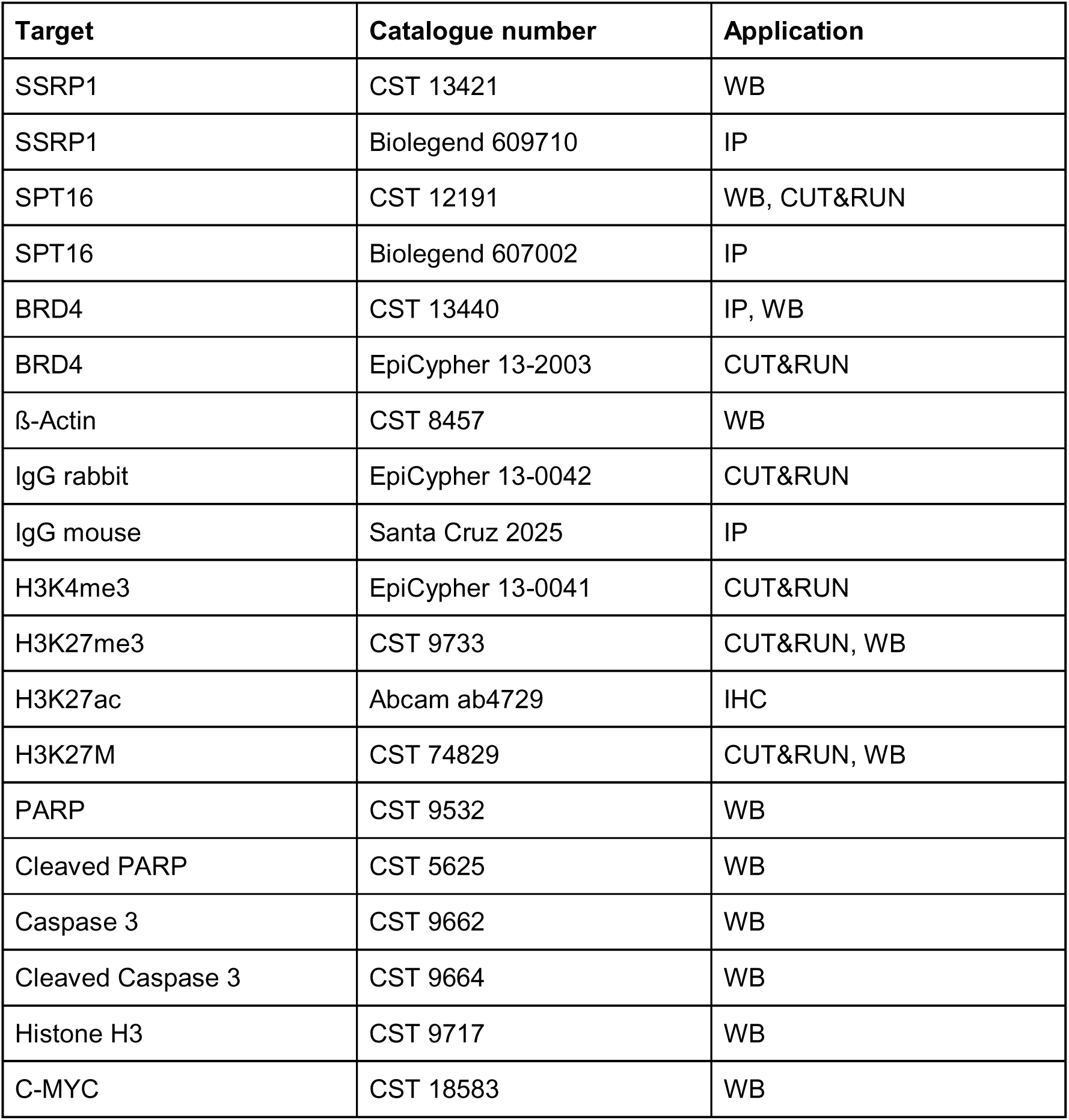
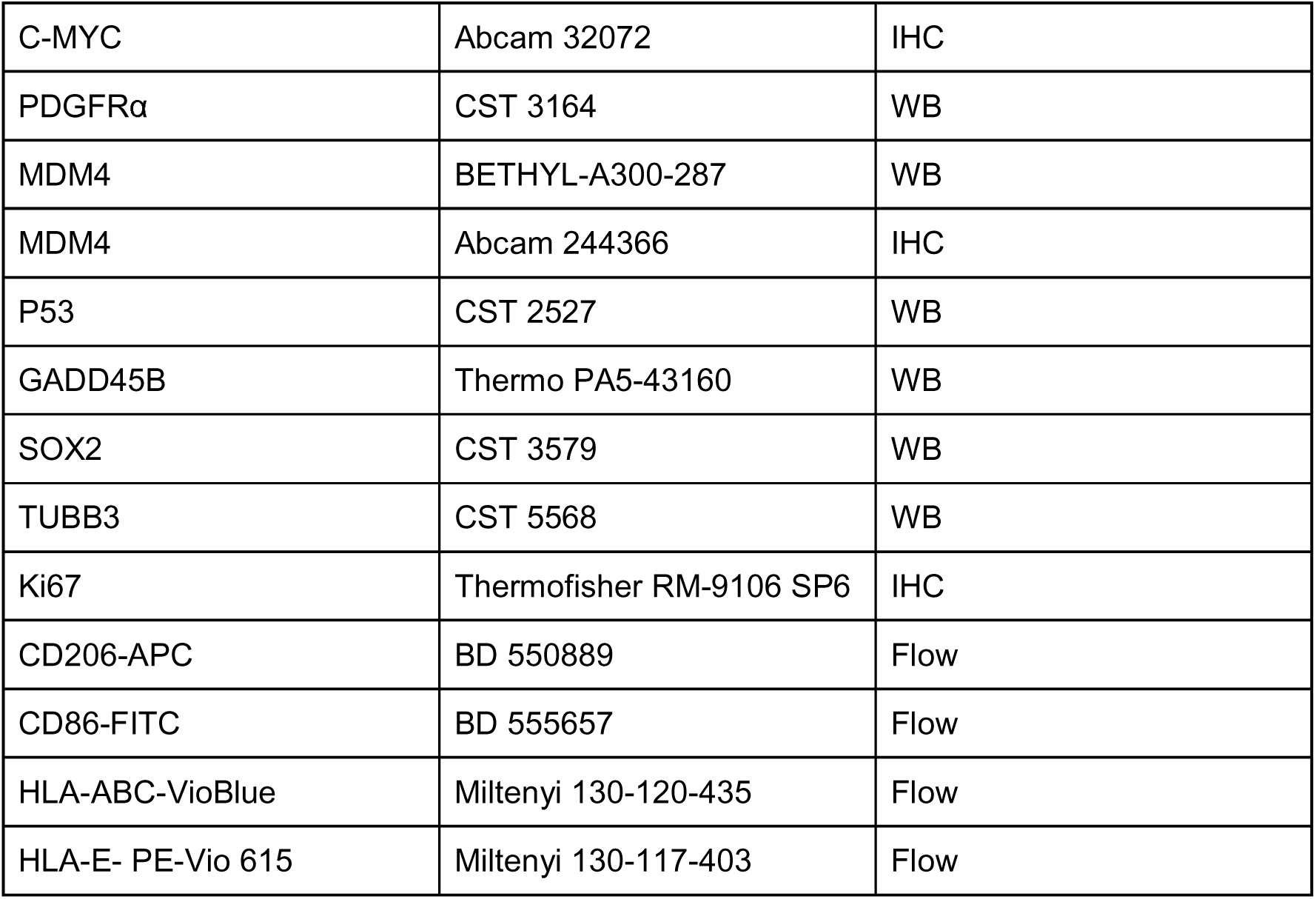
List of antibodies.

| Target | Catalogue number | Application |
| --- | --- | --- |
| SSRP1 | CST 13421 | WB |
| SSRP1 | Biologend 609710 | IP |
| SPT16 | CST 12191 | WB, CUT&RUN |
| SPT16 | Biologend 607002 | IP |
| BRD4 | CST 13440 | IP, WB |
| BRD4 | EpiCypher 13-2003 | CUT&RUN |
| β-Actin | CST 8457 | WB |
| IgG rabbit | EpiCypher 13-0042 | CUT&RUN |
| IgG mouse | Santa Cruz 2025 | IP |
| H3K4me3 | EpiCypher 13-0041 | CUT&RUN |
| H3K27me3 | CST 9733 | CUT&RUN, WB |
| H3K27ac | Abcam ab4729 | IHC |
| H3K27M | CST 74829 | CUT&RUN, WB |
| PARP | CST 9532 | WB |
| Cleaved PARP | CST 5625 | WB |
| Caspase 3 | CST 9662 | WB |
| Cleaved Caspase 3 | CST 9664 | WB |
| Histone H3 | CST 9717 | WB |
| C-MYC | CST 18583 | WB |
| C-MYC | Abcam 32072 | IHC |
| PDGFRα | CST 3164 | WB |
| MDM4 | BETHYL-A300-287 | WB |
| MDM4 | Abcam 244366 | IHC |
| P53 | CST 2527 | WB |
| GADD45B | Thermo PA5-43160 | WB |
| SOX2 | CST 3579 | WB |
| TUBB3 | CST 5568 | WB |
| Ki67 | Thermofisher RM-9106 SP6 | IHC |
| CD206-APC | BD 550889 | Flow |
| CD86-FITC | BD 555657 | Flow |
| HLA-ABC-VioBlue | Miltenyi 130-120-435 | Flow |
| HLA-E- PE-Vio 615 | Miltenyi 130-117-403 | Flow |

### CUT&RUN analysis

Quality control analysis of CUT&RUN reads was performed using fastQC (v0.11.9). Adapter trimming was performed using cutadapt (v4.2) with parameters: --error-rate 0.1 --times 1 --overlap 3 --action trim --minimum-length 17 --pair-filter any. Reads were aligned using bowtie2 (v2.5.0) with -k 1 against the GRCh38 human reference genome. BAM files were sorted, and optical duplicates marked using picard (v2.27.5) with parameters: --DUPLICATE_SCORING_STRATEGY SUM_OF_BASE_QUALITIES --OPTICAL_DUPLICATE_PIXEL_DISTANCE 100 --VALIDATION_STRINGENCY SILENT.

Marked duplicates were filtered and removed using samtools (v1.12). Peak calling was performed using MACS2 (v2.2.7.1) using IgG as the control. Narrow peaks parameters: --keep-dup all -p 1e-5 and broad peak using parameters: --keep-dup all -q 0.05 --broad --broad-cutoff 0.05. BigWig coverage files were generated using deeptools bamCoverage (v3.5.1) with parameters: --binSize 10 --normalizeUsing BPM --effectiveGenomeSize 2913022398. Broad peak settings were used for H3K27M and H3K27me3, and narrow peaks for SPT16, BRD4, H3K27ac, H3K4me3. Peaks from biological replicates were merged using BedTools [108].

BigWig coverage was visualized using DeepTools [109] at hg38 Ensembl genes with blacklisted CUT&RUN regions excluded [110], or at SPT16 and BRD4 MACS2 peaks. Integrated genomics viewer (IGV) [111] was used to visualize BigWig coverage at individual loci. Peaks were annotated using ChIPSeeker [112] and ClusterProfiler [113] was used for Gene Ontology over-representation analysis of the annotated genes. Transcription factor motif analysis was performed using the Cistrome SeqPos motif tool [114]. Differential binding analysis for H3K27me3 comparing between DMSO and CBL0137 treated cells was carried out using CSAW as described [115], filtering on genomic windows 5-fold higher than background, with a significance threshold of FDR < 0.05 (n=2).

### Co-immunoprecipitation (co-IP) assay

Co-IP was performed as described in [116]. Briefly, nuclei were isolated from 5 million cells with hypotonic buffer (20 mM Tris-HCl pH 7.4, 10 mM NaCl, 3 mM MgCl_2_, protease inhibitors (Roche)) for 15 min on ice, followed by addition of IGEPAL (0.5%; Sigma-Aldrich). Nuclei were collected by centrifugation (5 min, 1000 *x g*, 4°C) then lysed in non-denaturing lysis buffer (50 mM HEPES pH 7.4, 140 mM NaCl, 3 mM MgCl_2_, 1 mM EGTA, 1% Triton-X, 10% Glycerol, protease inhibitors (Roche)). The nuclear lysate was passed 6 times through a 25-G needle to shear the DNA, then incubated for 15 min on ice. Lysates were cleared by centrifuging at 13,000 rpm (top speed) for 15 min at 4°C to remove insoluble debris. The lysate supernatant was then incubated with 15 µL protein A/G magnetic beads (Pierce) pre-conjugated with 2.5 µg antibody (minimum 4 h, 4°C with rotation). Immunoprecipitation was carried out overnight at 4°C on a rotator. Magnetic beads were washed 3x with lysis buffer, and protein was eluted with 2X Laemmli Sample Buffer diluted in lysis buffer, boiled at 85°C for 10 min, followed by addition of DTT (100 mM; Sigma-Aldrich). Western blotting to detect immunoprecipitated proteins was performed as described below, except that VeriBlot (Abcam) was used for the secondary antibody at 1:1000.

### Western blotting

To extract total protein cells were incubated with RIPA buffer (CST) containing protease and phosphatase inhibitors (Roche) for 15 min on ice with occasional vortexing, followed by centrifugation at 13,000 rpm for 15 min at 4°C to remove debris. BioRad 4-20% proTEAN pre-cast gels were used for gel electrophoresis, and proteins were transferred for 1 h at 100 V onto 0.45 µm PVDF membranes for RIPA-soluble proteins, or 0.45 µm nitrocellulose membranes for histones. Membranes were blocked for 1 h in 5% milk diluted in Tris Buffered Saline with 0.01% Tween-20 (TBS-T). Western blotting was carried out overnight with the indicated antibodies (Table 1) diluted 5% Bovine Serum Albumin (BSA; Sigma-Aldrich) in TBS-T. Following washing, membranes were incubated in horseradish peroxidase-conjugated secondary antibody (CST) diluted in 5% skim milk/TBS-T (1:2000) for 1 hr at room temperature. Membranes were incubated with ECL substrate (Pierce) and bands imaged using a Bio-Rad ChemiDoc imaging system. Densitometry was carried out using Image Lab software, with local background subtraction, and normalization to the indicated housekeeping gene.

### Gene expression analysis from patient samples and cell lines

Patient expression data for *SSRP1, SUPT16H* and *BRD4* was obtained from the ZERO childhood cancer precision medicine program [55]. Normal brain expression data was obtained from McGill University [56]. Integrated RNA-seq expression data were batch corrected using ComBat-Seq from the Bioconductor package sva (v3.48.0). *SSRP1, SUPT16H* and *BRD4* expression in DMG and non-malignant cell lines was downloaded from the Childhood Cancer Model Atlas [57].

### CRISPR dependency in HGG cell lines

The CRISPR dependency scores for BRD4, SSRP1 and SUPT16H are obtained from Childhood Cancer Model Atlas (https://vicpcc.org.au/dashboard; CCMA [57]). The results are part of the whole-genome scale CRISPR screens for n=24 high grade glioma cell lines following the protocol described in [57, 117]. Data was obtained under the data transfer agreement with the CCMA team. Briefly, the primary cells were engineered with the Cas12 endonuclease and Blasticidin antibiotic resistance-expressing genes (Lenti-AsCpf1-Blast: 1xNLS-Cas12a lentiviral expression construct; Addgene #84750). The whole genome libraries used were Human CRISPR Knockout Humagne Set C and Set D Pooled Library (Addgene #172650 and #172651). Pooled knockout genetic screens were conducted over 28 and 35 days at a multiplicity of infection (MOI) of 0.3. The gene dependency score was determined using the chronos method [118], and scaling was performed across the whole CCMA cohort using the negative control genes and core essential genes (Achilles Common Essential Controls) from the depmap portal (https://depmap.org/portal; [119]).

### Cell Viability Assays

Cells were cultured in 96-well plates at a density of 2,500–3,000 cells per well. For adherent cells, drugs were added 24 hours after plating; for suspension cells, treatment was initiated 72 hours post-plating. After a 72-hour incubation period, cell proliferation was assessed using the resazurin assay (Sigma-Aldrich), with results expressed as a percentage relative to untreated controls. For combination treatments with CBL0137 (from Prof Andrei Gudkov and Prof Katerina Gurova) and BET inhibitors (JQ1 [Selleck S7110] and trotabresib [Synonym: CC-90010; MedChemExpress HY-137573]), cells were exposed to serial dilutions of each drug, and synergy was calculated using CalcuSyn, where CI values < 1 indicate synergy. For triple combination treatments with CBL0137, JQ1, and Sulanemadlin (Synonym: ALRN-6924; MedChemExpress HY-P4210), cells were treated for 96 hours in a matrix format, and BLISS synergy scores was assessed using SynergyFinder [120].

### Clonogenic Assays

Long-term clonogenic assays were conducted by plating 24-well plates with a base layer of 0.3 mL of 0.5% agar mixed with cell culture media. After 24 hours of incubation at 4°C, the base layer was overlaid with a 0.4 mL layer consisting of cells (500-1500 cells/well), 0.33% agar, cell culture media, and the drug at the indicated concentrations. The plates were then incubated in a 5% CO2 humidified atmosphere at 37°C. Following an incubation period of either 2 or 4 weeks, depending on the specific cell line, the colonies were stained with 3-(4,5-dimethylthiazol-2-yl)-2,5-diphenyltetrazolium bromide (MTT) solution (Sigma-Aldrich) at 5 mg/mL. After 1 hour, images of the colonies were captured using Image Lab Software. Colony quantification was performed using Image J software, and the data were presented as a percentage relative to untreated colonies.

### Cell cycle analysis

DMG cells were treated for 48 hours with CBL0137, JQ1, or trotabresib at cell line-specific concentrations. For SU-DIPGVI cells, 1 µM for each drug. For RA055, the doses were CBL0137 (0.4 µM), JQ1 (0.1 µM), and trotabresib (0.2 µM). HSJD-DIPG007 cells were treated with CBL0137 (0.2 µM), JQ1 (1 µM), and trotabresib (1 µM). Cell cycle analysis of treated DMG cells was performed as previously described [121].

### ATAC-seq

Neurospheres were treated for 4 h with CBL0137 and JQ1 (HSJD-DIPG007 1 µM CBL0137 1 µM JQ1; SU-DIPGVI 2 µM CBL0137 1 µM JQ1). DMSO was added to the control and single drug treatments to equal concentration in all conditions (maximum 0.03% DMSO). Cells were collected for both ATAC-seq and RNA-extraction. Tn5 Transposition was performed as described in the OMNI-ATAC-seq protocol [122]. Briefly, single cell suspensions from neurospheres were prepared as described for CUT&RUN and washed in ice-cold PBS and cell viability was confirmed to be over 95%. Nuclei from 50,000 viable cells were isolated, permeabilized and transposed with 2.1 µL Tn5 (Illumina) for 30 min with mixing (1000 rpm). DNA was then purified, and PCR amplified (7-14 cycles) with sequencing primers to generate libraries for sequencing. Library concentration was determined using a Qubit Fluorometers (Thermo) (HS dsDNA) and library size determined using a 4200 Tapestation system on a high sensitivity D1000 ScreenTape (Agilent). ATAC-seq libraries were sequenced with a minimum of 50 million paired-end reads per sample and read length 100bp. ATAC-seq experiments were performed in biological duplicate (HSJD-DIPG007) or triplicate (SU-DIPGVI).

### ATAC-seq analysis

ATAC-seq reads were trimmed with Trimmomatic (ILLUMINACLIP:Nextera.fa:2:30:10:8:TRUE SLIDINGWINDOW:5:20 LEADING:3 TRAILING:3 MINLEN:20 CROP:130 HEADCROP:150). Trimmed reads were mapped with bowtie2 using default settings [123] and the resulting BAM files deduplicated using Sambamba markdup [124]. Deduplicated BAM files were then filtered to remove reads with fragment sizes greater than 100bp and CPM BigWigs were generated using the DeepTools bamCoverage [109].

BigWig coverage was visualized as heatmaps and profile plots using DeepTools [109] at hg38 Ensembl genes and at SPT16 peaks and BRD4 peaks. PCA clustering of ATAC-seq average signal over gene bodies was performed using DeepTools. Differential accessibility analysis was carried out using CSAW as described [115], filtering on genomic windows 5-fold higher than background. Regions with an FDR<0.05 were considered significantly different, with ‘opened’ regions logFC > 1 and ‘closed’ regions logFC < -1. Closed regions between the 2 models were overlapped using BedTools [108], annotated using ChIPSeeker [112] and ClusterProfiler was used for Gene Ontology over-representation analysis of the annotated genes [113]. BigWig coverage was visualized at individual gene loci on IGV [111]. A hypergeometric test was used to test if the overlap in ‘closed regions’ between the 2 models was significantly over-enriched (k = 6119 (overlap), s = 11704 (closed in HSJD-DIPG007), M = 41,454 (closed in SU-DIPGXIII), N = 80,894 (number of windows analyzed), where expected = (s*M)/N.

### RNA-seq

DMG cells were treated with CBL0137 and JQ1 for 4 h as for ATAC-seq. Cells were collected for both ATAC-seq and RNA-extraction at the same time. Total RNA was extracted from treated cells using the ISOLATE II RNA Mini Kit from Meridian Bioscience. RNA concentration and purity was determined using a NanoDrop Spectrophotometer and integrity measured using a 4200 Tapestation system. cDNA libraries were prepared using the Illumina Stranded mRNA prep kit. RNA-seq libraries were sequenced with 100bp paired-end reads, with a minimum of 50 million reads per sample. Three biological replicates were performed for all RNA-seq experiments.

### RNA-seq analysis

Illumina paired-end fastq files were aligned to the human genome GRCh38 and gencode v41 using STAR (v2.7.8a) in two-pass mode with quantMode set to TranscriptomeSAM [125]. Transcriptome alignment was quantified using RSEM (v1.3.3) rsem-calculate-expression to obtain raw read counts, and normalized transcripts per million (TPM) and fragments per kilobase million (FPKM) values [126]. Degust (10.5281/zenodo.3258932) was used for differential expression analysis, using edgeR for normalization. Genes with low expression were filtered out using a cutoff of 0.5 counts per million (CPM) in at least 3 samples. Genes with FDR<0.05 and absolute logFC>0.585 (1.5-fold) were considered significantly differentially expressed. Multi-dimensional Scaling (MDS) clustering was performed using Degust. ClusterProfiler was used for Gene Ontology over-representation analysis of the differentially expressed genes [113].

MINTIE [127] was used for differential splicing analysis using parameters assembler=soap, min_logfc=2, min_cpm=0.1, fdr=0.05, and run_de_step=TRUE, with all 3 Vehicle replicates used as control. BedTools [108] was used to determine the common events between the 3 biological replicates. BAM files were visualized on IGV [111] ensuring the same scale.

### TimeLapse RNA-seq

DMG cells were treated with CBL0137 and JQ1 (HSJD-DIPG007 1 µM CBL0137 1 µM JQ1; SU-DIPGVI 2 µM CBL0137 1 µM JQ1) for 2 h. At the same time as drug treatment, 4-Thiouridine (4sU) (ab143718) was added to a final concentration of 100 µM. Cells without 4sU or drug were used as a control. RNA was isolated using a modified version of the RNeasy protocol (Qiagen) as described in [61]. RNA (2 µg) was then subjected to Timelapse chemistry to convert 4sU to C [61]. Libraries were prepared using the Illumina Stranded Total RNA plus RIboZero Plus kit and sequenced on 1 lane of a NovaSeq X Plus 10B 2x150bp, with a minimum 50 million reads per sample. Three biological replicates were performed for TimeLapse RNA-seq experiments.

### TimeLapse RNA-seq analysis

Reads were quality and adapter trimmed using trim-galore 0.6.10 using options –quality 20 -j 4 –paired. Trimmed reads were aligned to the human reference genome hg38 with star [125] 2.7.10 using options --runThreadN 23 --runMode alignReads --readFilesCommand zcat --outReadsUnmapped Fastx --outSAMattributes nM MD –outFileNamePrefix --peOverlapNbasesMin 10 --outFilterMismatchNmax 999 --outFilterMultimapNmax 1 --alignEndsType EndToEnd --outFilterMismatchNoverReadLmax 0.3 --outSAMtype BAM SortedByCoordinate and indexed using samtools 1.6. BAM files were converted to CIT format using the gedi Bam2CIT function. 4sU induced conversions were called using GrandSlam [128] using options -plot -full -D -allGenes -trim5p 5 -trim3p 5. The gene-wise estimates for the new-to-total ratio were input into GrandR [129]. Nascent differential expression was performed using the PairwiseDESeq2 function, with normalization with respect to total. P values were adjusted using Benjamini–Hochberg correction and a cut-off of p.adj < 0.05 and |LFC| > 1 were used to call differentially expressed genes.

### Label-free Total Proteomics Analysis

Following 24 h drug treatment (IC50 doses), cell pellets were washed 2x in PBS and incubated in Lysis Buffer (Cell Signalling, #98030) for 30min on ice with occasional vortexing followed by centrifugation at 13,000 rpm for 15 min at 4°C. Protein concentration was measured with Pierce BCA Protein Assay kit following the manufacturer’s instructions (ThermoScientific, #23225). From each sample, 50µg protein was used for total proteomics analysis. Mass spectrometry analysis was performed at the Bioanalytical Mass Spectrometry Facility at UNSW, Sydney, Australia. Protein masses were detected by an Orbitrap Fusion Lumos Tribrid mass spectrometer (Thermo Scientific, Bremen, Germany) as described previously [130].

Protein peaks were identified using MaxQuant Software and the Andromeda Algorithm version 2.4.13.0 [131] with the following standard settings: oxidation and acetyl (Protein N-term) as variable modifications and carbamidomethyl as fixed modification. Digestion mode was set to trypsin/P with a maximum of 2 missed cleavages. The main search peptide tolerance was +/- 4.5ppm and the isotope match tolerance was set to 2ppm. Protein peaks were searched against the reference human proteome (UniProt, UP000005640). Label-free Quantification (LFQ) was used with standard parameters (minimum ratio 2 and maximum ratio 5) to identify proteins.

Subsequent downstream analysis on the proteomics dataset was performed using Perseus Software (v.2.0.11) to identify differential protein expression [132]. Unnecessary and incorrect protein identifications were filtered out from the data set that were classified as potential contaminants, reverse or only identified by site. The data was then log transformed with base parameter 2, non-valid values were filtered out, and missing values were replaced from normal distribution with 0.3 width and 1.8 downshift standard parameters. Two-sample T-test was used to compare expression levels between vehicle and drug-treated samples. Proteins with a -log p value ≥ 2 were included in downstream pathway analysis. Gene Set Enrichment Pathway analysis was performed in R (v.4.4.2) using the ClusterProfiler package [113].

### siRNA transfection

SF8628 cells were seeded at a density of 10,000 cells/cm^2^ in 24 well plates in antibiotic free media. The following day, siRNA constructs (Dharmacon) were transfected into the cells using DharmaFECT-1 transfection reagent according to the manufacturer’s instructions at a concentration of 20nM. siRNA used were ON-TARGETplus Non-targeting Control Pool (D-001810), ON-TARGETplus MYC (4609) SMARTpool, ON-TARGETplus MDM4 (4194) SMARTpool, and ON-TARGETplus PDGFRA (5156) SMARTpool. For simultaneous knockdown of all 3 genes, total siRNA concentration of 60nM was used. Culture media was changed the following day.

### qPCR

Total RNA was extracted using the ISOLATE II RNA Mini Kit (Meridian Bioscience), and first-strand cDNA was synthesised using the SensiFAST cDNA Synthesis Kit (Meridian Bioscience). Quantitative PCR (qPCR) was performed with the SensiFAST SYBR No-ROX Kit (Meridian Bioscience) using primer sequences listed in **Table 2**, synthesised by Integrated DNA Technologies using a QuantStudio 3 qPCR machine. Relative mRNA expression levels of genes of interest were normalised to corresponding housekeeping genes using the ΔΔCt method [133]. qPCR experiments were performed in biological triplicate.

**Table 2.**
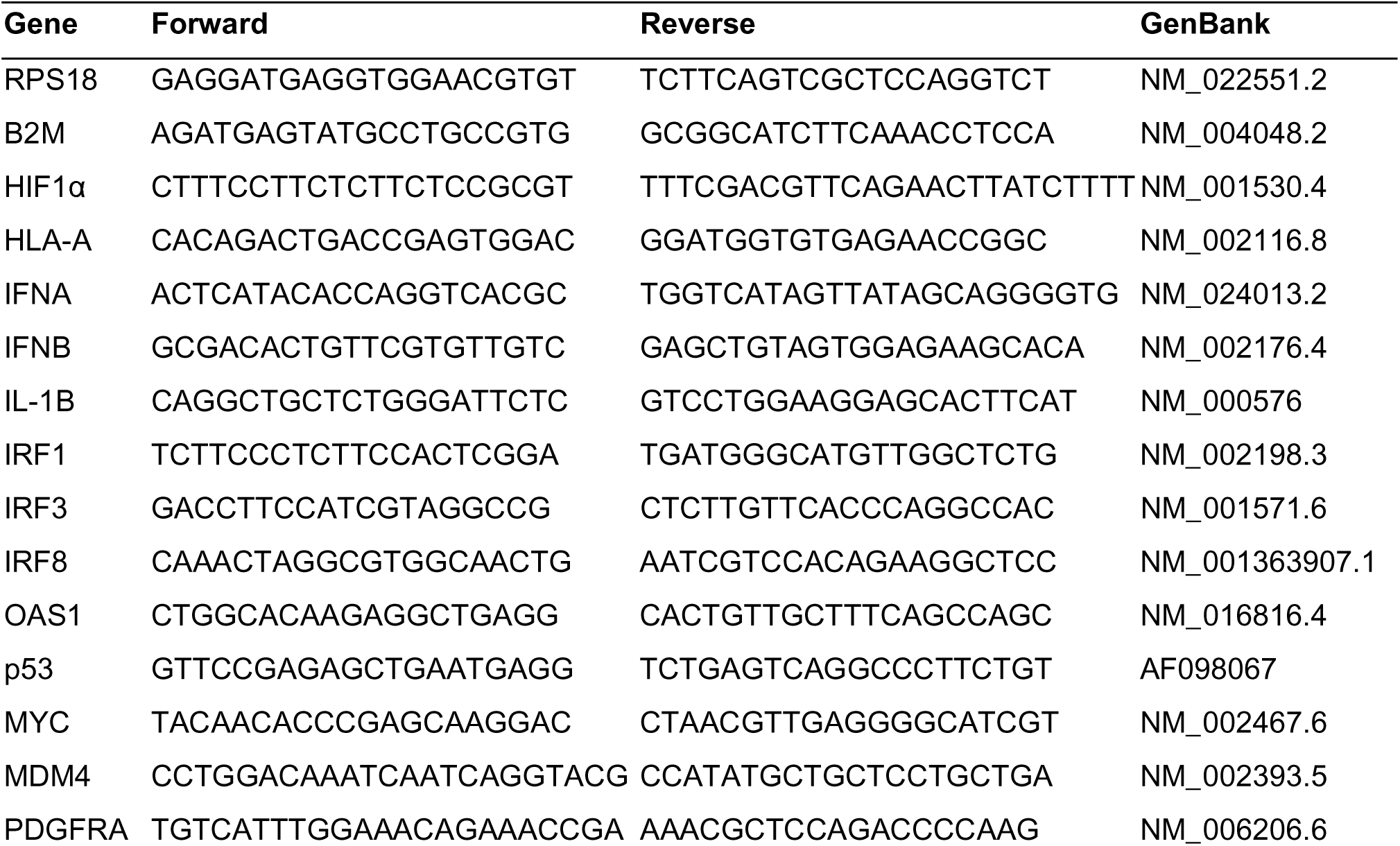
Sequences of qPCR primers.

### Timelapse microscopy proliferation assay

SF8628 cells were transfected with siRNA in duplicate wells, and the following day, the media was changed, and cells were treated with CBL0137 (0.2µM) and JQ1 (0.5µM). Cells were grown for 6 days, and phase-contrast confluence was monitored by time-lapse microscopy every 6 h using an Incuyte S3 Live Cell Analysis System. Incucyte experiments were repeated three times.

### Orthotopic model/drug treatment

The protocol for intracranial injections was implemented as described in the study conducted by Khan et al. [121]. The timing of treatment initiation varied for each model with SU-DIPGVI-Luc at 7 weeks, RA055 at 3 weeks. and HSJD-DIPG007 starting at 5 weeks post intracranial surgery.

In the SU-DIPGVI-Luc and RA055 models, the drug treatments were as follows: CBL0137 - dissolved in 5% dextrose, CBL0137 was administered intravenously at a dosage of 40 mg/kg/day. The treatment was given once every four days for a total of four weeks. JQ1 - dissolved in DMSO and subsequently in a solution containing 0.9% saline with 5% PEG400 and 5% TWEEN-80, JQ1 was administered at a dosage of 25 mg/kg/day intraperitoneally. The treatment was given five days a week over a period of four weeks.

In the HSJD-DIPG007 model, the drug treatments included: CBL0137 - dissolved in 5% dextrose, CBL0137 was administered intravenously at a dosage of 20 mg/kg/day. The treatment was given five days a week for a total of four weeks. JQ1 - Dissolved in DMSO and subsequently in a solution containing 0.9% saline with 5% PEG400 and 5% TWEEN-80, JQ1 was administered intraperitoneally at a dosage of 50 mg/kg/day. The treatment was given five days a week for a duration of four weeks.

For the immunocompetent model the protocol for intracranial injections was implemented as described above with 5-7 weeks old female C57BL6 mice, using IUE 24B7 cells that harbor H3.3K27M mutation [134, 135]. The timing of treatment initiation was 18 days post intracranial surgery. The drug treatments were as follows: CBL0137 - dissolved in 5% dextrose, CBL0137 was administered intravenously at a dosage of 40 mg/kg/day. The treatment was given once every four days for a total of three weeks. JQ1 - dissolved in DMSO and subsequently in a solution containing 0.9% saline with 5% PEG400 and 5% TWEEN-80, JQ1 was administered at a dosage of 50 mg/kg/day intraperitoneally. The treatment was given five days a week over three weeks. In combination group, CBL0137 was administered before JQ1. Throughout the study, the survival of the mice was closely monitored daily for neurological and clinical symptoms as described. Brain samples were collected at the end of the treatment period.

Throughout the study, the survival of the mice was closely monitored daily. Neurological and clinical symptoms were assessed to detect signs of tumor development. Mice exhibiting clinical signs of neurological decline, such as ataxia, circling, head tilting, with or without a 20% weight loss, were humanely euthanized. All animal experiments strictly adhered to the guidelines set by the Animal Care and Ethics Committee of the University of New South Wales and the Australian Code of Practice for the Care and Use of Animals for Scientific Purposes. The experimental protocols used in this study were approved under protocol number 22/46B.

### Hematologic and Biochemical Analysis

Hematologic and biochemical analyses were performed as previously described by Khan et al [121]. Briefly, healthy female BALB/c mice (12-14 weeks old) were treated four weeks with either vehicle or the indicated compounds, with three mice per treatment cohort. The study included six experimental groups: vehicle control (buffer A administered intraperitoneally and 5% dextrose intravenously), CBL0137 (40 mg/kg intravenously), JQ1 (25 mg/kg intraperitoneally), ZEN3694 (Selleckchem E1517) (50 mg/kg orally), CBL0137 + JQ1 combination, and CBL0137 + ZEN3694 combination. ZEN3694 was dissolved in a vehicle consisting of 5% DMSO, 40% PEG300, 5% Tween 80, and 50% ddH₂O. JQ1 was dissolved in DMSO to prepare a 10X stock solution, which was subsequently diluted with 9 volumes of buffer A. CBL0137 was dissolved in 5% dextrose. CBL0137 was administered once every four days via intravenous injection. JQ1 and ZEN3694 were administered on a 5 days on, 2 days off schedule via intraperitoneal for JQ1 and by oral gavage for ZEN3694. Vehicle control animals received intravenous 5% dextrose once every four days and intraperitoneal buffer A on weekdays to match the treatment schedules of the active compounds. Weekly tail vein blood samples were collected for hematological and biochemical analysis throughout the treatment period. For hematology, blood was collected in EDTA-containing tubes (Sarstedt, cat # 20.1339.100) (15 μL) and diluted by a factor of 2 with PBS before loading to the machine to avoid clot formation, then analyzed using a Mindray BC-5000 hematology analyzer. For biochemistry, blood samples were pooled from each cage and collected in lithium heparin microtainer tubes (Sarstedt, cat # 20.1345.100) (100 μL), then analyzed using an Abaxis VetScan Biochemistry Analyser with disposable rotors (Zoetis, Australia; Comprehensive diagnostic profile cat # 10023220). On the final day of treatment, animals were euthanised and samples collected. Blood was obtained via cardiac puncture for hematology, biochemistry, and plasma isolation.

### LC-MS/MS determination of drug concentration in blood and tissue samples

Mice were treated with the indicated compounds as described in the previous section. Brain tissue samples (100 mg) were suspended in 900 μL methanol + 0.1% formic acid and homogenized for compound extraction. Following homogenization, samples were agitated at 4°C for 30 minutes, then centrifuged at 17,000g. The resulting supernatant was collected for mass spectrometry analysis. Plasma samples (50-100 μL) were mixed with 9 volumes of ice-cold acetonitrile for protein precipitation. After incubation on ice for 5 minutes, samples were centrifuged at 17,000g and the supernatant collected for mass spectrometry analysis. All mass spectrometry analyses, including calibration curve preparation and sample analysis, were performed by the Mass Spectrometry Facility at the University of New South Wales (UNSW BMSF).

### Immunohistochemistry analysis of brain tissue

Brain samples were collected on the final day of treatment and fixed in a 10% formalin neutral buffered solution (Sigma-Aldrich). The fixed samples were then embedded in paraffin wax, and 5 µm sections were cut and mounted on glass slides. To prepare the sections for histological examination, dehydration was carried out using standard procedures. Subsequently, the sections were stained with hematoxylin/eosin and Ki67, H3K27ac, MDM4, and MYC (**Table 1**) using the BOND RX Fully Automated Research Stainer (Leica biosystems) for histological examination. Following staining, positive cells were identified and quantified using Andy’s Algorithms [136] on the image analysis pipeline on FIJI.

### Surface marker analysis by flow cytometry

DMG cells were treated with CBL0137 and JQ1 (HSJD-DIPG007 1 µM CBL0137 1 µM JQ1; SU-DIPGVI 2 µM CBL0137 1 µM JQ1) for 24 hours. Following treatment, cells were harvested, blocked with FcR blocking reagent for 10 min at 4°C, and stained with HLA-ABC-VioBlue and HLA-E-PE antibodies (1:50 dilution; Table 1) for an additional 10Cmin at 4°C. Washed cells were analyzed on a MACSQuant VYB Flow Cytometer and data processed using FlowJo software. Experiments were performed in at least three biological replicates.

Human monocyte derived macrophages were treated in technical duplicate for the following groups: M0 (media only), M1 (proinflammatory, LPS 1 ng/mL and IFNγ 20 ng/mL), M2 (anti-inflammatory, IL-4 20 ng/mL and IL-13 20 ng/mL), CBL1037 1 µM, JQ1 1 µM, and combination of the two drugs. After 48 hours, media containing treatment in plates were aspirated and replaced with warm Gey’s Balanced Salt Solution. Cells were then stained with CD86-FITC (1:50) and CD206-APC (1:50) (BD Biosciences) (**Table 1**) as per manufacturer’s protocol. After 1 hour of staining, cells were then washed with Gey’s Balanced Salt Solution and harvested using cell scrapers into FACS tubes for flow cytometry. Analysis was performed using FlowJo. Experiments were performed in biological triplicate.

### Phagocytosis Assay

pHrodo™ BioParticles™ conjugate (Cat. A10010, Thermo Fisher) was diluted in Gey’s solution to a final volume of 2 mL per vial. The suspension was transferred to a 20 mL glass Schott bottle and sonicated in an ultrasonic water bath at room temperature for 10 minutes to ensure uniform particle dispersion. For each well of HMDM cells seeded in a 12-well plate, 100 μL of the bead suspension was added, maintaining the manufacturer-recommended 1:10 ratio. No-cell background controls were similarly prepared. Following bead addition, cells were incubated at 37°C for 2–3 hours. Fluorescence imaging was performed using a Floid Cell Imaging Station (Thermo Fisher Scientific) with the RFP filter. Fluorescent signal per cell was quantified using ImageJ. A cell mask was generated, and total fluorescence intensity was divided by the number of cells per field of view. The experiment was performed in duplicate.

### Xenium In situ spatial profiling

Formalin-fixed brains were sectioned at 5 µm and mounted onto Xenium slides, with two brains per slide. Xenium Prime In Situ (10x Genomics) was performed according to the manufacturer’s instructions using the 5000 pre-designed gene panel and cell segmentation staining. Briefly, priming oligos were hybridized to the target RNA, followed by site-specific cleavage using RNase treatment to release the RNA strand from the hybridized oligo. The 5000-gene panel was then added to the sections to hybridize with the RNA, followed by ligation to generate circular DNA probes, which were subsequently amplified via rolling circle amplification. After a series of ethanol washes, the tissue was blocked and stained with the cell segmentation stain and DAPI. Slides were then loaded into the Xenium Analyzer for imaging and decoding. Spatial Transcriptomics Services for this study were provided by the Garvan Genomics Platform at the Garvan Institute of Medical Research.

### Xenium In situ spatial profiling analysis

Xenium output from the four treatment conditions were merged into a single Seurat object (Seurat v5.2.1). Cells with transcript counts <10 and above the 98^th^ percentile were filtered, and data were normalized with the median transcript count. The Seurat object was down sampled (25,000 cells per sample) using SketchData and LeverageScore to retain rare cell populations for subsequent scaling and clustering. From the subsetted data, variable features (n = 2000 genes) were identified using FindVariableFeatures, data were scaled using ScaleData and Dimensionality reduction performed using RunPCA. Batch correction was performed using Harmony Integration, followed by UMAP projection and clustering in Harmony space. Clustering was then projected to all cells. Integrated clusters were exported for visualization using Xenium Explorer (v3.0). Marker genes for each cluster were identified using FindAllMarkers. Tumour and immune clusters of interest were subset and re-analyzed with feature selection, scaling, PCA, Harmony integration, and UMAP clustering. Differential expression analysis between vehicle and combination treatment groups in select clusters was performed using FindMarkers. Genes expressed in fewer than 15% of cells in either group were excluded prior to GSEA using ClusterProfiler.

## Data Availability

Raw and processed NGS data has been uploaded onto the GEO repository. CUT&RUN (GSE272130), ATAC-Seq (GSE272129), nascent RNA-seq (GSE299509), RNA-seq (GSE272131), and Xenium (GSE299434).

## Acknowledgments

We would like to express our gratitude to Michelle Monje, Angel Montero Carcaboso, and Esther Hulleman for their generous provision of the SU-DIPG, HSJD-DIPG, and VUMC-DIPG10 cells, respectively, which were instrumental in the success of this study. We thank Nada Jabado and Brian Krug for providing the isogenic H3K27M KO SU-DIPGXIII models and Yunjia Zhang, Oluwaseyi Samson Ogunmodede, and Kerry-Anne Rye for providing the HMDM cells. We thank Katerina Gurova and Andrei Gudkov for providing the CBL0137 drug. We also acknowledge the Core Services Group at Children’s Cancer Institute for their valuable technical support throughout the research. We extend our appreciation to the Flow Cytometry Core Facility, BRIL Facility, and BMSF Facility (Russel Pickford) at UNSW assistance in flow cytometry, imaging, and pharmacokinetic testing, respectively, which greatly contributed to our data analysis and interpretation. We also acknowledge the Garvan Institute Histopathology Facility and Garvan Genomics Platform for their services. Furthermore, we would like to acknowledge the Zero Childhood Cancer Program and the Tumour Bank teams at CCI for their significant contributions to the establishment of primary DMG cultures, which were indispensable for our investigations. The authors thank the Children’s Cancer Institute Animal Facility for providing support to this study. We would like to thank Nathan Dahl and Lays Sobral for technical advice regarding CUT&RUN and ATAC-seq, and John Reeves and Quy Xiao Xuan Lin for technical support for spatial analyses. We would like to also acknowledge Vyoma Patel for proofreading this manuscript. This work has generously been funded by the Kids Cancer Alliance, We Love You Connie Foundation Grant, Levi’s Project, CINSW Fellowship (HH) (2022/ECF1425), The Kids Cancer Project Col Reynolds Fellowship (AK), NHMRC Synergy Grant (2019056), Cancer Institute NSW Translational Program Grant (2019/TPG2037), NHMRC Investigator grant (GNT2017898), and DIPG Collaborative: The Cure Starts Now (2018). Graphical abstract and treatment schemes created with BioRender.com.

## Author Contributions

**Conceptualization**: MT, DSZ.

**Methodology:** HH, AK, JL.

**Formal Analysis:** HH, AK, AE, NJ, SER, XG, HN, CM, BR, DK, CS.

**Investigation:** HH, AK, AE, AG, HN, AL, BR, MT, EW, DL, RL, KI, WL, RS, LRS.

**Data Curation:** HH, AK, CM, NJ, SER, CXS.

**Visualization:** HH, AK, SER, YCS, BR.

**Writing – Original Draft:** HH, AK.

**Writing – Review & Editing:** All.

**Supervision:** RF, RJW, FVM, TNP, CM, BR, MT, MD, DSZ.

**Project Administration:** HH, AK.

**Funding Acquisition:** HH, BR, MT, DSZ.

