## Supplemental Figures 1-8 and Supplemental Tables 2-10 for "Chromatin remodeling with combined FACT and BET inhibition disrupts oncogenic transcription in Diffuse Midline Glioma"

### Supplementary Data

- Graphical abstract
- Supplementary Figures 1-8
- Supplementary Tables 2-10

**Chromatin remodeling with combined FACT and BET inhibition disrupts oncogenic transcription in Diffuse Midline Glioma**

**Authors:** Holly Holliday[1,2*], Aaminah Khan[1,2*], Anahid Ehteda[1,2], Hieu Nguyen[1], Samuel E. Ross[3], Nisitha Jayatilleke[1], Anjana Gopalakrishnan[1], Eyden Wang[1], Yolanda Colino Sanguino[1,2], Daisy Kavanagh[4,5], Xinyi Guo[1,6], Jie Liu[1], David Lawrence[1,2], Claire X. Sun[6,7], Rebecca Lehmann[1,2], Chi Kin Ip[1,2], Alvin Lee[1], Laura Rangel-Sanchez[4,8], Wenyan Li[1], Robert Salomon[1,2], Ron Firestein[6,7], Robert J Weatheritt[4,5], Fatima Valdes-Mora[1,2], Marcel E. Dinger[3], Timothy N. Phoenix[9], Chelsea Mayoh[1,2], Benjamin S. Rayner[1,2], Maria Tsoli[1,2**], David S. Ziegler[1, 2, 10**]

*, ** equal contribution

**Affiliations:**

1. Children's Cancer Institute, Lowy Cancer Research Centre, UNSW Sydney, Sydney, NSW, Australia
2. School of Clinical Medicine, UNSW Sydney, Sydney, NSW, Australia
3. School of Life and Environmental Sciences, University of Sydney, NSW, Australia
4. Garvan Institute of Medical Research, Sydney, NSW, Australia
5. School of Biotechnology and Biomolecular Sciences, UNSW Sydney, Sydney, NSW, Australia
6. Centre for Cancer Research, Hudson Institute of Medical Research, Clayton, Australia
7. Department of Medicine, School of Clinical Sciences, Monash University, Clayton, Australia
8. School of Biomedical Engineering, Faculty of Engineering and Information Technology, University of Technology Sydney, Sydney, NSW Australia
9. Division of Pharmaceutical Sciences, James L. Winkle College of Pharmacy, University of Cincinnati, Cincinnati, OH, USA
10. Kid's Cancer Centre, Sydney Children's Hospital, Randwick, NSW, Australia

#### Graphical Abstract

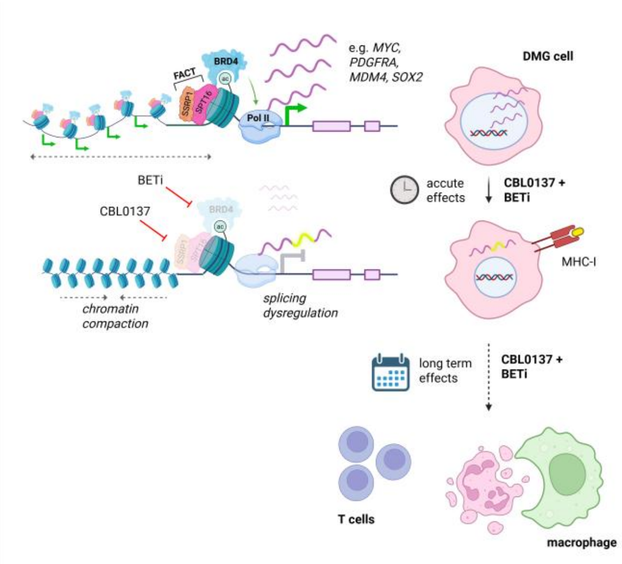

Fig S1

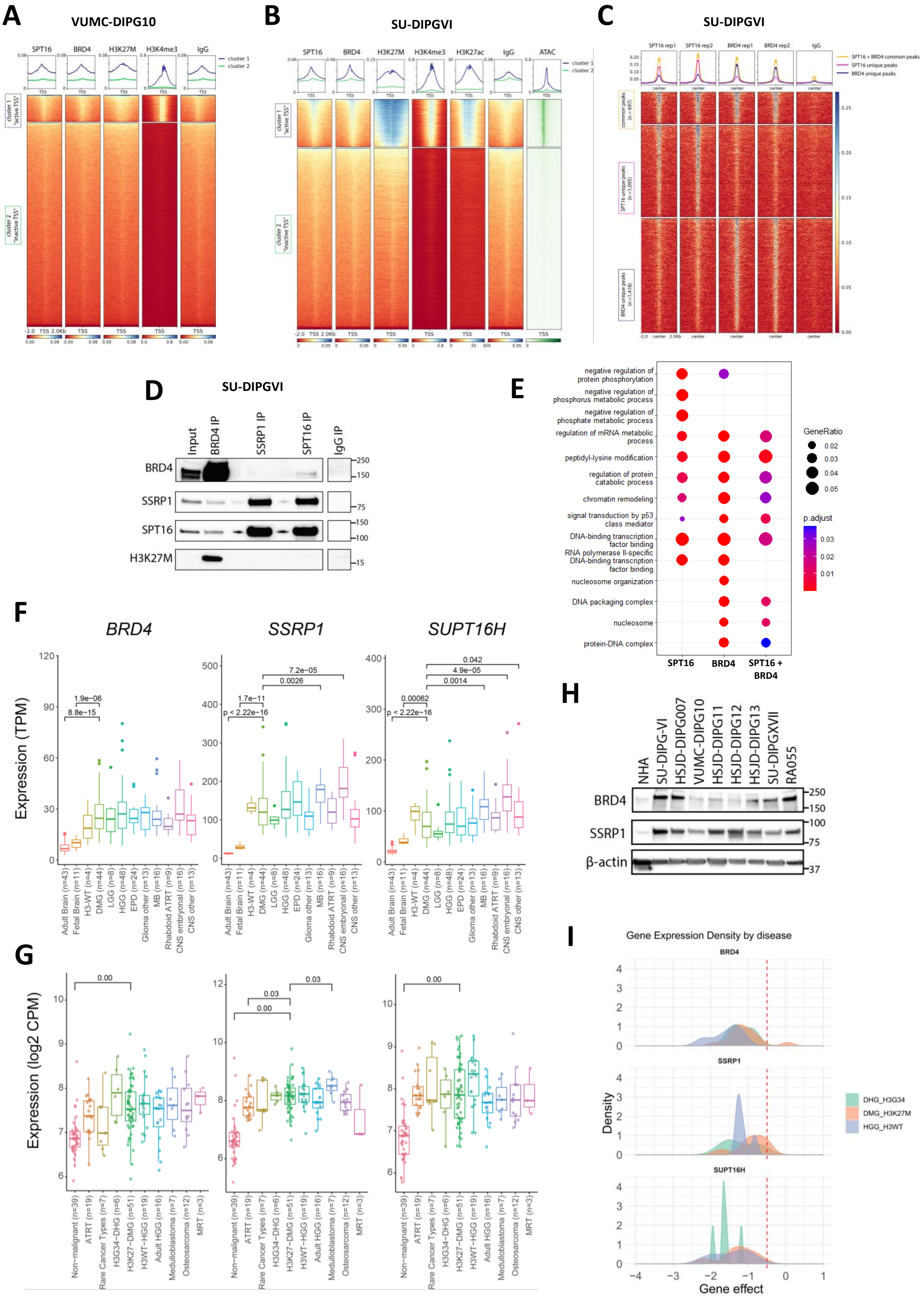
A
VUMC-DIPG10
SPT16
BRD4
H3K27M
H3K4me3
IgG
cluster 1
cluster 2
cluster 1 "active TSS"
cluster 2 "inactive TSS"
B
SU-DIPGVI
SPT16
BRD4
H3K27M
H3K4me3
H3K27ac
IgG
ATAC
C
SU-DIPGVI
SPT16 rep1
SPT16 rep2
BRD4 rep1
BRD4 rep2
IgG
SPT16 + BRD4 common peaks
SPT16 unique peaks
BRD4 unique peaks
D
SU-DIPGVI
Input
BRD4 IP
SSRP1 IP
SPT16 IP
IgG IP
BRD4
SSRP1
SPT16
H3K27M
250
150
75
150
100
15
E
negative regulation of protein phosphorylation
negative regulation of phosphorus metabolic process
negative regulation of phosphate metabolic process
regulation of mRNA metabolic process
peptidyl-lysine modification
regulation of protein catabolic process
chromatin remodeling
signal transduction by p53 class mediator
DNA-binding transcription factor binding
RNA polymerase II-specific DNA-binding transcription factor binding
nucleosome organization
DNA packaging complex
nucleosome
protein-DNA complex
SPT16
BRD4
SPT16 + BRD4
GeneRatio
0.02
0.03
0.04
0.05
p.adjust
0.03
0.02
0.01
F
BRD4
SSRP1
SUPT16H
Expression (TPM)
1.9e−06
8.8e−15
1.7e−11
p < 2.22e−16
7.2e−05
0.0026
0.00062
p < 2.22e−16
0.042
4.9e−05
0.0014
Adult Brain (n=43)
Fetal Brain (n=11)
H3-WT (n=4)
DMG (n=44)
LGG (n=8)
HGG (n=48)
EPD (n=24)
Glioma other (n=13)
MB (n=16)
Rhabdoid ATRT (n=9)
CNS embryonal (n=16)
CNS other (n=13)
G
Expression (log2 CPM)
0.00
0.03
0.00
0.03
0.00
Non−malignant (n=39)
ATRT (n=19)
Rare Cancer Types (n=7)
H3G34−DHG (n=6)
H3K27−DMG (n=51)
H3WT−HGG (n=19)
Adult HGG (n=16)
Medulloblastoma (n=7)
Osteosarcoma (n=12)
MRT (n=3)
H
NHA
SU-DIPG-VI
HSJD-DIPG007
VUMC-DIPG10
HSJD-DIPG11
HSJD-DIPG12
HSJD-DIPG13
SU-DIPGXVII
RA055
BRD4
SSRP1
β-actin
250
150
100
75
37
I
Gene Expression Density by disease
BRD4
SSRP1
SUPT16H
Density
Gene effect
DHG_H3G34
DMG_H3K27M
HGG_H3WT

**Figure S1. FACT and BRD4 expression and co-localization in DMG. Related to Figure 1.**
**A-B**) Profile plots and heatmaps depicting SPT16, BRD4, H3K27M, H3K4me3, IgG CUT&RUN signal in VUMC-DIPG10 (**A**) and SU-DIPGVI (**B**) neurospheres. Heatmaps are ranked based on SPT16 signal and separated into two k-means clusters. Representative data from n=2 experiments. For SU-DIPGVI, H3K27ac CUT&RUN from [47] and ATAC-seq signal is shown. **C)** Profile plots and heatmaps depicting the average SPT16, BRD4 and IgG CUT&RUN signal at SPT16 + BRD4 common peaks (orange), SPT16 unique peaks (pink), BRD4 unique peaks (blue) in SU-DIPGVI cells. **D**) Co-Immunoprecipitation and western blotting in SU-DIPGVI nuclear extracts showing direct interactions between BRD4 with the FACT subunits and H3K27M. **E**) Dot plot of over-represented gene ontologies for genes associated with SPT16 peaks, BRD4 peaks, and SPT16+BRD4 common peaks in HSJD-DIPG007 cells. **F**) Expression of FACT subunit genes *SSRP1* and *SUPT16H*, and *BRD4* in DMG and other CNS tumors compared to normal fetal and adult brain. RNA-seq data for tumor samples were obtained from the ZCC clinical trial [55] and normal brain samples from McGill University for NFB [56]. **G**) Expression of FACT subunit genes *SSRP1* and *SUPT16H*, and *BRD4* in H3K27M DMG cell lines compared to other pediatric cancer cell lines and to non-malignant cell lines. DMG samples were compared to all other samples using independent two-sample t-tests. **H)** Western blot for SSRP1 and BRD4 in a panel of DMG cell lines compared to Normal Human Astrocytes (NHA). **I**) CRISPR screen density plot showing gene dependency scores for *BRD4*, *SSRP1*, and *SUPT16H* across glioma subtypes. Gene effect score distributions are shown for H3G34-mutant Diffuse Hemispheric Glioma (green), H3K27M DMG (pink), and H3 wild-type high-grade glioma (blue). A threshold of < –0.5 (dashed line) indicates gene dependency.

#### Fig S2

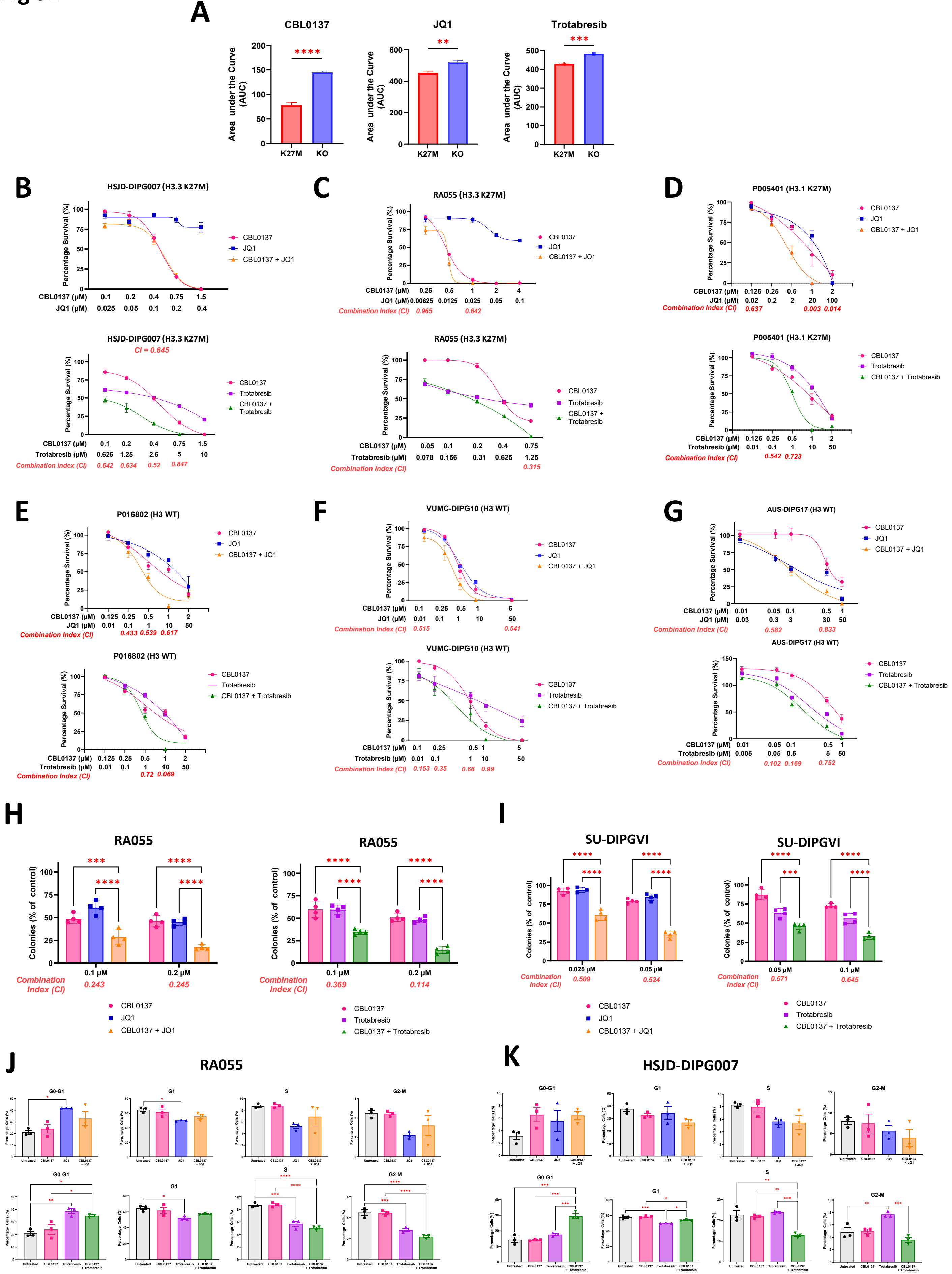

A
CBL0137
JQ1
Trotabresib
Area under the Curve (AUC)
K27M
KO
****
**
***
B
HSJD-DIPG007 (H3.3 K27M)
Percentage Survival (%)
CBL0137
JQ1
CBL0137 + JQ1
CBL0137 (µM) 0.1 0.2 0.4 0.75 1.5
JQ1 (µM) 0.025 0.05 0.1 0.2 0.4
C
RA055 (H3.3 K27M)
CBL0137 (µM) 0.25 0.5 1 2 4
JQ1 (µM) 0.00625 0.0125 0.025 0.05 0.1
Combination Index (CI) 0.965 0.642
D
P005401 (H3.1 K27M)
CBL0137 (µM) 0.125 0.25 0.5 1 2
JQ1 (µM) 0.02 0.2 2 20 100
Combination Index (CI) 0.637 0.003 0.014
HSJD-DIPG007 (H3.3 K27M)
CI = 0.645
Trotabresib
CBL0137 + Trotabresib
CBL0137 (µM) 0.1 0.2 0.4 0.75 1.5
Trotabresib (µM) 0.625 1.25 2.5 5 10
Combination Index (CI) 0.642 0.634 0.52 0.847
RA055 (H3.3 K27M)
CBL0137 (µM) 0.05 0.1 0.2 0.4 0.75
Trotabresib (µM) 0.078 0.156 0.31 0.625 1.25
Combination Index (CI) 0.315
P005401 (H3.1 K27M)
CBL0137 (µM) 0.125 0.25 0.5 1 2
Trotabresib (µM) 0.01 0.1 1 10 50
Combination Index (CI) 0.542 0.723
E
P016802 (H3 WT)
CBL0137 (µM) 0.125 0.25 0.5 1 2
JQ1 (µM) 0.01 0.1 1 10 50
Combination Index (CI) 0.433 0.539 0.617
F
VUMC-DIPG10 (H3 WT)
CBL0137 (µM) 0.1 0.25 0.5 1 5
JQ1 (µM) 0.01 0.1 1 10 50
Combination Index (CI) 0.515 0.541
G
AUS-DIPG17 (H3 WT)
CBL0137 (µM) 0.01 0.05 0.1 0.5 1
JQ1 (µM) 0.03 0.3 3 30 50
Combination Index (CI) 0.582 0.833
P016802 (H3 WT)
CBL0137 (µM) 0.125 0.25 0.5 1 2
Trotabresib (µM) 0.01 0.1 1 10 50
Combination Index (CI) 0.72 0.069
VUMC-DIPG10 (H3 WT)
CBL0137 (µM) 0.1 0.25 0.5 1 5
Trotabresib (µM) 0.01 0.1 1 10 50
Combination Index (CI) 0.153 0.35 0.66 0.99
AUS-DIPG17 (H3 WT)
CBL0137 (µM) 0.01 0.05 0.1 0.5 1
Trotabresib (µM) 0.005 0.05 0.5 5 50
Combination Index (CI) 0.102 0.169 0.752
H
RA055
Colonies (% of control)
0.1 µM 0.2 µM
Combination Index (CI) 0.243 0.245
Combination Index (CI) 0.369 0.114
I
SU-DIPGVI
0.025 µM 0.05 µM
Combination Index (CI) 0.509 0.524
0.05 µM 0.1 µM
Combination Index (CI) 0.571 0.645
J
RA055
K
HSJD-DIPG007
G0-G1
G1
S
G2-M
Percentage Cells (%)
Untreated

**Figure S2. In vitro therapeutic efficacy of combinatorial FACT and BRD4 inhibition. Related to Figure 2.**
**A)** Area under the curve (AUC) values for dose-response curves comparing SU-DIPGXIII isogenic cells -/+ H3K27M. P-values were calculated using a two-tail t-test. (**B-D**) Combination dose-response curves for Histone mutant HSJD-DIPG007 (**B**), RA055 (**C**), and P005401 (**D**) cells treated for 72 hours (**E-G**) Combination dose-response curves for Histone wildtype P016802 **(E)**, VUMC-DIPG10 (**F**), and AUS-DIPG17 (**G**) cells treated for 72 hours. All dose response curves are represented as percentage viability compared to untreated controls, mean ± SEM (n = 3). Combination index (CI) values were calculated using Calcusyn, with synergistic CI values indicated. (**H-I**) Colony formation assays in RA055 (**H**) and SU-DIPGVI (**I**) cells treated with the indicated drugs for 2 or 4 weeks. Data is presented as mean ± SEM (n=4). (**J-K**) Flow cytometry cell cycle distributions in RA055 (**J**) and HSJD-DIPG007 (**K**) cells treated with CBL0137 combined with JQ1 or trotabresib for 48 hours (n=3). Significance was calculated using a one-way ANOVA with Tukey's multiple comparisons test for single and combination treatments. * $p<0.05$, ** $p<0.01$, *** $p<0.001$, **** $p<0.0001$.

Fig S3

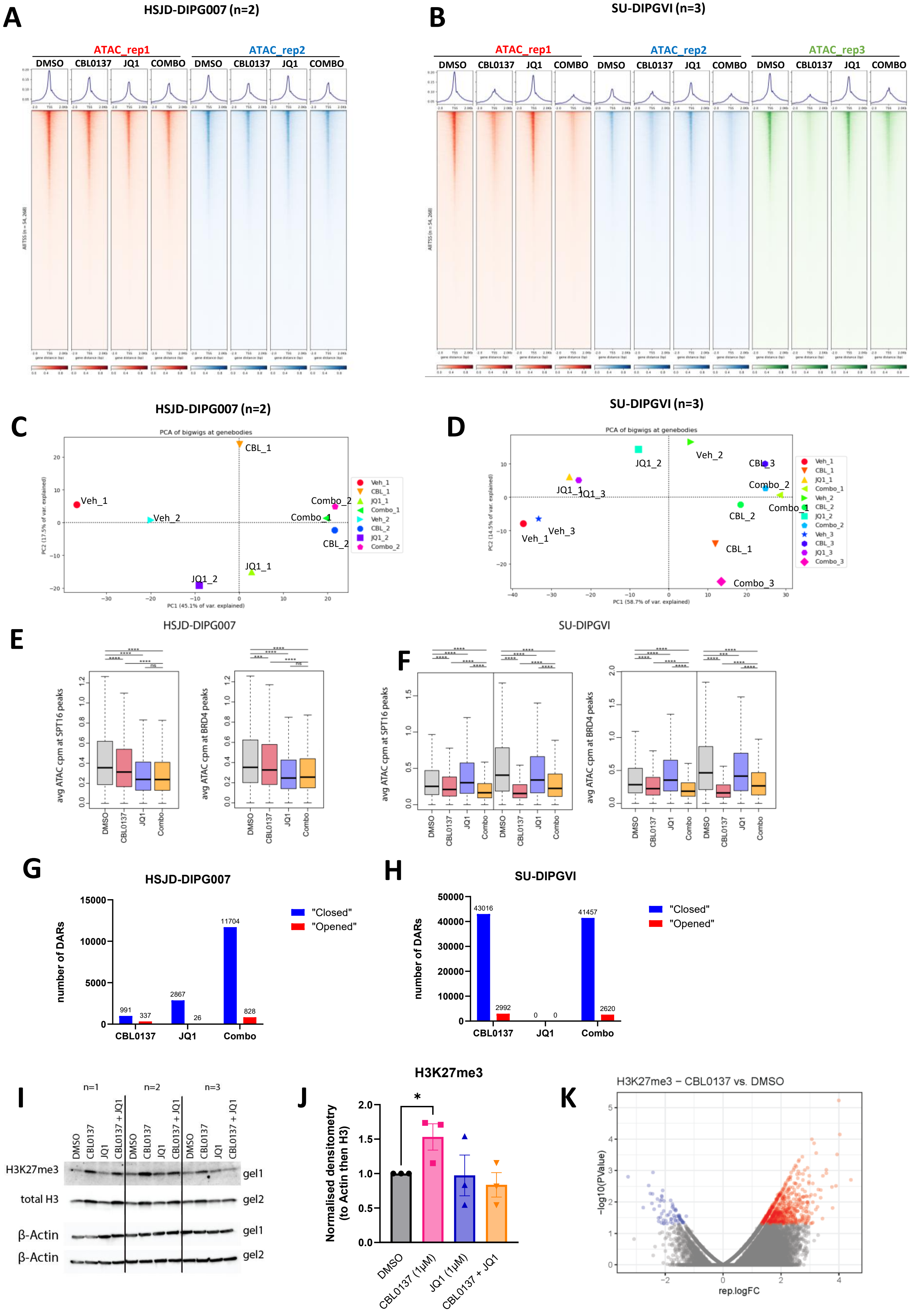
A
HSJD-DIPG007 (n=2)
ATAC_rep1
ATAC_rep2
DMSO
CBL0137
JQ1
COMBO
B
SU-DIPGVI (n=3)
ATAC_rep1
ATAC_rep2
ATAC_rep3
C
HSJD-DIPG007 (n=2)
PCA of bigwigs at genebodies
PC1 (45.1% of var. explained)
PC2 (17.5% of var. explained)
Veh_1
Veh_2
CBL_1
CBL_2
JQ1_1
JQ1_2
Combo_1
Combo_2
D
SU-DIPGVI (n=3)
PCA of bigwigs at genebodies
PC1 (58.7% of var. explained)
PC2 (14.5% of var. explained)
Veh_3
CBL_3
JQ1_3
Combo_3
E
HSJD-DIPG007
avg ATAC cpm at SPT16 peaks
avg ATAC cpm at BRD4 peaks
ns
Combo
F
SU-DIPGVI
G
HSJD-DIPG007
number of DARs
"Closed"
"Opened"
991
337
2867
26
11704
828
CBL0137
JQ1
Combo
H
SU-DIPGVI
43016
2992
0
0
41457
2620
I
n=1
n=2
n=3
CBL0137 + JQ1
H3K27me3
total H3
β-Actin
gel1
gel2
J
H3K27me3
Normalised densitometry (to Actin then H3)
*
DMSO
CBL0137 (1μM)
JQ1 (1μM)
CBL0137 + JQ1
K
H3K27me3 – CBL0137 vs. DMSO
−log10(PValue)
rep.logFC

**Figure S3. CBL0137 and JQ1 condense chromatin at gene promoters. Related to Figure 3.**
**A-B)** Heatmaps depicting ATAC-seq signal at transcription start sites in HSJD-DIPG007 (n=2) (**A**) and SU-DIPGVI (n=3) (**B**) cells treated with DMSO, CBL0137 (1µM for HSJD-DIPG007 and 2µM for SU-DIPGVI), JQ1 (1µM) or the combination for 4 hours. Average signal is shown in the profile plots above. **C-D)** Principal component analysis (PCA) plots showing the clustering of samples based on signal at TSS in HSJD-DIPG007 (**C**) and SU-DIPGVI (**D**). **E-F**) Box plots displaying ATAC-seq signal at SPT16 peaks (left) and BRD4 peaks (right) in DMG cells treated with DMSO, CBL0137 (1µM for HSJD-DIPG007 and 2µM for SU-DIPGVI), JQ1 (1µM) or the combination for 4 hours. A Wilcox test was used to test significance between treatment groups. * $p<0.05$, ** $p<0.01$, *** $p<0.001$, **** $p<0.0001$. **G-H**) Bar charts depicting the number of significantly differentially accessible regions (DARs). Blue represents significantly (FDR < 0.05, LFC < -1) downregulated "closed" regions, red data represents significantly upregulated "opened" (LFC > 1). **I**) Western blot for H3K27me3, total H3, and β-actin in HSJD-DIPG007 cells treated for 24 hours with the indicated treatment groups. **J)** Densitometry bar graphs (mean +/- SEM; n=3) normalized to β-actin and total H3, and expressed as a fold-change relative to the DMSO control. An unpaired t-test was used to calculate significance. * $p<0.05$. **K**) H3K27me3 CUT&RUN volcano plot displaying CSAW differential binding analysis in HSJD-DIPG007 cells treated with 1µM CBL0137 for 4h. Significant threshold p-val<0.05 (n=2).

Fig S4

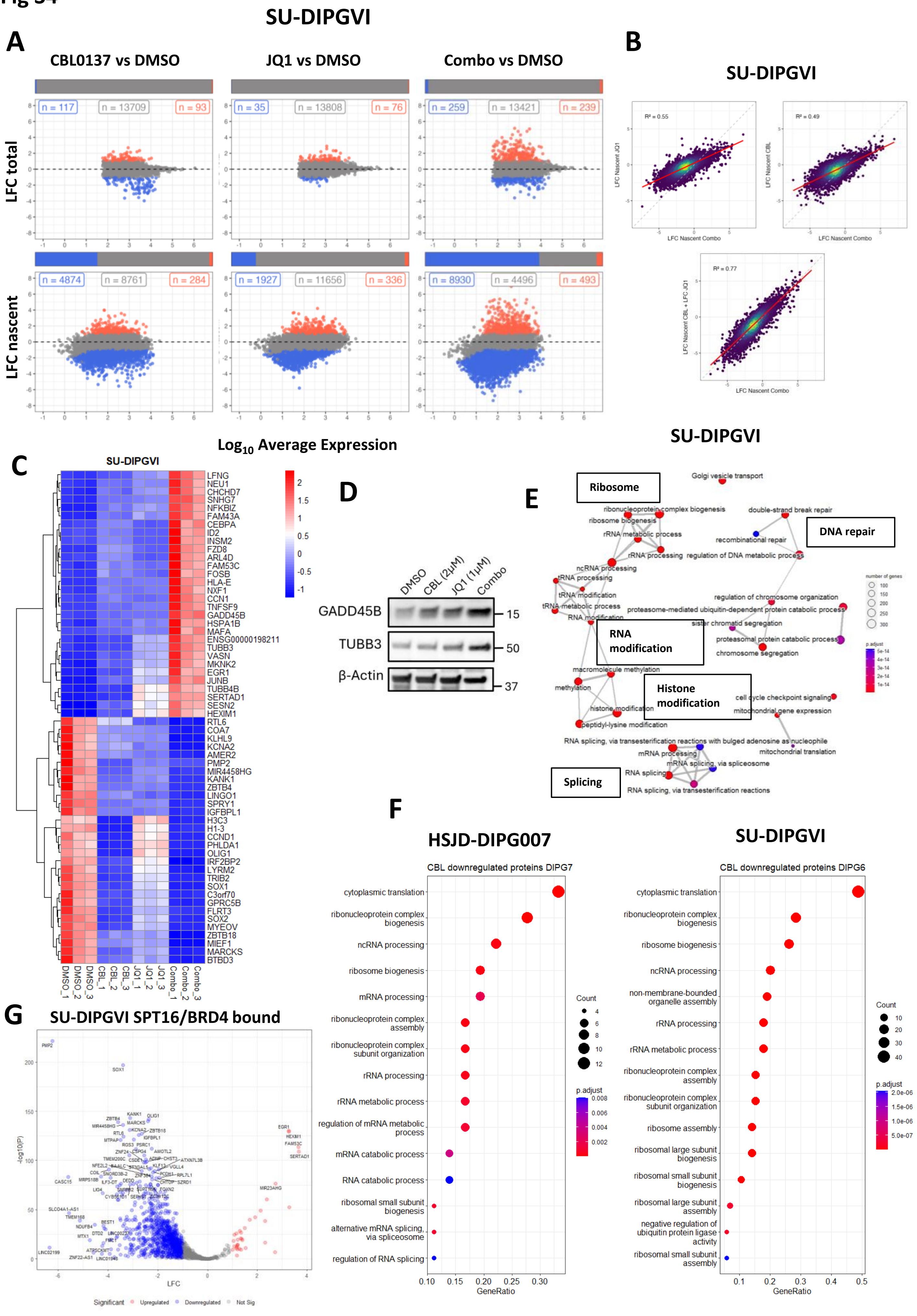
SU-DIPGVI
A
CBL0137 vs DMSO
JQ1 vs DMSO
Combo vs DMSO
LFC total
n = 117
n = 13709
n = 93
n = 35
n = 13808
n = 76
n = 259
n = 13421
n = 239
LFC nascent
n = 4874
n = 8761
n = 284
n = 1927
n = 11656
n = 336
n = 8930
n = 4496
n = 493
Log10 Average Expression
B
SU-DIPGVI
R² = 0.55
R² = 0.49
R² = 0.77
LFC Nascent JQ1
LFC Nascent CBL
LFC Nascent CBL + LFC JQ1
LFC Nascent Combo
C
SU-DIPGVI
LFNG
NEU1
CHCHD7
SNHG7
NFKBIZ
FAM43A
CEBPA
ID2
INSM2
FZD8
ARL4D
FAM53C
FOSB
HLA-E
NXF1
CCN1
TNFSF9
GADD45B
HSPA1B
MAFA
ENSG00000198211
TUBB3
VASN
MKNK2
EGR1
JUNB
TUBB4B
SERTAD1
SESN2
HEXIM1
RTL6
COA7
KLHL9
KCNA2
AMER2
PMP2
MIR4458HG
KANK1
ZBTB4
LINGO1
SPRY1
IGFBPL1
H3C3
H1-3
CCND1
PHLDA1
OLIG1
IRF2BP2
LYRM2
TRIB2
SOX1
C3orf70
GPRC5B
FLRT3
SOX2
MYEOV
ZBTB18
MIEF1
MARCKS
BTBD3
DMSO_1
DMSO_2
DMSO_3
CBL_1
CBL_2
CBL_3
JQ1_1
JQ1_2
JQ1_3
Combo_1
Combo_2
Combo_3
D
DMSO
CBL (2μM)
JQ1 (1μM)
Combo
GADD45B
TUBB3
β-Actin
15
50
37
E
SU-DIPGVI
Ribosome
DNA repair
RNA modification
Histone modification
Splicing
Golgi vesicle transport
ribonucleoprotein complex biogenesis
ribosome biogenesis
rRNA metabolic process
rRNA processing
ncRNA processing
tRNA processing
tRNA modification
tRNA metabolic process
RNA modification
double-strand break repair
recombinational repair
regulation of DNA metabolic process
regulation of chromosome organization
proteasome-mediated ubiquitin-dependent protein catabolic process
sister chromatid segregation
proteasomal protein catabolic process
chromosome segregation
macromolecule methylation
methylation
histone modification
peptidyl-lysine modification
cell cycle checkpoint signaling
mitochondrial gene expression
RNA splicing, via transesterification reactions with bulged adenosine as nucleophile
mitochondrial translation
mRNA processing
mRNA splicing, via spliceosome
RNA splicing
RNA splicing, via transesterification reactions
number of genes
p.adjust
F
HSJD-DIPG007
CBL downregulated proteins DIPG7
cytoplasmic translation
ribonucleoprotein complex biogenesis
ncRNA processing
ribosome biogenesis
mRNA processing
ribonucleoprotein complex assembly
ribonucleoprotein complex subunit organization
rRNA processing
rRNA metabolic process
regulation of mRNA metabolic process
mRNA catabolic process
RNA catabolic process
ribosomal small subunit biogenesis
alternative mRNA splicing, via spliceosome
regulation of RNA splicing
Count
p.adjust
GeneRatio
SU-DIPGVI
CBL downregulated proteins DIPG6
cytoplasmic translation
ribonucleoprotein complex biogenesis
ribosome biogenesis
ncRNA processing
non-membrane-bounded organelle assembly
rRNA processing
rRNA metabolic process
ribonucleoprotein complex assembly
ribonucleoprotein complex subunit organization
ribosome assembly
ribosomal large subunit biogenesis
ribosomal small subunit biogenesis
ribosomal large subunit assembly
negative regulation of ubiquitin protein ligase activity
ribosomal small subunit assembly
Count
p.adjust
GeneRatio
G
SU-DIPGVI SPT16/BRD4 bound
-log10(P)
LFC
Significant
Upregulated
Downregulated
Not Sig

**Figure S4. CBL0137 and JQ1 induce a broad suppression of transcription. Related to Fig 4.**
**A)** Total RNA (upper) and nascent RNA (lower) MA plots showing significantly differentially expressed genes in SU-DIPGVI cells treated with CBL0137 (2µM), JQ1 (1µM), or the combination for 4 h. Blue points represent significantly (FDR < 0.05, LFC < -1) downregulated genes, red points represent significantly upregulated genes (FDR < 0.05, LFC >1) with the number of genes indicated. Bars indicate the proportion of DE genes. **B**) Correlations of log fold-change in nascently transcribed genes between combination therapy, each monotherapy, and the sum of both monotherapies in SU-DIPGVI cells. **C)** Nascent RNA-seq heatmap displaying top 30 up and downregulated genes DMSO vs Combination treated SU-DIPGVI cells. **D**) GADD45B and TUBB3 western blot from treated SU-DIPGVI cells after 24 h. representative blot from 3 independent experiments. **E**) Enrichment map of over-represented gene ontologies among nascently transcribed genes downregulated by combination treatment in SU-DIPGVI cells, with high-level processes highlighted in labelled boxes. **F**) Over-represented ontologies of significantly downregulated proteins identified by mass spectrometry in HSJD-DIPG007 (left) and SU-DIPGVI (right) cells treated with CBL0137 at IC50 doses. **G**) Volcano plot showing differential expression of nascently transcribed genes associated with SPT16 and BRD4 peaks in SU-DIPGVI cells.

Fig S5

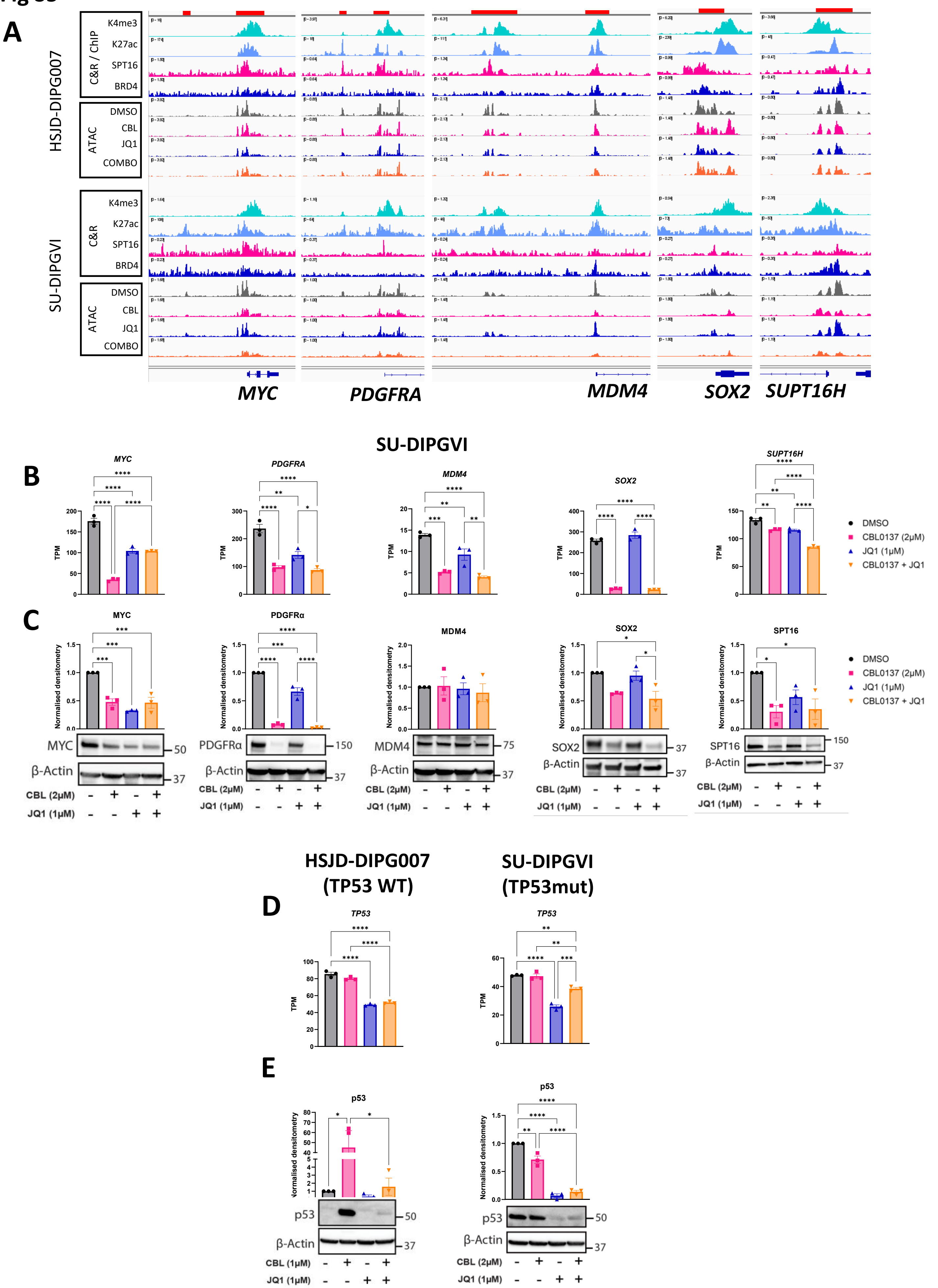

A
HSJD-DIPG007
SU-DIPGVI
C&R / ChIP
K4me3
K27ac
SPT16
BRD4
ATAC
DMSO
CBL
JQ1
COMBO
C&R
MYC
PDGFRA
MDM4
SOX2
SUPT16H
B
SU-DIPGVI
TPM
DMSO
CBL0137 (2µM)
JQ1 (1µM)
CBL0137 + JQ1
C
MYC
PDGFRα
MDM4
SOX2
SPT16
Normalised densitometry
β-Actin
CBL (2µM)
JQ1 (1µM)
D
HSJD-DIPG007 (TP53 WT)
SU-DIPGVI (TP53mut)
TP53
E
p53
CBL (1µM)

**Fig S5 cont.**

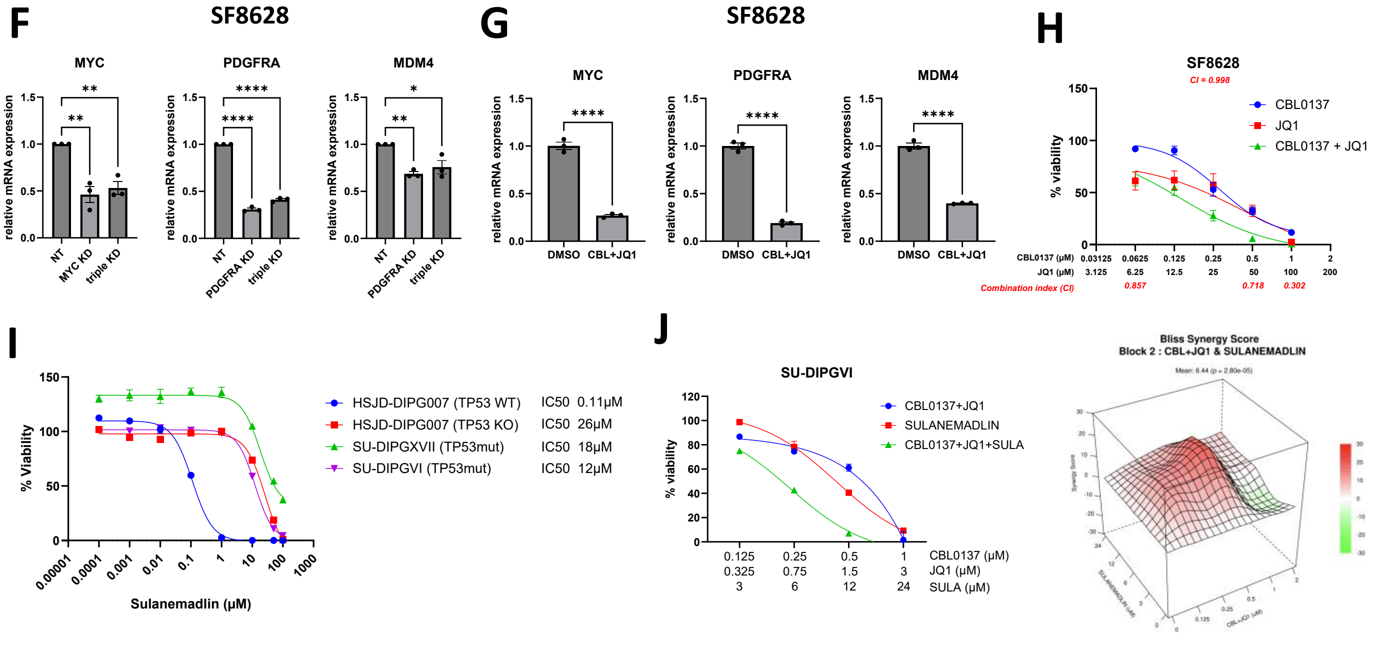


**Figure S5. Chromatin compaction by CBL0137 and JQ1 decreases expression of oncogenes. Related to Figure 5.**

**A)** Gene tracks displaying H3K4me3, H3K27ac [41, 47], IgG, SPT16, BRD4 CUT&RUN, and ATAC-seq signal across the 4 indicated treatments in HSJD-DIPG007 and SU-DIPGVI cells at the *MYC*, *PDGFRA*, *MDM4, SOX2* and *SUPT16H* loci. **B**) RNA-seq TPM values (mean +/- SEM; n=3) for *MYC, PDGFRA, MDM4, SOX2,* and *SUPT16H* in SU-DIPGVI cells treated for 4 h with CBL0137 and JQ1 **C**) Representative MYC, PDGFRα, MDM4, SOX2 and SPT16 western blots and bar charts displaying densitometry (mean +/- SEM; n=3) normalized to β-Actin and expressed as a fold-change relative to the DMSO control. 24 h treatment. PDGFRα and MDM4 were run on the same gel, and use the same loading control. SOX2 was run on same gel as Fig S4D and use the same loading control. **D**) RNA-seq TPM values (mean +/- SEM; n=3) for TP53 in the indicated cell lines. 4h treatment. **E**) Representative p53 western blots and bar charts displaying densitometry (mean +/- SEM; n=3) normalized to β-Actin and expressed as a fold-change relative to the DMSO control. 24 h treatment. **F**) qPCR bar plots illustrating gene knockdown of MYC, PDGFRA, and MDM4 in SF8628 cells. Cells were transfected with siRNA targeting each gene individually or in combination (triple knockdown) for 24 hours. **G**) qPCR bar plots illustrating relative expression of MYC, PDGFRA and MDM4 in SF8628 cells treated with DMSO or the combination of CBL0137 (0.6 μM) and JQ1 (1 μM) for 24 h. **H**) cytotoxicity assay in SF8628 DMG cells treated with CBL0137 and JQ1. Combination index values were calculated with Calcusyn. **I**) Sulanemadlin toxicity assay in HSJD-DIPG007 cells with and without TP53, SU-DIPGVI (p53 mutant), and SU-DIPGXVII (p53 mutant) cells. **J**) Cytotoxicity assay comparing CBL0137+JQ1, sulanemadlin, and the combination of all 3 drugs in SU-DIPGVI cells after 96 h and Bliss synergy score matrix comparing CBL0137+JQ1 and sulanemadlin. **(B-F)** Significance was calculated using a one-way ANOVA with Sidak's multiple comparisons test for single and combination treatments. For **G)** Statistical significance was determined using an unpaired student's t-test. * p<0.05, **p<0.01, ***p<0.001, ****p<0.0001.

**Fig S6**

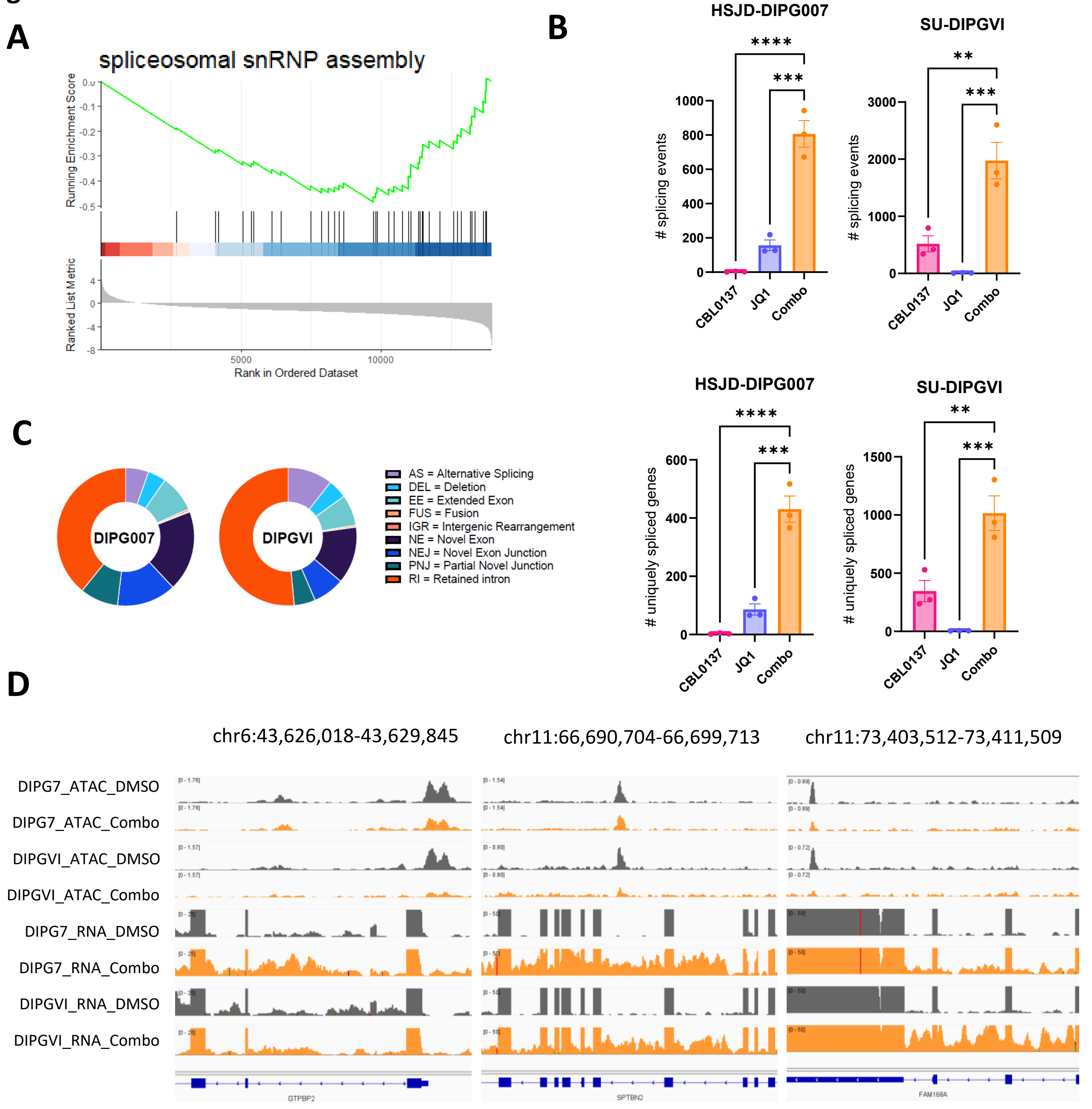


**Figure S6. Combination treatment leads to alternative splicing in DMG cells**

**A)** GSEA enrichment plot showing negative enrichment (downregulation) of the spliceosomal snRNP assembly pathway in SU-DIPGVI cells (DMSO vs Combination). **B-C**) Bar graph displaying the total number of splicing events (upper) or uniquely spliced genes (lower) in HSJD-DIPG007 and SU-DIPGVI cells with the indicated treatments. Significance was calculated using a one-way ANOVA with Dunnet's multiple comparisons test comparing single and combination treatments. $^{**}p<0.01$, $^{***}p<0.001$, $^{****}p<0.0001$. **C**) Pie charts depicting the breakdown of differential splicing events common to 3 replicates for the combination vs DMSO condition, in each model. **D**) Gene tracks display ATAC-seq signal (upper tracks) and RNA-seq coverage (lower tracks) in DMSO control and combination treated HSJD-DIPG007 and SU-DIPGVI cells. Representative tracks for n=2-3 experiments for each model.

### Fig S7

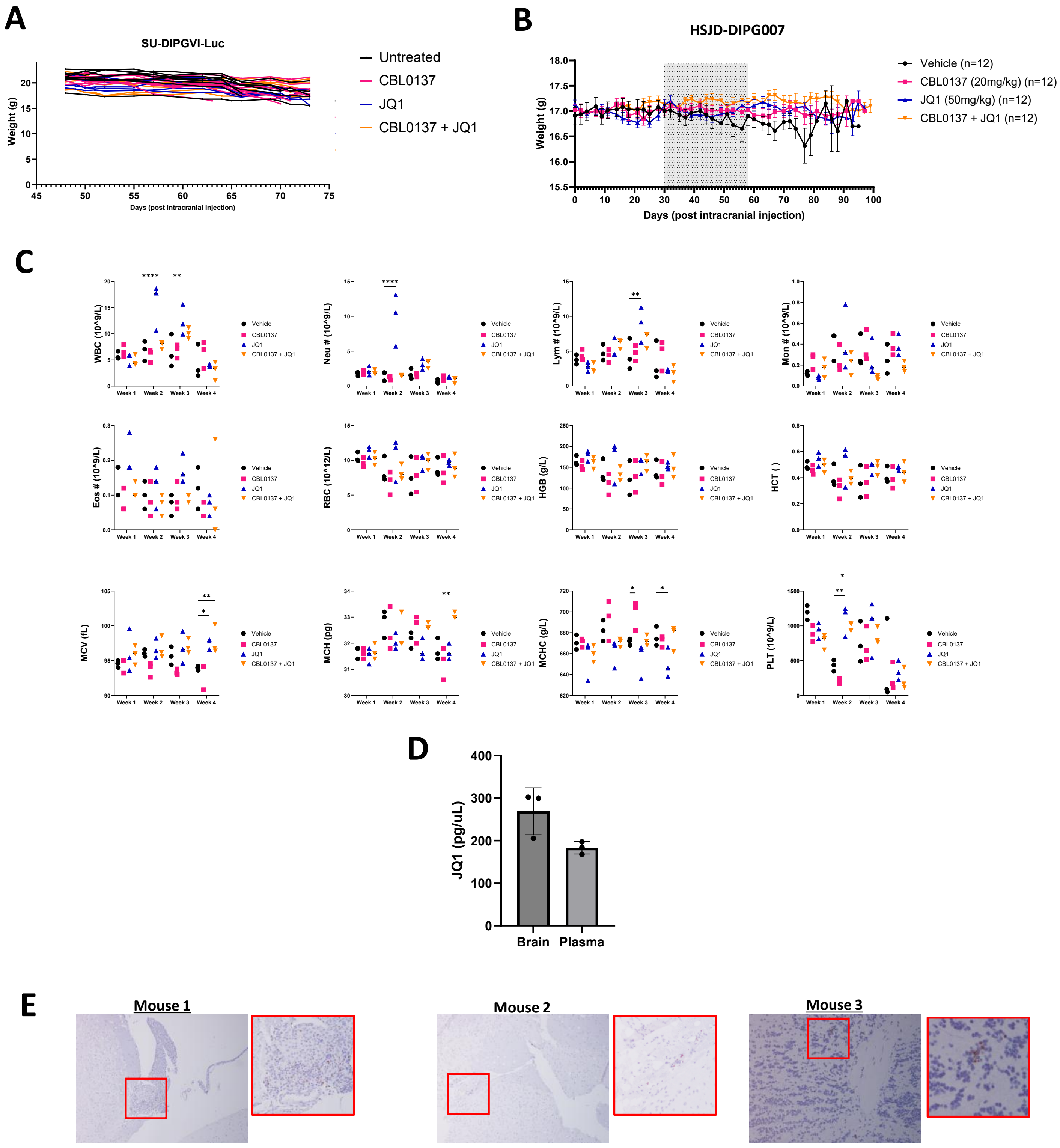
A
SU-DIPGVI-Luc
Weight (g)
Days (post intracranial injection)
Untreated
CBL0137
JQ1
CBL0137 + JQ1
B
HSJD-DIPG007
Weight (g)
Days (post intracranial injection)
Vehicle (n=12)
CBL0137 (20mg/kg) (n=12)
JQ1 (50mg/kg) (n=12)
CBL0137 + JQ1 (n=12)
C
WBC (10^9/L)
Neu # (10^9/L)
Lym # (10^9/L)
Mon # (10^9/L)
Eos # (10^9/L)
RBC (10^12/L)
HGB (g/L)
HCT ( )
MCV (fL)
MCH (pg)
MCHC (g/L)
PLT (10^9/L)
Week 1
Week 2
Week 3
Week 4
Vehicle
CBL0137
JQ1
CBL0137 + JQ1
D
JQ1 (pg/uL)
Brain
Plasma
E
Mouse 1
Mouse 2
Mouse 3

#### Fig S7 continued

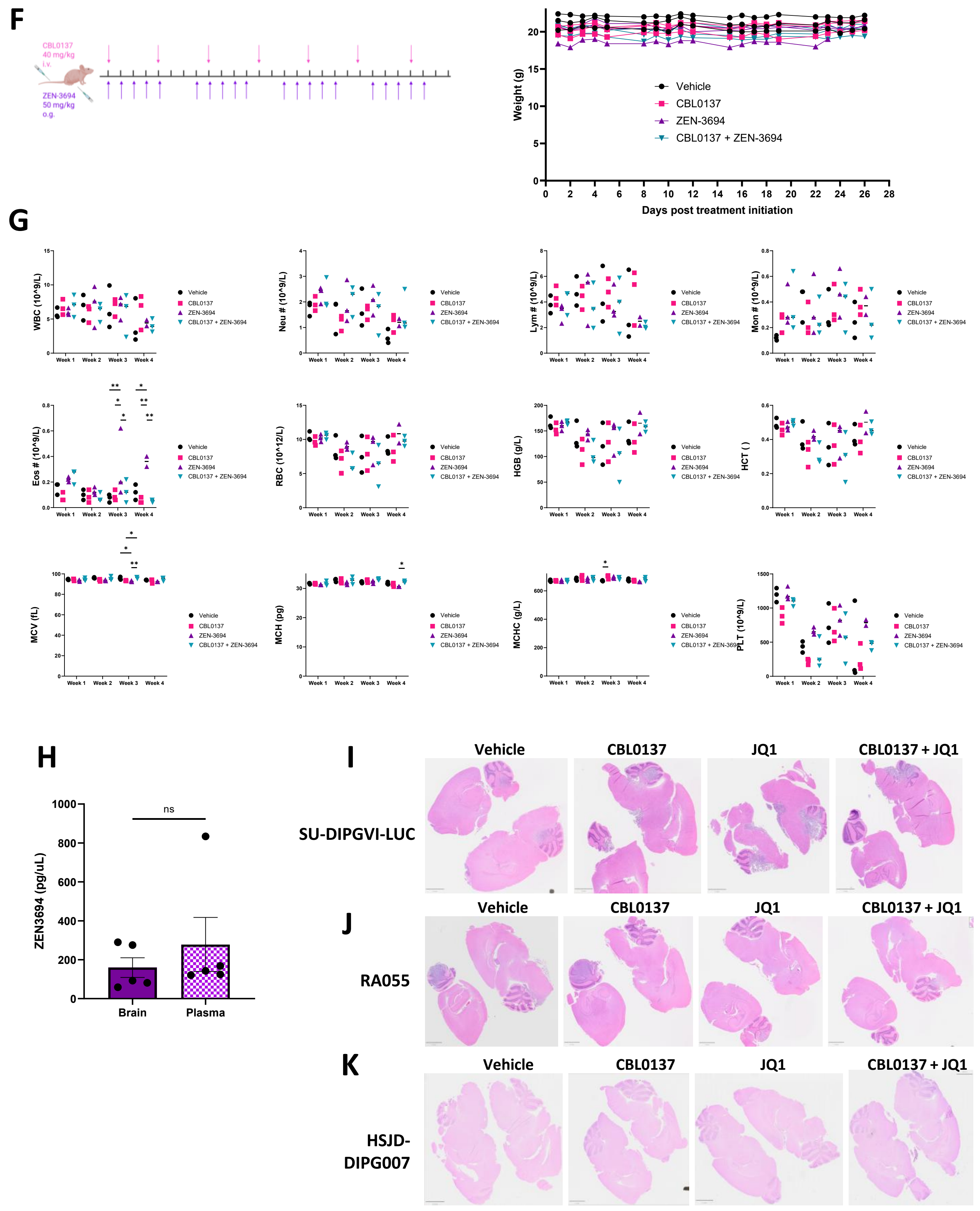

**Figure S7. Therapeutic efficacy of CBL0137 and JQ1 combination treatment in orthotopic models of DMG related to Figure 6.**

**A-B**) Weights of mice before, during and after treatment period for the SU-DIPGVI-Luc (**A)** and HSJD-DIPG007 model (**B**). Data for individual mice is shown in (**A**) and averaged mice are shown in (**B**). (**C**) Hematologic measurements from non-tumor-bearing mice. A two-way ANOVA with Dunnett's multiple comparisons test was used to assess significance between vehicle and treatment groups (n = 3 per cohort). *p < 0.05, **p < 0.01, ***p < 0.001, ****p < 0.0001.(**D**) Pharmacokinetic analysis of JQ1 in brain and plasma following 4 weeks. **(E**) IHC staining for Ki67 on brain samples from endpoint surviving mice in RA055 model demonstrating positive engraftment. **(F)** Treatment scheme for CBL0137 and ZEN-3694 administration, and weights of mice before, during and after treatment period. **(G)** Hematologic measurements from non-tumor-bearing mice. A two-way ANOVA with Dunnett's multiple comparisons test was used to assess significance between vehicle and treatment groups (n = 3 per cohort). *p < 0.05, **p < 0.01, ***p < 0.001, ****p < 0.0001. **(H)** Pharmacokinetic analysis of ZEN-3694 in brain and plasma following 4 weeks **(I-K)** Whole brain scans of H&E stained brains from the SU-DIPGVI-LUC model (**I**), RA055 model **(J**) and HSJD-DIPG007 model (**K**). Representative images shown. Scale = 2000µM.

Fig S8

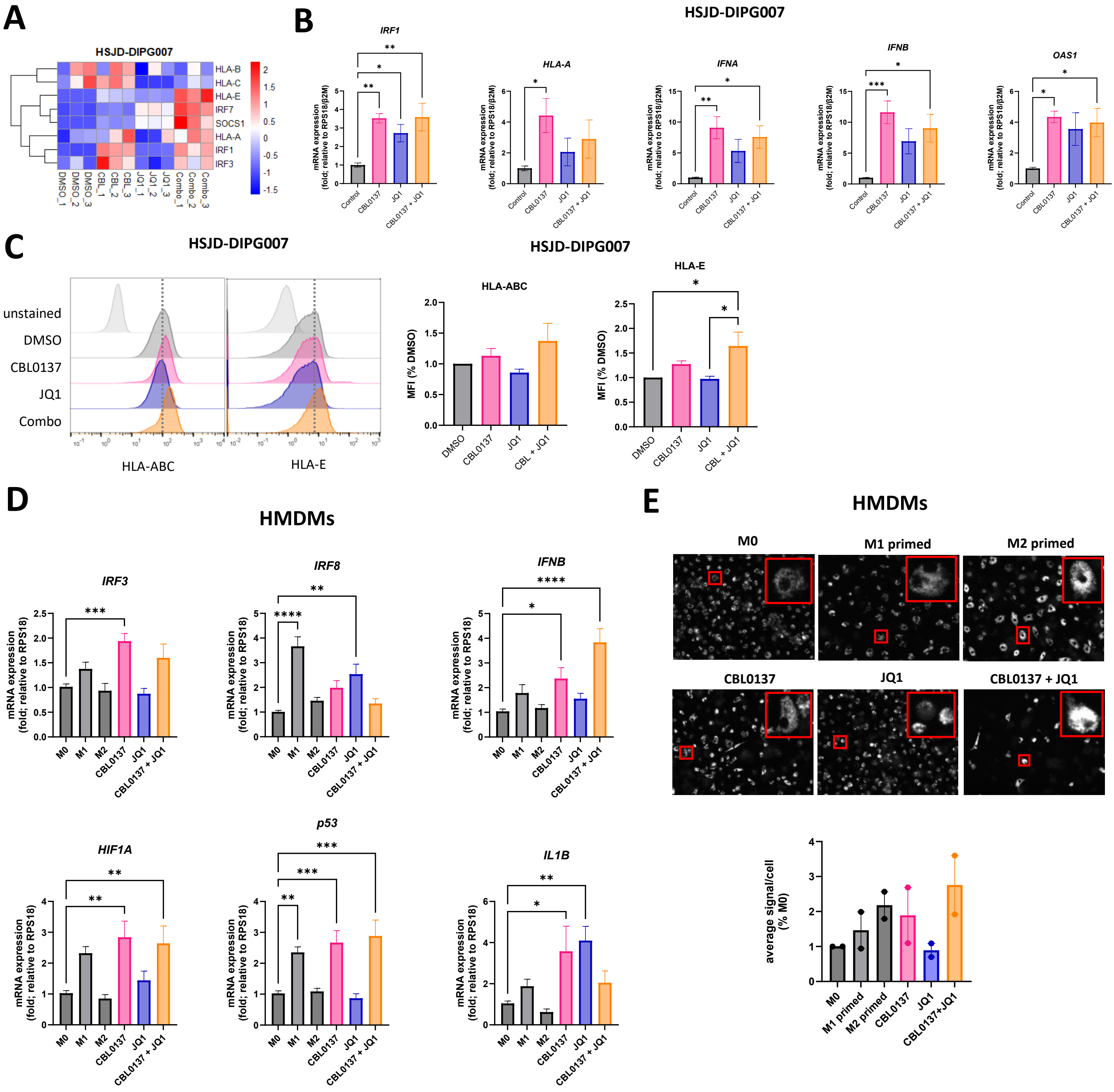

A
HSJD-DIPG007
HLA-B
HLA-C
HLA-E
IRF7
SOCS1
HLA-A
IRF1
IRF3
DMSO_1
DMSO_2
DMSO_3
CBL_1
CBL_2
CBL_3
JQ1_1
JQ1_2
JQ1_3
Combo_1
Combo_2
Combo_3
B
HSJD-DIPG007
IRF1
HLA-A
IFNA
IFNB
OAS1
mRNA expression (fold; relative to RPS18/β2M)
Control
CBL0137
JQ1
CBL0137 + JQ1
C
HSJD-DIPG007
unstained
DMSO
CBL0137
JQ1
Combo
HLA-ABC
HLA-E
HSJD-DIPG007
HLA-ABC
HLA-E
MFI (% DMSO)
DMSO
CBL0137
JQ1
CBL + JQ1
D
HMDMs
IRF3
IRF8
IFNB
HIF1A
p53
IL1B
mRNA expression (fold; relative to RPS18)
M0
M1
M2
CBL0137
JQ1
CBL0137 + JQ1
E
HMDMs
M0
M1 primed
M2 primed
CBL0137
JQ1
CBL0137 + JQ1
average signal/cell (% M0)
M0
M1 primed
M2 primed
CBL0137
JQ1
CBL0137+JQ1

**Fig S8 cont.**

F

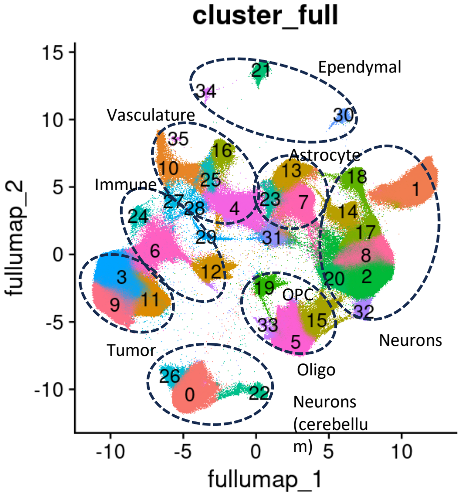


G

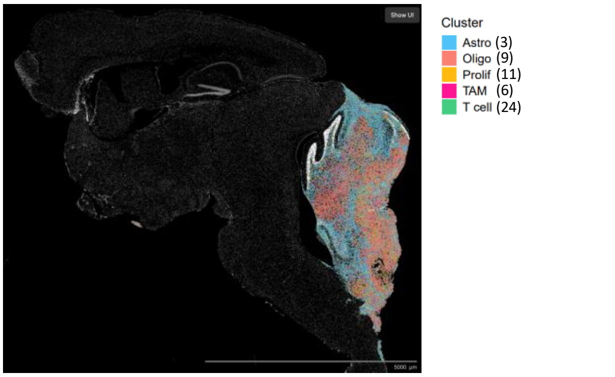


H

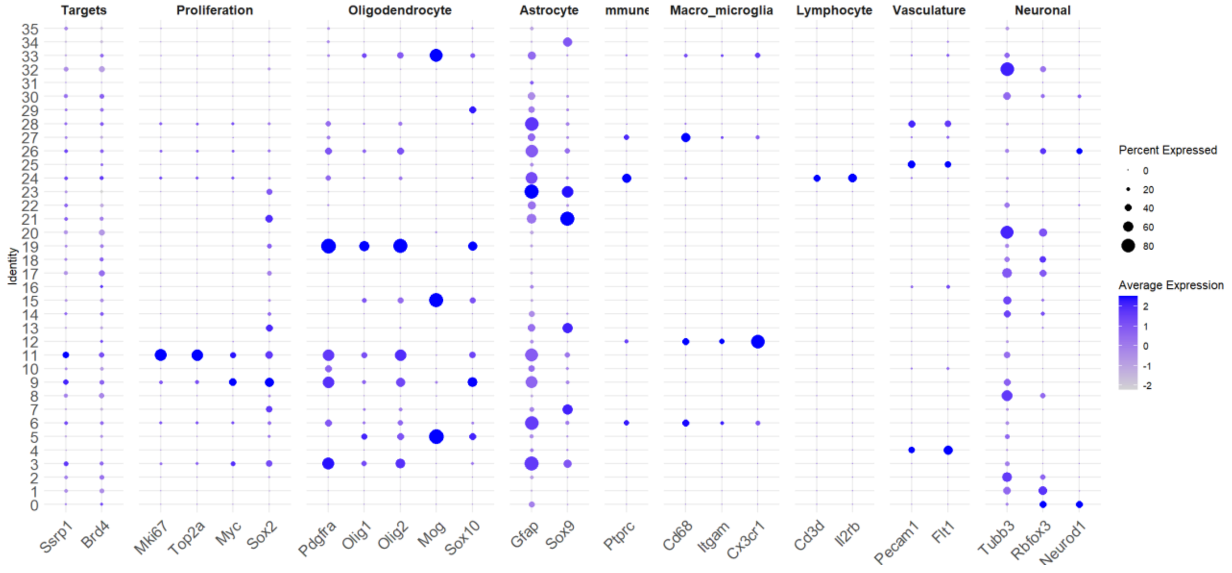


I

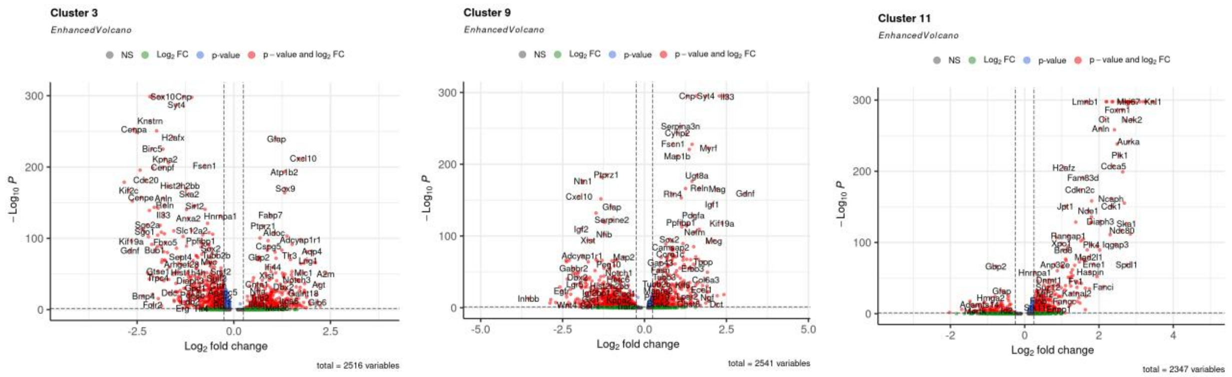


J

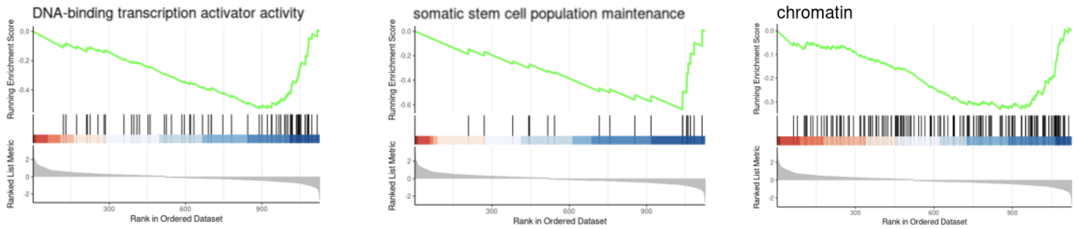


K

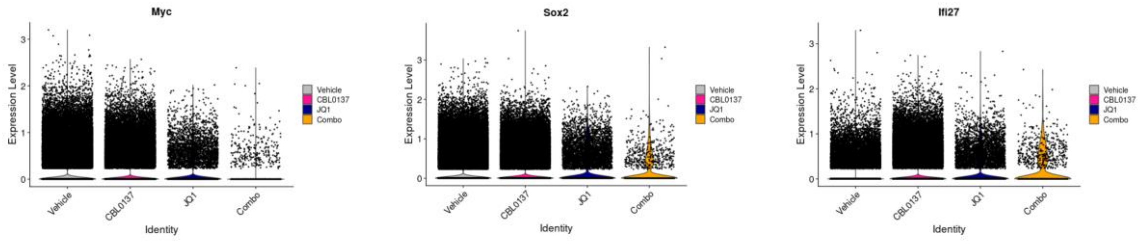

**Figure S8. CBL0137 and JQ1 influence immune-associated pathways in DMG and tumor-infiltrating immune cells, related to Figure 7.**

**A)** Nascent RNA-seq heatmap displaying interferon and antigen presentation genes in the indicated treatment groups for HSJD-DIPG007 cells. **B**) qPCR analysis of interferon and inflammatory genes in HSJD-DIPG007 cells treated for 48h with CBL0137 (0.5μM) or JQ1 (1μM). (n=3). **C)** Flow cytometry histograms for HLA-ABC and HLA-E in treated SU-DIPG007 cells (24h). Representative data from at least 3 independent experiments. Median Fluorescence Intensity (MFI) values are shown. **D**) qPCR analysis of human monocyte-derived macrophage (HMDM) expression M1 proinflammatory genes following treatment with 1μM CBL0137 or 1μM JQ1, alone or in combination for 48 h, compared to untreated macrophages (M0) and macrophages exposed to M1 and M2 priming conditions. N=3. **E**) Fluorescent bead-uptake phagocytosis assay of HMDMs in M1 or M2 priming conditions or treated with CBL0137 and JQ1 for 48 hours quantification of fluorescent signal/cell calculated from 2 wells. P-values were calculated using a one-way ANOVA test with Dunnett's multiple comparison test. Data is presented as mean values ± SEM. *p<0.05, **p<0.01, ***p<0.001, ****p<0.0001. **F**) Xenium single cell UMAP projection of 4 integrated samples. Major clusters are annotated based on lineage markers. **G**) Vehicle treated brain slice stained with DAPI (white) with tumor (clusters 3, 9, 11) and immune (clusters 6, 24) associated clusters overlaid. **H**) Dot plot displaying marker gene expression across the 35 UMAP clusters. **I**) Volcano plots displaying significantly differentially expressed genes in each of the 3 tumor clusters compared to the other 2 tumor clusters. **J)** GSEA enrichment plots comparing combination to vehicle (red = upregulated, blue = downregulated) in tumor cells (clusters 3+9+11). **K**) Violin plots for select genes (*Myc, Sox2,* and *Ifi27)* significantly differentially expressed genes in the vehicle vs combination treated tumor cells.

**Supplementary Table 1 (Excel): Gene lists from CUT&RUN, ATAC-seq, RNA-seq, proteomics, and spatial in situ experiments.** (All lists are ranked by FDR significance.)

**A)** Chromosome coordinates for SPT16 and BRD4 MACS2 peaks from merged CUT&RUN replicates in HSJD-DIPG007, SU-DIPGVI, and VUMC-DIPG10 DMG cells. **B)** Genes associated with SPT16 and BRD4 peaks in HSJD-DIPG007, SU-DIPGVI, and VUMC-DIPG10 DMG cells. **C)** Genes linked to differentially accessible regions (DARs) identified by ATAC-seq CSAW analysis ($FDR < 0.05$, $LFC > 1$ = opened; $LFC < -1$ = closed) in HSJD-DIPG007 and SU-DIPGVI cells treated with CBL0137 (1 μM for HSJD-DIPG007, 2 μM for SU-DIPGVI) or JQ1 (1 μM) for 4 h. **D)** Total and nascent RNA-seq differentially expressed genes ($FDR < 0.05$, $|LFC| > 1$) in HSJD-DIPG007 and SU-DIPGVI cells treated with CBL0137 or JQ1 for 2 h. **E)** RNA-seq differentially expressed genes ($FDR < 0.05$, $|LFC| > 1$) in HSJD-DIPG007 and SU-DIPGVI cells treated with CBL0137 or JQ1 for 4 h. **F)** Genes with differential splicing events common across three independent replicates in HSJD-DIPG007 and SU-DIPGVI cells treated with CBL0137 and JQ1 (4 h). **G)** Differentially expressed proteins identified by proteomics in HSJD-DIPG007 and SU-DIPGVI cells treated with CBL0137 (0.4 μM and 0.9 μM, respectively) for 24 h. **H)** Xenium spatial transcriptomics marker genes differentially expressed ($FDR < 0.05$, $LFC > 1$) in each cluster compared to all clusters. **I)** Xenium marker genes differentially expressed between tumor clusters ($FDR < 0.05$, $LFC > 1$). **J)** Xenium differentially expressed genes comparing combination therapy versus vehicle in tumor, TAM, and T cell clusters ($FDR < 0.05$, $|LFC| > 0.5$).

**Supplementary Table 2.** $IC_{50}$ values for panel of DMG cells treated with JQ1 from cytotoxicity assays.

| JQ1 | |
|---|---|
| **Cell Line** | **IC50 (µM)** |
| HSJD-DIPG007 | 18.87 |
| SU-DIPGVI | 2.326 |
| RA055 | 3.142 |
| VUMC-DIPG10 | 5.929 |
| P003302 | 3.142 |
| AUS-DIPG017 | 47.10 |
| P002306 | 8.702 |

**Supplementary Table 3.** $IC_{50}$ values for panel of DMG cells treated with Trotabresib from cytotoxicity assays.

| Trotabresib | |
|---|---|
| **Cell Line** | **IC50 (µM)** |
| HSJD-DIPG007 | 6.780 |
| SU-DIPGVI | 2.535 |
| RA055 | 0.6552 |
| SU-DIPGXII_K27M | 5.9875 |
| SU-DIPGXII_KO | 24.6025 |
| VUMC-DIPG10 | 1.947 |

**Supplementary Table 4.** $IC_{50}$ values for DMG isogenic cell lines, with or without the H3K27M mutation, treated with CBL0137, JQ1 or Trotabresib from cytotoxicity assays.

| | CBL0137 | | JQ1 | | Trotabresib | |
|---|---|---|---|---|---|---|
| | **K27M** | **KO** | **K27M** | **KO** | **K27M** | **KO** |
| **IC50 (µM)** | 1.001 | 5.123 | 8.08 | 43.123 | 5.988 | 24.603 |

**Supplementary Table 5.** Key mutations in DMG/HGG cell lines used in this study

| Cell Line | Known mutations |
|---|---|
| HSJD-DIPG007 | H3.3K27M, TP53wt, ACVR1mut, PPM1Dmut, PIK3CAmut, MYCamp |
| SU-DIPGVI | H3.3K27M, TP53mut |
| RA055 | H3.3K27M, TP53mut, PDGFRAmut, PDGFRA amp |
| SU-DIPGXIII | H3.3K27M, TP53mut |
| VUMC-DIPG10 | H3 WT, TP53mut, MYCNamp |
| P003302 | H3.1K27M, TP53mut, Chr17p loss, ERBB4gain, ACVR1mut, ACVR1gain |
| P002306 | H3.3K27M, ACVR1, PPM1Dmut, PIK3CAmut, ACVR1mut |
| AUS-DIPG17 | H3 WT, TP53mut, CDKN2A/B loss |
| P005401 | H3.1K27M, p53mut, EGFRdel |
| P016802 | H3 WT, TP53mut, MYCNamp, HMGA1 amp, EGFR amp |
| SF8628 | H3.3K27M |
| SU-DIPGXVII | H3.3K27M, TP53mut |
| HSJD-DIPG11 | H3.3K27M |
| HSJD-DIPG12 | H3.3K27M, TP53mut |
| HSJD-DIPG13 | H3.3K27M, TP53mut |

**Supplementary Table 6:** Multiple comparisons test for SU-DIPGVI-Luc orthotopic animal model

| SU-DIPGVI-Luc | P-value | Q value | Discovery |
|---|---|---|---|
| Vehicle vs JQ1 | 0.0141 | 0.0036 | Yes |
| Vehicle vs CBL0137 + JQ1 | 0.0010 | 0.0010 | Yes |
| CBL0137 vs CBL0137 + JQ1 | 0.0021 | 0.0011 | Yes |
| JQ1 vs CBL0137 + JQ1 | 0.0071 | 0.0024 | Yes |

**Supplementary Table 7:** Multiple comparisons test for HSJD-DIPG007 orthotopic animal model

| HSJD-DIPG007 | P-value | Q value | Discovery |
|---|---|---|---|
| Vehicle vs CBL0137 | 0.0044 | 0.0059 | Yes |
| Vehicle vs JQ1 | 0.0007 | 0.0014 | Yes |
| Vehicle vs CBL0137 + JQ1 | 0.0001 | 0.0004 | Yes |
| CBL0137 vs CBL0137 + JQ1 | 0.0095 | 0.0096 | Yes |

**Supplementary Table 8:** Multiple comparisons test for RA055 orthotopic animal model

| RA055 | P-value | Q value | Discovery |
|---|---|---|---|
| Vehicle vs CBL0137 | 0.00640 | 0.00162 | Yes |
| Vehicle vs JQ1 | 0.00030 | 0.00015 | Yes |
| Vehicle vs CBL0137 + JQ1 | 0.00010 | 0.00010 | Yes |
| CBL0137 vs CBL0137 + JQ1 | 0.00370 | 0.00125 | Yes |
| JQ1 vs CBL0137 + JQ1 | 0.02480 | 0.00501 | Yes |

**Supplementary Table 9:** Biochemical parameters in CBL0137/JQ1 treated mice

| Biochemical Parameter | Two weeks post treatment | | | | Four weeks post treatment | | | |
|---|---|---|---|---|---|---|---|---|
| | Vehicle | CBL0137 | JQ1 | CBL0137 + JQ1 | Vehicle | CBL0137 | JQ1 | CBL0137 + JQ1 |
| Albumin (g/L) | 41 | 25 | 22 | 37 | 40 | 44 | 39 | 43 |
| Alkaline Phosphatase (U/L) | 80 | 208 | 52 | 129 | 76 | 58 | 70 | 64 |
| Alanine Aminotransferase (U/L) | 19 | 21 | 20 | 20 | 44 | 39 | 40 | 51 |
| Amylase (U/L) | 915 | 546 | 282 | 535 | 1130 | 1281 | 3377 | 513 |
| Total Bilirubin (umol/L) | 6 | 5 | 10.26 | 6 | 5 | - | 5 | - |
| Blood Urea Nitrogen (mmol/L) | 5.9 | 5 | 5 | 5.9 | 7.3 | 7.86 | 6.4 | 6.6 |
| Calcium (mmol/L) | 2.52 | 1.71 | <2 | 2.5 | 3.03 | 3.27 | 3.42 | 3.29 |
| Phosphorus (mmol/L) | 2.19 | 5.36 | - | 2.14 | 4.15 | 4.84 | 4.2 | 4.48 |
| Creatinine (umol/L) | 21 | <18 | <18 | <18 | <18 | <18 | <18 | <18 |
| Glucose (mmol/L) | 7.3 | 4.4 | 6 | 9.3 | 14.5 | 14.2 | 18.4 | 13.8 |
| Sodium (mmol/L) | 151 | 153 | 150 | 146 | 158 | 155 | 151 | 158 |
| Potassium (mmol/L) | 7.4 | 7.4 | 5.6 | 8.1 | >8.5 | - | >8.5 | - |
| Total Protein (g/L) | 55 | 35 | <40 | 54 | 52 | 55 | 55 | 57 |
| Globulin (g/L) | 14 | 10 | - | 16 | 12 | 12 | 16 | 14 |

**Supplementary Table 10:** Biochemical parameters in CBL0137 + ZEN-3694 treated mice

| Biochemical Parameter | Two weeks post treatment | | | | Four weeks post treatment | | | |
|---|---|---|---|---|---|---|---|---|
| | Vehicle | CBL0137 | ZEN-3694 | CBL0137 + ZEN-3694 | Vehicle | CBL0137 | ZEN-3694 | CBL0137 + ZEN-3694 |
| Albumin (g/L) | 41 | 25 | 43 | 32 | 40 | 44 | 42 | 41 |
| Alkaline Phosphatase (U/L) | 80 | 208 | 73 | 95 | 76 | 58 | 71 | 75 |
| Alanine Aminotransferase (U/L) | 19 | 21 | 32 | 29 | 44 | 39 | 121 | 52 |
| Amylase (U/L) | 915 | 546 | 804 | 488 | 1130 | 1281 | 1059 | 881 |
| Total Bilirubin (umol/L) | 6 | 5 | 6.84 | 5.13 | 5 | - | 5.13 | 5.13 |
| Blood Urea Nitrogen (mmol/L) | 5.9 | 5 | 6.07 | 4.29 | 7.3 | 7.86 | 5.36 | 6.43 |
| Calcium (mmol/L) | 2.52 | 1.71 | 2.52 | 1.77 | 3.03 | 3.27 | 3.19 | 3.37 |
| Phosphorus (mmol/L) | 2.19 | 5.36 | 2.03 | 4.90 | 4.15 | 4.84 | 4.23 | 4.84 |
| Creatinine (umol/L) | 21 | <18 | <18 | <18 | <18 | <18 | <18 | <18 |
| Glucose (mmol/L) | 7.3 | 4.4 | 7.5 | 4.4 | 14.5 | 14.2 | 11.7 | 15.1 |
| Sodium (mmol/L) | 151 | 153 | 152 | 151 | 158 | 155 | 158 | 159 |
| Potassium (mmol/L) | 7.4 | 7.4 | 6.5 | 6.4 | >8.5 | - | >8.5 | >8.5 |
| Total Protein (g/L) | 55 | 35 | 56 | 41 | 52 | 55 | 54 | 52 |
| Globulin (g/L) | 14 | 10 | 11.4 | 9 | 12 | 12 | 12 | 11 |
